# Deep-learning-assisted Sort-Seq enables high-throughput profiling of gene expression characteristics with high precision

**DOI:** 10.1101/2022.11.23.517705

**Authors:** Huibao Feng, Fan Li, Tianmin Wang, Xin-hui Xing, An-ping Zeng, Chong Zhang

## Abstract

As an essential physiological process, gene expression determines the function of each cell. However, owing to the complex nondeterministic and nonlinear nature of gene expression, the steady-state intracellular protein abundance of a clonal population forms a distribution. The characteristics of this distribution, including expression strength and noise, are closely related to cellular behavior. Therefore, quantitative description of these characteristics is an important goal in biology. This task, however, has so far relied on arrayed methods, which are time-consuming and labor-intensive. To address this issue, we propose a deep-learning-assisted Sort-Seq approach (dSort-Seq) in this work, enabling high-throughput profiling of expression properties with high precision. We demonstrated the validity of dSort-Seq for large-scale assaying of the dose–response relationships of biosensors. In addition, we comprehensively investigated the contribution of transcription and translation to noise production in *E. coli*, from which we discovered that the expression noise is strongly coupled with the mean expression level instead of translation strength, even in the case of weak transcription. We also discovered that the transcriptional interference caused by overlapping RpoD-binding sites contributes to noise production, which suggested the existence of a simple and feasible noise control strategy in *E. coli*. Overall, dSort-Seq is able to efficiently determine the strength-noise landscape, which has promising applications in studies related to gene expression.

## INTRODUCTION

Cells are sophisticated instruments driven by the central dogma that build varieties of lives. For each cell, gene expression is a vital process by which information from genes flows to RNA and then to proteins, determining the traits of the cell. However, gene expression is often stochastic, as it involves many random events requiring the participation of various low-copy-number chemical components^1–6^. In addition, this process can be chaotic due to the high complexity of the regulatory network^7,8^. As a result, phenotypic heterogeneity exists among genetically identical cells even under the same environmental conditions^1^. Therefore, steady-state protein production in a clonal population exhibits a distribution, wherein the mean of the distribution (Mean) indicates the expression strength, and the squared coefficient of variation (CV^2^) exhibits the expression noise. These two characteristics are both important indicators that are closely related to the phenotypes of a population, such as the bioproduction efficiency^9,10^, drug resistance^11,12^ and antibiotic persistence^13^. To date, the quantitative description of expression strength and noise has been an important goal in biology to illustrate cellular behavior^14^. However, this task has relied on fluorescence microscopy^1,15^ and flow cytometry^14,16^ (FCM) assays of individual clonal populations, which are time-consuming and labor-intensive when testing large amounts of genetic variants. Therefore, a general, precise and high-throughput method for the profiling of expression properties is urgently needed.

To address the above issue, we focused on Sort-Seq^17–19^ (also named FlowSeq, FACS-seq), by which a library of cells with different expression intensities can be sorted into different bins and then quantified through next-generation sequencing (NGS) to derive the expression pattern of each genotype. This approach has been broadly used in profiling sequence-function relationships associated with transcriptional regulation^17,20–22^, translational regulation^17,18,23^, regulatory RNAs^24,25^, protein-sequence interactions^26^, etc. In addition, the validity of Sort-Seq has been demonstrated in a wide range of organisms, including bacteria, yeast and mammalian cells^19^. However, it remains difficult to derive precise expression characteristics from Sort-Seq data. Existing methods have focused on fitting the binned distribution to a log-normal^18,19,24,27^ or gamma distribution^20,21,28^, which are limited by the inexact representation capability of these probability densities^6,29,30^. On the other hand, the parameter learning process of these methods still needs to be improved. For instance, apart from the binned distribution, other data, such as the overall fluorescence intensity density, should be considered. Hence, to obtain expression properties with high throughput and high precision, a common, rigorous data processing method for Sort-Seq is needed.

Therefore, we have developed dSort-Seq, a deep-learning-assisted Sort-Seq approach (**Fig. 1**). In this method, instead of using log-normal or gamma distribution, we applied a two-component log-Gaussian mixture model (LGMM) to match the steady-state gene expression density, which is more precise and robust in fitting the real data. To decode Sort-Seq data, for the first time, we adopted a Bayesian neural network to perform parameter learning. These innovations significantly improve the accuracy of Sort-Seq to derive expression characteristics for thousands of variants. We demonstrated the validity of this pipeline from two aspects. First, dSort-Seq enables large-scale assays of dose–response relationships of biosensors with high precision, with which the optimal design can be efficiently identified. Second, it also supports the high-throughput exploration of noise production mechanisms.

**Figure 1.**
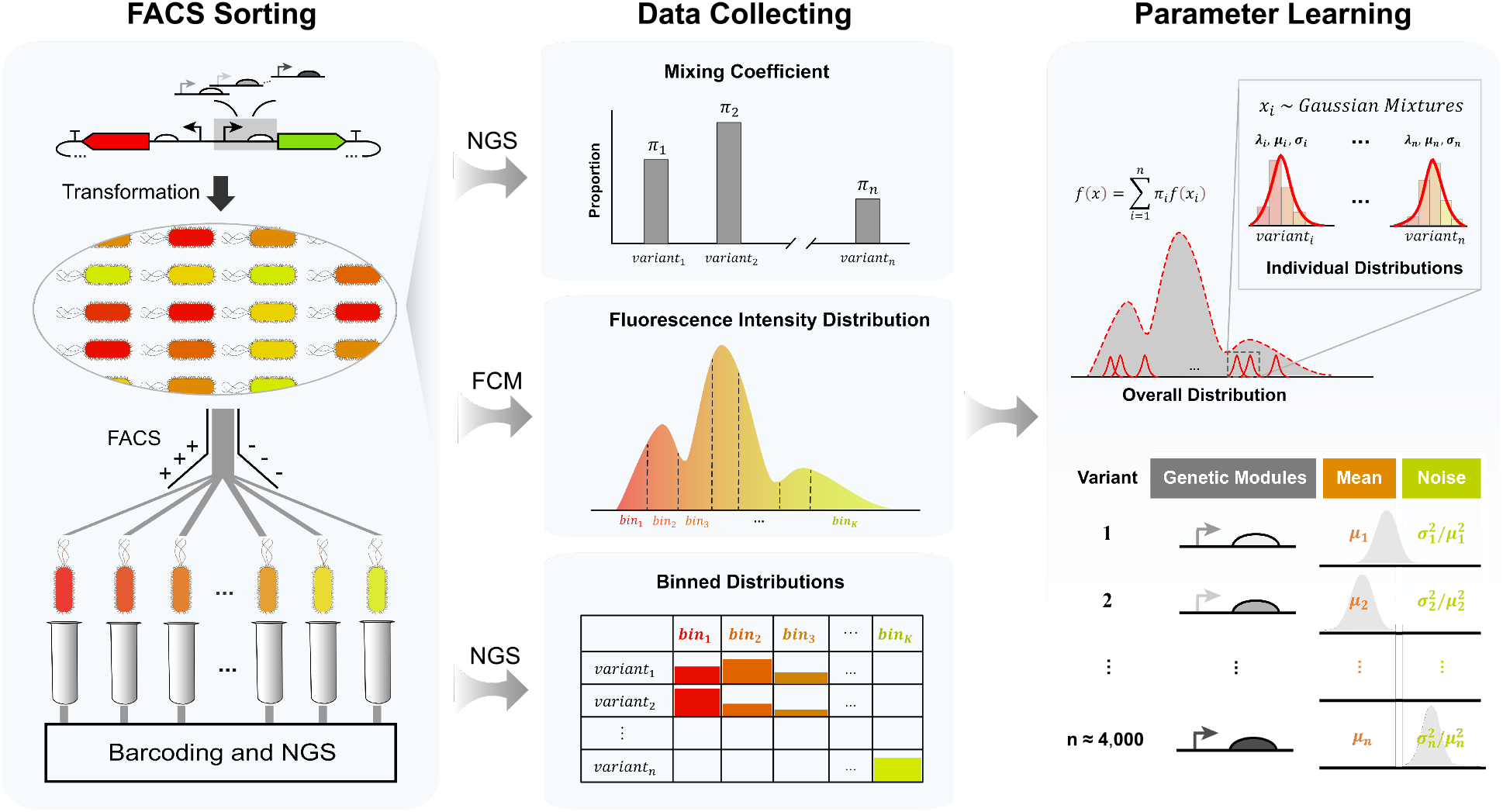
Schematic overview of the dSort-Seq data workflow. (**a**) During Sort-Seq, a library with different expression patterns is sorted into customized bins based on the fluorescence intensity value. (**b**) The mixing coefficients are quantified via next-generation sequencing (NGS). (**c**) The overall fluorescence density is measured by flow cytometry (FCM), and the sorting boundaries are specified based on the overall fluorescence intensity density. (**d**) The read count number across all bins as quantified by NGS reveals the binned distribution of each variant in the library. (**e**) Through parameter learning, the mean, expression noise and their relationships can be precisely identified.

For instance, we applied dSort-Seq to determine the effects of transcription and translation on expression noise in *E. coli* and found them to have comparable contributions, contradicting the commonly accepted translational bursting mechanism^3^. In addition, we also revealed that overlapping RpoD-binding sites would lead to high expression noise, which suggested an effective noise regulation strategy. Overall, our method, which provides significant mathematical and biological insights, can serve as a promising high-throughput tool for use in various studies associated with gene expression.

## RESULTS

### Framework and superior performance of dSort-Seq

Recent research on the stochastic nature of gene expression has shown that steady-state protein production in a clonal population follows a gamma (negative binomial)^3,4^ or log-normal^5,6^ distribution. However, neither of them can precisely match the real expression data (**Fig. 2a-2c**). To address this issue, dSort-Seq applied a two-component log-Gaussian mixture model (LGMM) to represent the steady-state protein production density. This distribution was selected for several reasons, the first and foremost of which is that the mixture of Gaussians can theoretically approximate any continuous density given enough components^31^, ensuring its ability to fit more complex densities compared with conventionally used models. In addition, the outliers (in more extreme cases, one peak of the bimodal expression densities^30^), which have a great impact on matching^19,29^, can be viewed as being generated by a Gaussian component^32–34^. To verify the model’s ability to match expression distributions, we compared it with gamma and log-normal distributions in fitting quantitative datasets from independent resources^27,35^ (**Fig. 2a-2c**). Our method exhibited higher precision in representation, and hence, we used the LGMM for subsequent analyses.

**Figure 2.**
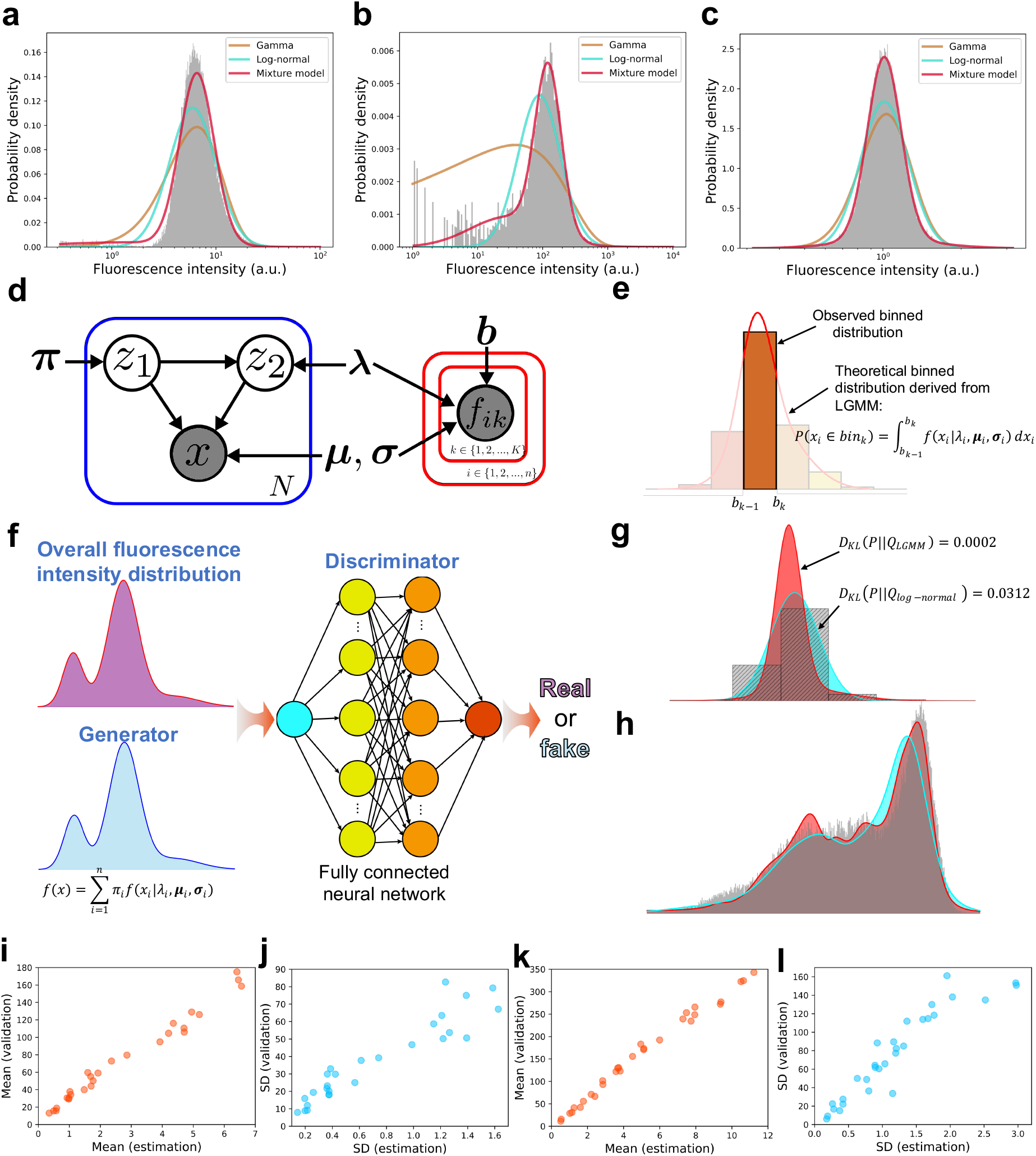
Framework and performance of dSort-Seq. (**a-c**) Two-component log-mixture of Gaussians can better represent the gene expression distribution compared with conventionally used model-driven methods. (**a**) Gene expression controlled by the LmrA repressor^35^; the histogram denotes the cytometry data of the unrepressed state. (**b**) Density of gene expression under the control of the *tnaC* variant K11R_CGC^27^. The data were measured under 100 μM Ala-Trp. (**c**) Gene expression driven by the promoter *yebVp2* (this study). In **a-c**, the red, cyan and brown lines represent the fitting result of the two-component log-mixture of Gaussian, log-normal and gamma distributions, respectively. (**d**) Graphical representation of the model. (**e**) Theoretical fraction of the probability density within the corresponding boundaries. (**f**) Matching the mixture of two-component Gaussian mixture models to the overall fluorescence intensity distribution. The real data are sampled from experimental cytometry data; the fake data are generated from the LGMM. A fully connected neural network is used as a discriminator to determine whether the data are real or fake. (**g**) An example (V8A_GCC, 0 μM Ala-Trp, replicate 1) to illustrate the superior performance of dSort-Seq in matching the binned distribution compared to the log-normal-based method. The Kullback–Leibler divergence shows the performance of each fit. (**h**) An example (100 μM Ala-Trp, replicate 1) to illustrate the superior performance of dSort-Seq in matching the overall fluorescence distribution compared to the log-normal-based method. In (**g**) and (**h**), the red and cyan distributions refer to the results derived from dSort-Seq and the log-normal-based method, respectively. The gray distribution refers to the real data. (**i-l**) Individually analyzed expression characteristics of reconstructed *tnaC* variants by cytometry were highly correlated with those estimated via dSort-Seq in terms of both their means (**i**, 0 μM Ala-Trp, *n* = 26; **k**, 100 μM Ala-Trp, *n* =30) and SDs (**j,** 0 μM Ala-Trp; **l**, 100 μM Ala-Trp).

Next, to derive gene expression characteristics from Sort-Seq data, we revisited the experimental procedure and considered incorporating more data into the parameter learning step (**Fig. 1**). To intuitively represent the dSort-Seq method, we defined the following terms: (1) the mixing coefficients, denoted by **π** = (π_1_, π_2_,…, π*n*), where π_*i*_ is the proportion of the *i*th variant in the library; (2) the log-scaled sorting boundaries, denoted by *b* = (*b*_0_ = −∞, *b*_1_, …, *b_K_* = +∞); (3) the parameters involved in LGMM, denoted by **λ** = (λ_1_, λ_2_, …, λ_*n*_), **μ** = (**μ**_1_, **μ**_2_, …, **μ**_*n*_) and **σ** = (**σ**_1_, **σ**_2_, …, **σ**^*n*^), where **μ**_*i*_ = (μ_1*i*_, μ_2*i*_)^T^, **σ**_*i*_ = (σ_1*i*_, σ_2_*i*)^T^; and (4) the probability of sorting the *i*th variant into the *kth* bin, denoted by *P¿k*. Based on these definitions, we built a Bayesian network to show the data generative process and dependencies among variables (see **Methods**, **Fig. 2d**). For parameter learning, instead of only matching the binned distribution via maximum likelihood estimation as in previous methods^19,21,27^, we constructed a Bayesian neural network to fit both the binned distribution and the overall fluorescence intensity density. Specifically, two objective functions were designed, where the first was defined as the cross-entropy of the observed binned distribution relative to the theoretical binned distribution derived from LGMM (see **Methods**, **Fig. 2e**), by minimizing which the parameters of each LGMM can be optimized for approximation to the observed sorting data. The second objective was aimed at matching the overall fluorescence intensity distribution of the whole library. For this purpose, we applied a generative adversarial network^36^ (**Fig. 2f**). To elaborate, a generator was designed based on the data generative process (see **Methods**). A fully connected neural network was applied as the discriminator. During training, data generated from the generator are sent to the discriminator along with the real fluorescence intensity values, and then the discriminator determines whether each piece of data is real or not. Hence, a two-player game is played between the generator and the discriminator, and the overall fluorescence intensity distribution can be matched by the generator when they are in equilibrium. We included these two objectives in the Bayesian neural network, with which the parameters can be learned through backpropagation (**Supplementary Fig. 1**).

Subsequently, we tested the validity of dSort-Seq with data from our previous Sort-Seq profiling of a comprehensive codon-level mutagenesis library of *tnaC*^27^. This experiment was performed under 3 different ligand concentrations (0, 100 and 500 μM Ala-Trp), each with two biological replicates (**Supplementary Fig. 2a**). However, by fitting each binned distribution to the nonrobust log-normal density, their results were obviously affected by outliers and could not precisely match the experimental observations (see **Methods**) in terms of both the binned distribution (**Fig. 2g**) and the overall fluorescence intensity distribution (**Fig. 2h**). In addition, the expression characteristics derived from the log-normal distribution were also subject to error (see **Methods**, **Supplementary Fig. 5**). Therefore, we applied dSort-Seq in this case to derive the expression properties (see **Methods**, **Supplementary Data 1**). As a result, the strong correlations of the mean (**Supplementary Figs. 2b-2d**, Pearson’s *r* = 0.989, 0.979, and 0.978 for 0, 100, and 500 μM Ala-Trp, respectively) and standard deviation (SD; **Supplementary Figs. 2e-2g**, Pearson’s *r* = 0.914, 0.903, and 0.891 for 0, 100, and 500 μM Ala-Trp, respectively) of expression between biological replicates indicated the reliability of dSort-Seq profiling. In addition, the individual validation data (**Supplementary Figs. 3 and 4**), even if measured via another flow cytometer, were highly consistent with the calculation results (**Fig. 2i**, mean for 0 μM Ala-Trp, Pearson’s *r* = 0.991; **Fig. 2j**, SD for 0 μM Ala-Trp, Pearson’s *r* = 0.942; **Fig. 2k**, mean for 100 μM Ala-Trp, Pearson’s *r* = 0.994; **Fig. 2l**, SD for 100 μM Ala-Trp, Pearson’s *r*= 0.931), which proved the model’s ability to precisely capture the expression characteristics.

### DSort-Seq enables the screening of biosensors with desired response features

Given the superior performance of dSort-Seq in characterizing expression properties, we applied it to practical scenarios to highlight its applicability. First, as an example of expression strength mining, we focused on the metabolite biosensor, through which the intracellular concentration could be converted to a change in gene expression. The key performance indicators of a biosensor include sensitivity, specificity, dynamic range and operational range^37^, most of which can be determined from dose–response relationships. However, to the best of the authors’ knowledge, a method for large-scale profiling of the dose–response curves with high precision is lacking. Therefore, we applied dSort-Seq to address this problem. For instance, we tested it in our previously reported dataset by Zhou et al.^38^, which contained Sort-Seq results for 5,184 FapR-*fapO*-based biosensors, consisting of combinations of 6 transcription factor dosages (*pGPD, pENO2, pHSP12, pEXG1, pCYC1, pULI1*), 4 operator insertion schemes (TATA_OP, OP_TATA, OP_TATA_OP, N30_OP), and 216 arrangements of upstream enhancer sequences (UASs; 3 tandem UASs selected from UAS_A_, UAS_B_, UAS_C_, UAS_D_, UAS_E_ and UAS_F_). Each combination was encoded by a specific DNA barcode to ensure its identification via NGS. The library was transformed into *Saccharomyces cerevisiae* BY4700 and assayed through Sort-Seq under 6 different cerulenin concentrations (0, 1, 2, 3, 5, 8 mg/L), each with two biological replicates (**Fig. 3a**). However, owing to the limited precision and robustness of log-normal-based analysis, the dose–response curves derived from Sort-Seq were imprecise (**Supplementary Fig. 6**) and were inconsistent with the individual characterization data^38^. Therefore, we applied dSort-Seq to this case to determine whether it could accurately evaluate the response performance. The resulting responses showed strong correlations between biological replicates at different concentrations (**Supplementary Fig. 7**, Pearson’s *r* > 0.950 for all experimental conditions), indicating the reliability of the calculation.

**Figure 3.**
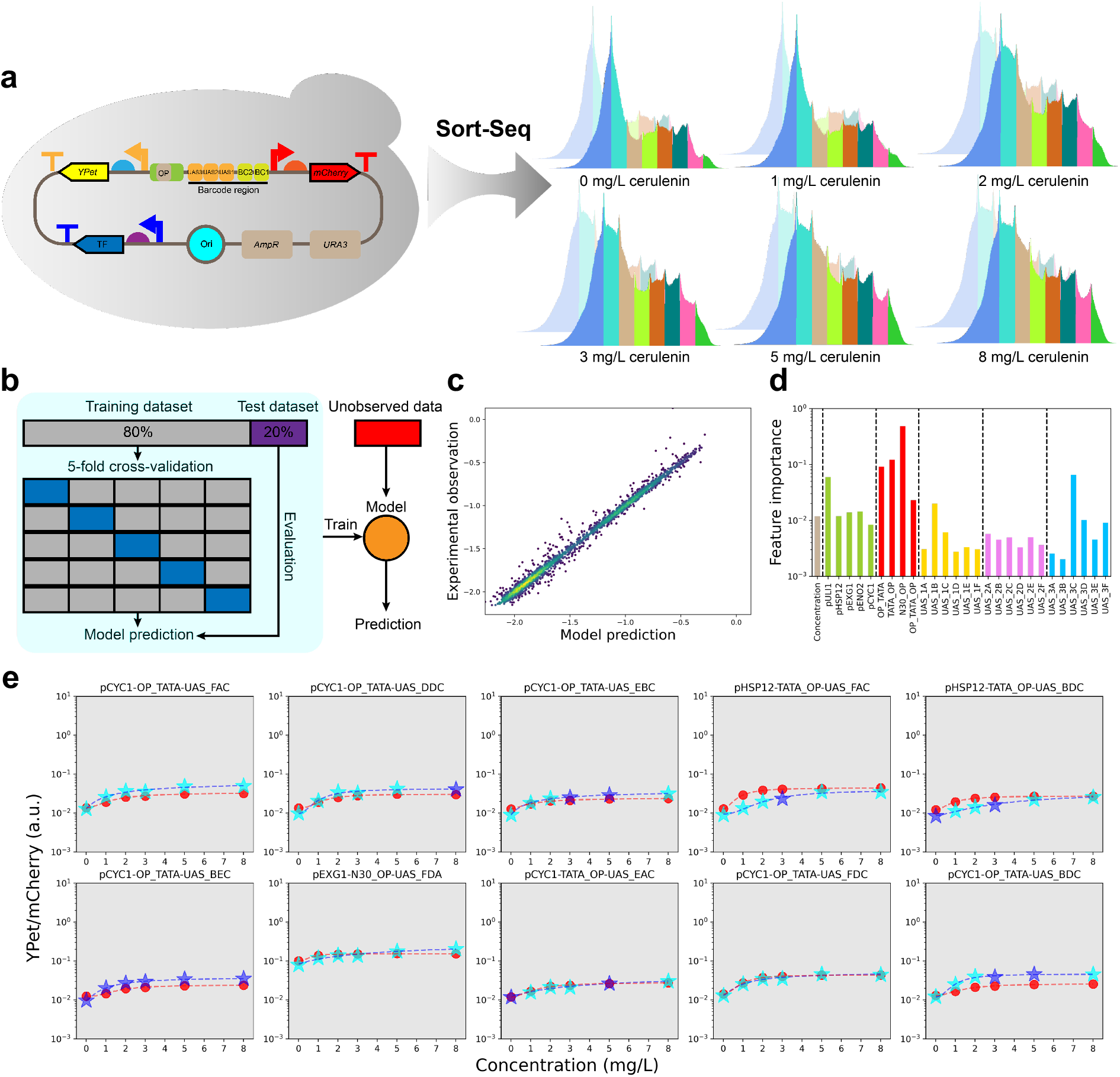
The dSort-Seq profiling of FapR-*fapO*-based malonyl-CoA-dependent gene expression. (**a**) Sort-Seq characterization of the malonyl-CoA biosensor library under 6 different cerulenin concentrations (0, 1, 2, 3, 5, 8 mg/L). Cells were sorted into 8 bins according to their responses to ligand. Two biological replicates were examined for each Sort-Seq experiment. (**b**) Schematic diagram of the machine learning process. Gradient boosting regression was used here to interpret the relationship between features and expression strengths. The hyperparameters were optimized through 5-fold cross-validation; then, the whole training dataset was used to train the model parameters, and the test dataset was used to evaluate the generalization capacity of the model. Finally, the model was trained on the entire observed dataset to obtain predictions for unobserved data. (**c**) The model performance in the test dataset showed a good generalization capacity (*n* = 2,275). (**d**) Gini importance that contributes to the gradient boosting regression tree. (**e**) Dose–response curves of the top 10 combinations with the highest dynamic ranges. Data points represent the mean values of YPet/mCherry under different cerulenin concentrations, where red dots represent individual characterization data, cyan stars represent data from dSort-Seq characterizations, and blue stars denote data from machine learning predictions. The dashed lines represent response curves fitted by the Hill equation (see **Methods**).

As the library was nonuniform, we could obtain only 12,779 expression strengths of 2,616 combinations through dSort-Seq. To obtain the rest of the data, we applied a machine learning approach to generate predictions. Specifically, each combination was encoded as a 27-dimensional vector (see **Methods**), along with the cerulenin concentration as the input feature. Gradient boosting regression was applied to fit the log-scaled expression strength. Note that as combinations containing the promoter *pGPD* suffered from a heavy metabolic burden, leading to them being underrepresented in the library, we excluded them from the machine learning analysis. Therefore, the data used to train the model contained 11,375 responses, covering 43.9% of the whole combinatorial space (**Supplementary Data 2**). To avoid overfitting, we randomly split the dataset into two subgroups, with 80% of the data used as the training dataset to optimize the hyperparameters through 5-fold validation as well as train the model parameters. The remaining 20% were used as the test dataset to check the generalization capacity of the model (**Fig. 3b**). The performances in the test dataset (*r*^2^ = 0.989, **Fig. 3c**) indicated that the model had reasonable generalization capacity and captured biological signals. Subsequently, we trained the model on the whole dataset and predicted the uncharacterized responses. The dSort-Seq data accompanied by the predicted results were then applied to generate the dose–response curves for all combinations (**Supplementary Data 2**). We validated these dose–response relationships using 92 individual characterization results^38^, and linear regression was applied to fit the data values within the same scale. The resulting high consistencies (Pearson’s *r* > 0.970 for all cases, **Supplementary Figs. 8 and 9**) demonstrated that with dSort-Seq and machine learning, the expression properties of the enormous combinatorial space could be effectively explored.

We then analyzed the features that contributed most to model predictions via Gini importance (**Fig. 3d**). Overall, consistent with previous discoveries^38^, the responses of the biosensor were mostly affected by the operator insertion schemes. In addition, a strong determinant of the result was observed if the third UAS was UAS_C_. Next, we focused on the dynamic range, which measures the signal-to-noise ratio of a biosensor, by increasing which the true signal is more likely to be discerned from noise. Hence, we fitted each dose–response relationship to the Hill equation to derive the corresponding dynamic range (see **Methods**). The top 10 combinations with the highest dynamic ranges were individually constructed and assayed through FCM (**Fig. 3e**). Their response performances were consistent with dSort-Seq and machine learning calculations, of which pHSP12-TATA_OP-UAS_FAC achieved the highest dynamic range of 3.5. Notably, the third UAS of most of the 10 combinations was UAS_C_, indicating that UAS_C_ is important for the interactions of yeast synthetic promoters with FapR when it is located at the third position. In addition to dynamic range, other indicators, including operational range and sensitivity, can also be evaluated and optimized in a similar manner. Therefore, with dSort-Seq, the optimal design with desired response features can be effectively identified.

### DSort-Seq profiling of the mean noise landscape of *E. coli* endogenous promoters

In addition to the expression strength, expression noise is also an important factor affecting gene expression that leads to phenotypic diversity among genetically identical individuals. Previous association studies have found that expression noise is a heritable trait^39^ and is determined by expression modules^14–16,30^. Hence, for a given organism, how different expression modules shape the patterns of noise is a fundamental question. On the other hand, in terms of noise production mechanisms, the commonly accepted translational bursting model suggests that the protein within a cell is produced in bursts, where the burst size (noise strength, *η* = SD^2^/Mean) is related to only translation and is independent of transcription^3,4^ (**Fig. 4b**). However, relevant experimental evidence is still scarce. The problem with the translational bursting mechanism is not only that the gamma distribution cannot accurately represent the gene expression distribution^6^ but also that it cannot interpret the dependence between transcription and noise strength^15^. However, limited by the low throughput of classic quantitative methods, research on transcriptional and translational contributions to expression noise is always based on the analysis of a small amount of data^15,40^, which is susceptible to experimental error as well as outlier samples. To address these issues, our approach may serve as a promising method due to its ability to produce high-quality data in a massively parallel manner. Therefore, as a proof of concept, we performed systematic profiling of transcriptional effects on expression noise in *E. coli* based on dSort-Seq.

**Figure 4.**
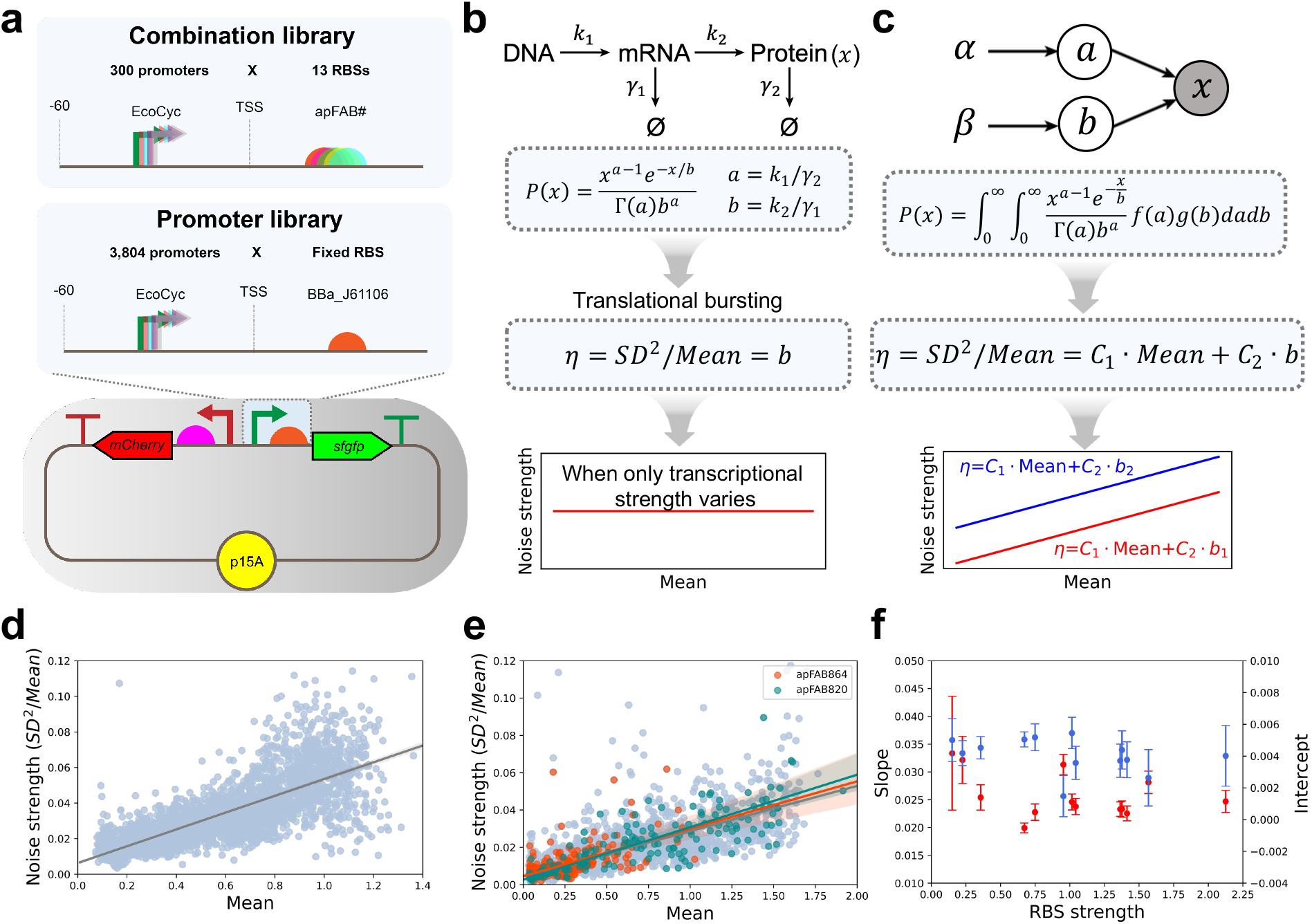
The dSort-Seq profiling of transcriptional and translational effects on noise production in *E. coli* K12 MG1655. (**a**) The design schemes of the promoter and the combination library. (**b**) According to the translational bursting mechanism, steady-state protein production follows a gamma distribution^3^; as a corollary, the burst size, denoted by the Fano factor, is linearly correlated with the translation rate and independent of the transcription rate. (**c**) According to the hierarchical Bayesian model, the intercept in the relationship between noise strength and the mean expression level is proportional to translational strength, indicating that translational bursting still dominates noise production at low expression levels^30^. (**d**) The noise strength is linearly correlated with the mean expression level when only transcriptional strength varies. The gray line exhibits the linear regression result, which is shaded to show the 95% confidence interval. (**e**) The relationships between noise strength and mean expression level are similar when the translation module varies. The gray, orange, and green lines represent the regressions of all combinations and combinations with RBS apFAB864 and apFAB820, respectively. (RBS strength: apFAB820 (1.57) > apFAB864 (0.36), see **Methods**). (**f**) The linear regression slopes and intercepts of noise strength and mean expression level are not significantly correlated with translational strength. The error bars indicate the 95% confidence intervals.

For library construction, 3,804 endogenous promoters of *E. coli* K12 MG1655 were collected from the EcoCyc database^41^ (https://www.ecocyc.org/), for which the 60-nt sequence upstream of each transcription start site was regarded as the promoter region. The oligonucleotide library composed of collected promoters was high-throughput synthesized and assembled into a low-copy-number, dual-fluorescence plasmid, pMPTPV_dual_fluorescence, in which a superfolder green fluorescent protein (sfGFP) was used as the response reporter that was under the control of a particular promoter with a fixed ribosome-binding site (RBS; BBa_J61106). In addition, a constitutively expressed reporter, mCherry, served as an internal reference to eliminate cell-to-cell variations such as cell volume and plasmid copy number. After electroporation into MG1655 cells, a cell library with broad levels of sfGFP expression was obtained. The cultivated cell library was characterized by FCM with three biological replicates and then sorted into 12 bins based on the fluorescence intensity of sfGFP relative to mCherry, followed by NGS to quantify the proportion of each variant in each bin. The acquired datasets were then processed and analyzed by our method (**Fig. 4a**). The results showed that 2,920 (76.8% of the total library) promoters were highly consistent among all three replicates (**Supplementary Fig. 15**, **Supplementary Data 3**). Validation of this result was carried out through individual cytometry assays of 60 randomly picked single colonies (**Supplementary Fig. 16**). The strong consistency with the dSort-Seq results (**Supplementary Fig. 17**, Pearson’s *r* = 0.981 and 0.921 for the mean and SD, respectively) indicated the reliability of the profiling.

Autofluorescence was quantified by assaying the pMPTPV strain with only mCherry expression and no sfGFP expression, and the result showed that autofluorescence could be neglected relative to the fluorescence intensity of each candidate of the library (**Supplementary Fig. 12**).

The bulk data generated a comprehensive landscape of promoter strength and expression noise along the *E. coli* genome (**Supplementary Data 3**, we also visualized it through D3GB^42^ at http://www.thu-big.net/Escherichia_coli_K12_MG1655_promoters), which was beneficial for understanding the transcriptional strategies for different genes. For instance, we investigated whether essential and nonessential genes of *E. coli* possess different expression patterns (see **Methods**). As a result, the essential genes showed greater transcriptional intensities than the nonessential genes (**Supplementary Fig. 18**, *P* = 4.22e-16, one-tailed t test). Given that the high transcriptional strength is usually related to low expression noise^15^, these functionally important genes are more likely to confer lower levels of noise, which is consistent with the results of a previous genome-wide association study^16^. Subsequently, we investigated the relationship between noise strength and mean expression level. The results showed that the noise strength was linearly correlated with the expression strength when the transcription module varied (**Fig. 4c**, Pearson’s *r* = 0.745); hence, transcription contributed to expression noise. This discovery, however, is inconsistent with the inference of the translational bursting model (**Fig. 4b**), suggesting the limitation of the model in interpreting noise production mechanisms in *E. coli*.

### Transcription and translation make comparable contributions to noise production

Although the translational bursting mechanism is unable to account for the contribution of transcription to noise production, the hierarchical Bayesian model, developed by introducing transcriptional and translational fluctuations into the translational bursting model, can successfully explain this phenomenon^30^ (**Fig. 4c**). However, the model showed that different translation modules would lead to varying intercepts^15,30^ in the relationship between noise strength and the mean expression level (**Figure 4c**, *η* = *C*_1_ · Mean + *C*_2_ · *b*, where *b* represents the translation strength). Hence, at low levels of transcription, the expression noise is still dominated by translation. This conclusion, although confirmed by a fluorescence microscopy experiment that analyzed 40 *B. subtilis* strains expressing GFPmut3 with different combinations of transcription and translation modules^15^, still needs to be verified, as it was based on regression analysis of a small sample. To date, the contribution of transcription and translation to noise production has not been comprehensively and directly observed. As our method has greatly expanded the test throughput of expression noise, we applied dSort-Seq here to examine these features.

Therefore, we designed a combination library comprising different combinations of 300 promoters and 13 RBSs. The promoters were randomly selected from the EcoCyc database, whereas the RBSs, which were chosen for their varying translational strengths, were from the apFAB# series^43^. The combination library was prepared in the same manner as the promoter library in the pMPTPV_dual_fluorescence plasmid. After electroporation, we performed a dSort-Seq assay of the cell library, with three independent biological replicates to ensure the reliability of the results. After data processing, 2,733 combinations (70.1% of the whole library) were highly consistent among the replicates and were retained for subsequent analysis (**Supplementary Fig. 20**, **Supplementary Data 4**). Subsequently, 60 single colonies with different genotypes were randomly picked and assayed individually with FCM (**Supplementary Fig. 21**). Their means and SDs of expression were strongly correlated with the dSort-seq results (**Supplementary Fig. 22**, Pearson’s *r* = 0.976 and 0.937 for the mean and SD, respectively), proving the validity of the profiling.

We then performed regression analysis between noise strength and the expression mean for different translational modules (**Fig. 4e**, **Supplementary Fig. 24**) to test the hierarchical Bayesian model. However, their correlation barely changed when the translation module varied in terms of both slope and intercept (**Fig. 4f, Supplementary Table 6**). Instead, our results showed that the noise strength was highly coupled with the mean expression level, indicating the difficulty of adjusting expression noise independently of the mean protein abundance by tuning the strength of the transcription and translation modules. Furthermore, to determine whether translation bursting dominates noise production at low transcription levels, we constructed several weak expression combinations and performed cytometry assays. As a result, the combinations with comparable mean expression levels showed similar fluorescence intensity distributions (**Supplementary Fig. 25**), suggesting that the contributions of transcription and translation to noise are comparable, even in the case of weak expression strength.

### Overlapping RpoD-binding sites can lead to high expression noise

We then analyzed the relationship between expression noise (CV^2^ = SD^2^/Mean^2^) and strength (**Fig. 5a and 5b**). The expression noise exhibited a strong negative correlation with the mean protein abundance at low levels of expression and then reached a plateau after a critical point, which is consistent with previous observations^15,30^. In addition to the general correlation, some unique expression features also piqued our interest, especially for those sequences exhibiting high expression noise at their corresponding mean expression levels. To ensure that these outliers were not the results of experimental error, we reconstructed and assayed 20 high-noise candidates from the promoter library and 25 from the combination library and then performed an individual cytometry assay (see **Methods**), proving the credibility of the discovery. Moreover, the expression noise of these variants was apparently higher than that of randomly selected colonies (**Fig. 5c and 5d**). Notably, among the 25 high-noise combinations, several promoters appeared frequently (e.g., *fliEp1, ileSp2, yihVp3, folEp*, **Fig. 5d**), suggesting that the extra noise may be derived from transcription rather than translation.

**Figure 5.**
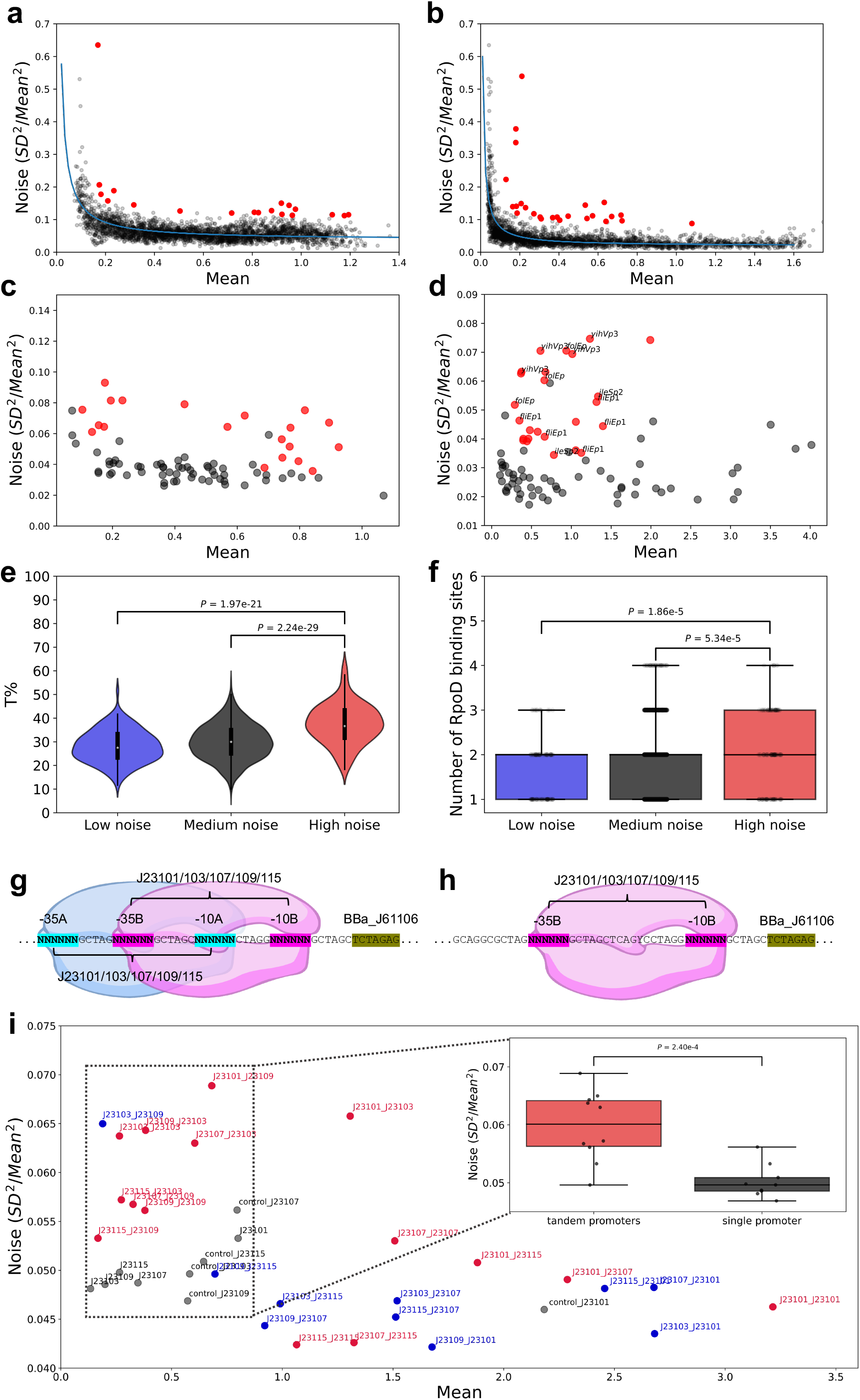
Overlapping RpoD-binding sites result in high expression noise. (**a** and **b**) Correlation of the expression noise with expression strength in the (**a**) promoter library and (**b**) combination library. At low mean expression levels, the noise decreases as the expression strength increases; at high mean expression levels, the noise converges to a constant value. The blue lines show the regression results (see **Methods**). Twenty promoters and 25 combinations exhibiting high expression noise are marked as red dots. (**c**) Twenty sequences from the promoter library and (**d**) 25 sequences from the combination library showing high expression noise were constructed and assayed through FCM. As a result, their expression noise (indicated by red dots) is higher than that of randomly selected variants (indicated by black dots) at their corresponding mean expression levels. (**e**) The high-noise group had a significantly higher thymine content than the low-noise group (*P* = 1.97e-21, one-tailed t test) and the medium-noise group (*P* = 2.24e-29). (**f**) The number of potential RpoD-binding sites in the high-noise group was significantly higher than that in the low-noise group (*P* = 1.86e-5, one-tailed t test) as well as in the medium-noise group (*P* =5.34e-5). (**g**) Design scheme of 25 tandem promoters, each containing two overlapping RpoD-binding sites. (**h**) Design scheme of 5 constitutive promoters with the same length as the tandem promoter, each with only one RpoD-binding site. (**i**) Compared to promoters with a single RpoD-binding site, the tandem promoters exhibited significantly higher expression noise (*P* = 2.40e-4, one-tailed t test), especially when the stronger promoter was located upstream of the weaker promoter (indicated by red dots).

Next, to identify the factor that contributed to the additional expression noise, we divided the *E. coli* promoters into 3 groups based on the noise values (high-noise, medium-noise and low-noise groups; see **Methods**). Subsequently, we analyzed the sequence features of the three groups and found that the thymidine proportion was significantly higher in the high-noise group (average 37.5%, *n* = 138) than in the low-noise group (**Fig. 5e**, average 28.0%, *n* = 138; *P* = 1.97e-21, one-tailed t test) as well as in the medium-noise group (average 29.9%, *n* = 2,481; *P* = 2.24e-29). Furthermore, there was no position preference for this phenomenon (**Supplementary Fig. 30**), and no common regulator was found to be associated with the sequences within the same group (**Supplementary Data 5**). Therefore, we focused on transcription initiation factors, especially RpoD (σ^70^ factor), which transcribes most genes in *E. coli*. Promoters recognized by RpoD generally contain two consensus hexamers centered at 10 and 35 nucleotides upstream of the transcription start site. These two regions are rich in adenosine and thymine, especially thymine (average of 4.73 per promoter compared to 3.75 for adenosine, 1.80 for cytosine and 1.70 for guanine^44^). Based on this, we hypothesized that the high-noise group contains more RpoD-binding sites than other groups. To test this hypothesis, we searched the DPinteract database^45^ for potential RpoD-binding sites in three groups (see **Methods**). The sequences with no RpoD-binding hit were excluded from subsequent analysis. As a result, the high-noise group (average 2.10, *n* = 127) showed more potential RpoD-binding sites than the low-noise group (**Fig. 5f**, average 1.65, *n* = 106; *P* = 1.86e-5, one-tailed t test) and medium-noise group (average 1.82, *n* = 2,091; *P* = 5.34e-5).

Subsequently, to determine whether overlapping RpoD-binding sites would result in high expression noise, we constructed 25 tandem promoters based on combinations of 5 Anderson promoters (J23101/103/107/109/115; **Fig. 5g**) to drive the expression of *sfgfp*. In addition, the 5 constitutive promoters were also individually constructed as controls. To exclude the effect of promoter length and the −35 region-proximal sequence on the results, we also constructed 5 promoters of the same length as the tandem promoter, while preserving only the downstream RpoD-binding site (**Fig. 5h**). Subsequently, we performed an individual cytometry assay of the 35 promoters (**Supplementary Fig. 32**). As a result, the gene expression driven by tandem promoters showed higher noise compared to a single promoter (**Fig. 5i**, *P* = 2.40e-4, one-tailed t test), especially when the stronger promoter was located upstream of the weaker promoter (**Fig. 5i**), suggesting that the transcriptional interference caused by the occlusion of promoters contributed to noise production. Hence, our results uncovered a feasible and simple noise modulation strategy in *E. coli* by tuning the number and relative positions of sigma factors upstream of the transcription start site.

## DISCUSSION

Gene expression dosage is directly associated with a variety of phenotypes of a population^46^; hence, there is no doubt that high-throughput profiling will deepen our understanding of cellular behavior. In this paper, we focused on biosensors that can sense metabolite concentrations and regulate gene expression. According to the different output signals, biosensors have various applications, including in high-throughput screening^47^, medical diagnosis^48^, and cell imaging^49^. For each application, the dose–response relationship is a key indicator that needs to be tuned to meet practical needs. Fortunately, dSort-Seq was shown to be a powerful tool for characterizing and optimizing the biosensor response performance in a high-throughput and high-precision manner, enabling the engineering of biosensors with desired properties. Compared to positive and negative screening^50,51^, by which only the dynamic range could be optimized, dSort-Seq can yield more comprehensive information to meet the needs of various situations. In addition, it is much more efficient than traditional trial-and-error approaches. Therefore, dSort-Seq provides a solution for profiling the expression landscape of the combinatorial sequence space. On the other hand, the high-quality dSort-Seq dataset also has the potential to serve as a basis for deciphering physiological mechanisms^25,27^.

Noise in biological systems has been widely demonstrated to influence various intracellular processes^52^ and the physiological properties of a population^13,53^, although the effects of noise strength vary across different situations. For instance, low noise can ensure stable biosynthesis pathways and robust synthetic gene circuits^54^. In contrast, high phenotypic variability promotes evolvability^55–57^. Therefore, it is necessary to understand the origin of the noise, as well as control noise rationally for various applications. Regarding noise regulation, various strategies have been proposed to control gene expression noise independently, including engineering transcription and translation in synthetic gene circuits^58,59^, introducing pulsatile input to control the promoter activation frequency and transcription rate independently^60^ and expressing two copies of the target gene from separate circuits with different characteristics^61^, among others. Through dSort-Seq profiling of different combinations of promoters and RBSs in *E. coli*, the transcriptional effect on gene expression noise was revealed. Specifically, a higher thymidine proportion without position preference in the promoter sequence would lead to a higher level of noise. One hypothesis to explain this phenomenon is that RpoD-binding sites, which are rich in thymine, influence noise production. We have proven that promoters with overlapping RpoD-binding sites contribute to noise production due to occlusion of promoters. Moreover, in *E. coli*, 831 genes have been found to be under the control of tandem promoters^62^, suggesting the broadness of the regulatory scheme. Hence, in-depth research on modeling molecular events in the transcription process is needed to elucidate the effect of promoter architecture on expression noise.

From the methodology point of view, the design-build-test-learn (DBTL) cycle is emerging as a key workflow in synthetic biology, where the test is the rate-limiting step due to its low throughput^63^. The development of Sort-Seq has undoubtedly greatly extended the test throughput, enabling more efficient characterization and optimization of biological parts. The dSort-Seq approach shows that the learn can be encapsulated into the test to improve its capability by modeling the data generative process for the high-throughput experiment. Moreover, it is worth mentioning that as the ability of the Gaussian mixture models in distribution matching can be improved by increasing the number of mixture components, dSort-Seq can be easily transferred to more complex situations in which multiple feedback circuits are involved^64^. Thus, this pipeline has great potential to determine the mean-noise space for various gene expression modules, providing diverse synthetic parts that can be applied to different fields, such as biosynthesis^65^, laboratory-based adaptive evolution^56^, transcriptional regulation^66^, and protein–protein interactions^67,68^. Overall, owing to the flexibility, high precision and high throughput of this method, we believe dSort-Seq can serve as a powerful tool that provides a wide range of novel research opportunities.

## METHODS

### Parameter-learning algorithm of dSort-Seq

We represent the log-scaled expression density of each variant by a two-component Gaussian mixture model (where *x_i_* denotes the log-scaled intensity value of the *i*th variant):

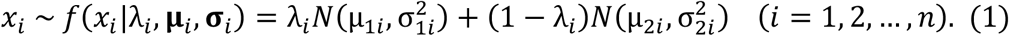

Hence, the overall logarithmic fluorescence intensity distribution can be modeled as a mixture of Gaussian mixture models:

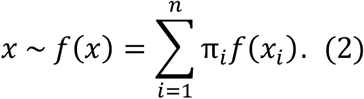

The generative process for each fluorescence intensity value should be as follows: (1) choose a variant *z*_1_ ~ *Categorical*(**π**), (2) choose a Gaussian component *z*_2_ from *Bernoulli*(*z*_2_|*z*_l_) and (3) choose a log-scaled intensity value *x* from *Z*(*x*|*z*_1_, *z*_2_) (**Fig. 3d**). The distributions of variables involved in the model are listed as follows:

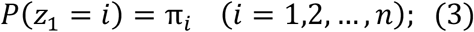

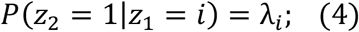

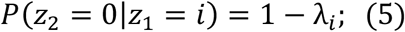

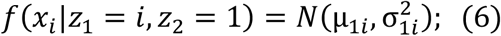

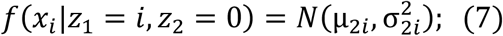

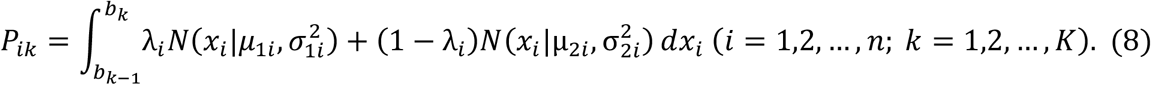

Among the parameters involved in the model, the sets **λ**, **μ** and **σ** cannot be identified experimentally. To estimate them, we designed a probabilistic artificial neural network in which a double-objective optimization is performed. The first objective function is defined as the cross-entropy (H) of the observed binned distribution relative to the integral of the probability density over adjacent boundaries (**Fig. 3e**), which is shown as follows:

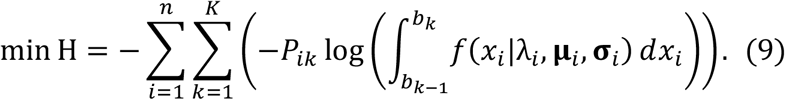

By minimizing the above loss function, the binned distribution can be fitted. The other objective is to match the overall fluorescence intensity density. To this end, a generative adversarial network^36^ is applied. Specifically, a generator is constructed based on the abovementioned generative process. For the discriminator, a fully connected neural network is used to determine whether the data are real or fake (**Fig. 3f**). During training, a two-player game is played between the generator *G* and the discriminator *D* with value function *V*(*G, D*):

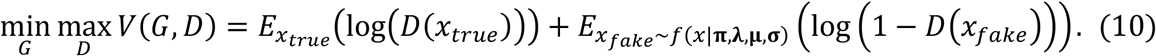

Combining the above two parts, we can obtain the whole algorithm for parameter learning, as shown in **Supplementary Fig. 1**.

### Obtaining expression characteristics from cytometry data

For each individual cytometry assay, the log10-transformed fluorescence intensity distribution was fitted by a two-component Gaussian mixture model (**Supplementary Figs. 3, 4, 12, 16, 21, 25, 26, 28 and 32**) via the expectation–maximization (EM) algorithm, which resulted in a representation of 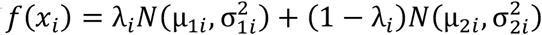. The mean expression strength was calculated with Eq. 11.

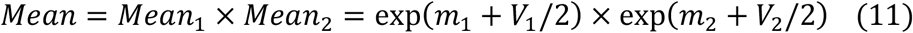

where *m_i_* = *λ_i_μ*_1*i*_log(10), *V*_1_ = (*λ*_*i*_,σ_1*i*_log(10))^2^, *m*_2_ =(1 – *λ_i_* log(10) and *V*_2_ = ((1 – *λ_i_*)σ_2*i*_log(10))^2^. The SD of each expression density was calculated with Eq. 12.

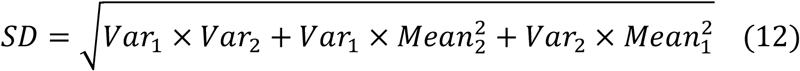

where *Var*_1_ = exp(2*m*_1_ + *V*_1_) (exp(*V*_1_, – 1)), *Var*_2_ = exp(2*m*_2_ + *V*_2_) (exp(*V*_2_ – 1)).

### Comparison of dSort-Seq and the log-normal-based method

The dSort-Seq results were calculated as mentioned above, and the log-normal results were obtained from previously reported data^27^ (note that since the actual slope of the sorting boundary lines on the log-log plot of eGFP-mCherry in the experiment was 0.8810 instead of 1, each boundary value was shifted to the right by 0.3801 compared to the previous analysis, **Supplementary Table 4**). Next, as an example, we compared the performances of the two methods in matching the binned distribution of variant V8A_GCC under 0 μM Ala-Trp and calculated the Kullback–Leibler divergences (Eq. 13) of the observation from the theoretical binned distributions derived from the log-normal-based method and dSort-Seq. The results showed that dSort-Seq is more precise and robust than log-normal (**Fig. 2g**).

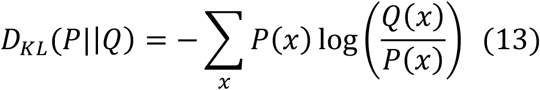

Subsequently, we compared these two methods in fitting the overall fluorescence intensity distribution. For instance, we calculated the theoretical log-scaled overall distribution (100 μM Ala-Trp, replicate 1) derived from the log-normal-based method (Eq. 14) and dSort-Seq (Eq. 2). As a result, dSort-Seq also showed better performance (**Fig. 2h**).

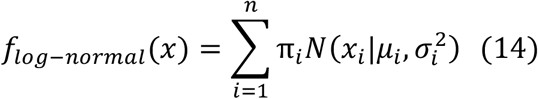

Moreover, as mentioned above, the log-normal-based method is unable to fit the individual cytometry data, which usually serve as criteria for validating the Sort-Seq results (**Supplementary Figs. 3 and 4**). To measure the error, we calculated the expression strength and SD of individual validation data with both log-normal (Eq. 15 and Eq. 16) and the LGMM (Eq. 11 and Eq. 12), where the LGMM results were applied as ground truth to evaluate the precision of log-normal results. As a result, the response and SD inferred from log-normal showed significant deviations (**Supplementary Fig. 5**). Therefore, with log-normal, it is difficult to infer accurate expression properties from Sort-Seq experiments.

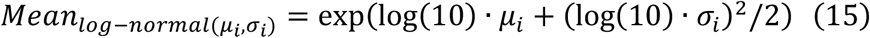

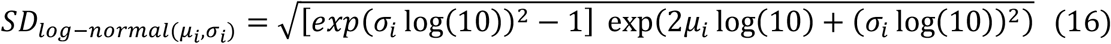

### DNA manipulation and reagents

Plasmid extraction and DNA fragment purification were performed using kits from Omega Bio-Tek. PCRs were carried out using a KAPA HiFi PCR Kit from KAPA Biosystems. The restriction enzyme FastDigest *Esp*3I (namely, *Bsm*BI) and T4 DNA ligase were purchased from Thermo Scientific. Cerulenin was ordered from Yuanye Bio-Technology. All strains and plasmids used in this work are summarized in **Supplementary Table 1**. All oligonucleotides (**Supplementary Table 2**) were ordered from Azenta. Molecular cloning was performed with *E. coli* DH5α (BioMed) as the host. The concentrations of the antibiotics kanamycin and ampicillin were 50 mg/L and 100 mg/L, respectively. In all experiments, bacteria and yeast were grown at 37 and 30°C, respectively.

### Featurization and gradient boosting regression

We applied one-hot encoding to transform each biosensor combination into a 27-dimensional vector. Among these dimensions, 5 of them represent the promoters of the transcription factor (*pULI1, pHSP12, pEXG1, pENO2, pCYC1*), 4 of them represent the operator insertion schemes (OP_TATA_OP, TATA_OP, N30_OP, OP_TATA), and 18 of them represent the sequences of the tandem UAS (UAS_1A/B/C/D/E/F, UAS_2A/B/C/D/E/F, UAS_3A/B/C/D/E/F). These vectors then served as input features along with the cerulenin concentration (0/1/2/3/5/8). Gradient boosting regression was applied to predict the log-scaled expression strength. During training, the hyperparameters were optimized following the given order (min_samples_split, max_depth, min_samples_leaf, max_features, subsample, learning_rate and n_estimators) through the grid search method.

### Fitting the dose–response relationship to the Hill equation

Each dose–response relationship was fitted by Eq. 17 via nonlinear least squares.

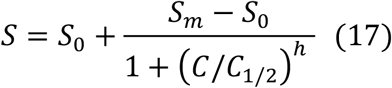

where *S*_0_ and *S_m_* are the values of the sensor response at zero and saturating ligand concentrations, *C*_1/2_ is the concentration at half saturation, and *h* is the Hill coefficient. The lower bounds and upper bounds of (*S*_0_, *S_m_, C*_1/2_, *h*) were set to (0, 0, 0, 1) and (1, 2, 8, 3), respectively. The dynamic range was calculated with Eq. 18.

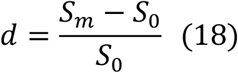

### Individual characterization of the dose–response relationships for malonyl-CoA biosensors

One variant (pHSP12-TATA_OP-UAS_FAC) was obtained from library stock, and the other 9 biosensor variants (pCYC1-OP_TATA-UAS_FAC, pCYC1-OP_TATA-UAS_DDC, pCYC1-OP_TATA-UAS_EBC, pHSP12-TATA_OP-UAS_BDC, pCYC1-OP_TATA-UAS_BEC, pEXG1-N30_OP-UAS_FDA, pCYC1-TATA_OP-UAS_EAC, pCYC1-OP_TATA-UAS_FDC, pCYC1-OP_TATA-UAS_BDC) were constructed via Golden Gate Assembly (**Supplementary Table 3**). After transformation of these plasmids into BY4700, the strains were inoculated into 48-well deep-well plates with 1 mL of SC-Ura medium (synthetic complete medium lacking uracil) in each well. After culturing for 12 h at 30°C and 250 rpm, 2 μL of cerulenin solutions of six distinct concentrations (0.5, 1.0, 1.5, 2.5, 4 mg/mL) was added to the corresponding well. The strains were then cultured for another 12 h. For sample preparation, cells were collected by centrifugation (4°C; 8,000 × g for 10 min) and resuspended in prechilled phosphate-buffered saline (PBS) to an OD_600_ of 2. BY4700 was used as a negative control. BY4700/POT1-pTEF2-mCherry-tADH1 and BY4700/POT1-pCYC1-YPet-tPGK1 were used as positive controls for mCherry and YPet, respectively. These control samples were prepared the same way as above. The fluorescence intensities of the cells were characterized on an LSRFortessa (BD Biosciences). The double-positive area, named Q2, was determined by the control samples, as described in a previous work^38^. For each sample, 100,000 events in the Q2 area were analyzed.

### Construction of the two-reporter plasmid

The two-reporter plasmid pMPTPV_dual_fluorescence was derived from the common vector pACYCDuet-1 by replacing the chloramphenicol resistance gene with the kanamycin resistance gene, replacing the *lacI* expression cassette with *mcherry*, and inserting the *sfgfp* cassette into the opposite strand of *mcherry*. The *mcherry* gene is controlled by a constitutive promoter, pL_M1-37^69^. To facilitate library construction, *sfgfp* is controlled by a variable region containing two *Bsm*BI restriction sites.

### Feasibility verification of the plasmid

Ten promoters were randomly selected from the EcoCyc database. The sequence of each promoter was defined as the 60 nt preceding the transcriptional start site. In addition, a medium-strength RBS, BBa_J61106 (TCTAGAGAAAGATAGGAGACACTAGT), was chosen for all strains to ensure the survival of strains (note that the combination of a strong promoter and a strong RBS is lethal to *E. coli*^17^). After transformation of the plasmids containing different promoters into *E. coli* K12 MG1655, the resulting strains were individually cultured in LB medium containing kanamycin (initial OD_600_ = 0.02), with three biological replicates for each promoter. During cultivation, the growth rate and the expression of fluorescent protein were monitored by sampling and testing every hour. The OD_600_ was measured by a microplate reader (Tecan Infinite 200Pro), and the fluorescence intensity was assayed via flow cytometry (BD LSRFortessa). The 10 strains showed no apparent differences in growth (**Supplementary Fig. 10a**), and the median value of sfGFP/mCherry remained stable after culturing for 16 h (**Supplementary Fig. 10b**). Hence, we chose 16 h for cultivation in subsequent experiments.

### Preparation of library cells

The two plasmid libraries (the promoter library and combination library) were both ordered from Genewiz. We transformed each library into *E. coli* K12 MG1655 via a BTX Harvard ECM 630 High Throughput Electroporation System using optimized parameter settings (2.1 kV, 1 kΩ, 25 μF, 100 ng plasmids/100 μL competent cells). The transformed cells were incubated in LB medium (four times the volume of the competent cells) for 1 h at 37°C for recovery and then plated onto 37 Φ150 LB agar plates containing kanamycin with an EasySpiral Pro (Interscience). Generally, ~10^4^ single colonies per plate can be harvested with this protocol (data not shown), enabling ~100 times coverage of the designated library. All colonies on the plates were rinsed off using sterile LB medium supplemented with kanamycin, collected by centrifugation (4°C; 8,000 × g for 10 min) and then resuspended and thoroughly mixed to an OD_600_ of 10 using fresh sterile LB medium containing kanamycin. The cell suspension was stored at −80°C in glycerol (final OD_600_ = 5).

### Characterization of transcriptional impact on growth

We further tested whether different promoters have varying influences on growth. To this end, the stored promoter library cells were cultured in LB medium containing kanamycin (initial OD_600_ = 0.02) at 37°C for 16 h. Cell samples were collected before and after cultivation, followed by plasmid extraction. The promoter regions were amplified by PCR (KAPA HiFi PCR Kit; 95°C for 3 min, 25 cycles [98°C for 20 s, 63°C for 15 s, 72°C for 5 s], 72°C for 30 s) using the Lib_F and Lib-R primers (**Supplementary Table 2**).

In a 50-μL reaction, 5 ng of the plasmid library was added as the PCR template. The sequencing library was prepared according to the NEBNext Ultra II DNA Library Prep Kit for Illumina (NEB). Specifically, 30 ng of each purified PCR product was used to prepare the sequencing library. The DNA fragments were treated with NEBNext End Prep for end repair, 5’ phosphorylation and dA-tailing. Then, the fragments were ligated to NEBNext Adaptors, followed by USER Enzyme excision. Subsequently, the products were purified using NEBNext Sample Purification Beads and amplified by PCR for six cycles using the P5 and P7 primers. The products were again purified using NEBNext Sample Purification Beads, validated with an Agilent 2100 Bioanalyzer (Agilent Technologies) and quantified with a Qubit 4 Fluorometer (Invitrogen). Subsequently, the libraries were delivered to Novogene for sequencing.

Two biological replicates were analyzed in parallel in this experiment, which generated three NGS raw datasets. After the production of clean data by demultiplexing and removing adaptor regions, pairs of paired-end data were merged by FLASH script^70^, and those reads without detected pairs were removed. Python scripts generated in house were then used to search for the ‘GGATN86ATGC’ 94-mer in the sequencing reads (and the reverse complementary sequence), and those carrying mutations within the upstream (GGAT) or downstream (ATGC) flanking regions (4 nt each) were removed. The read counts were then adjusted using Eq. 19, where *n* is the number of sequencing libraries, to normalize the different sequencing depths of each library.

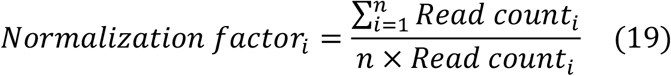

The library showed negligible variation in growth (**Supplementary Fig. 11**), which ensured the feasibility of using the library in subsequent Sort-Seq experiments.

### Sort-Seq experiments

For both the promoter library and the combination library, a frozen glycerol stock of library cells (*E. coli* MG1655) was inoculated into 100-mL flasks containing 20 mL of LB medium with kanamycin to an initial OD_600_ of 0.02. Library cells were grown for 16 h at 37°C and 220 rpm. The grown cells were transferred to fresh LB medium to an initial OD_600_ of 0.02 and grown again under the same conditions as above. A third round of dilution and growth was carried out to improve the expression stability of the fluorescent proteins. After growth, 500 μL of culture medium was chilled on ice immediately, and the cells were collected by centrifugation (4°C; 8,000 × g for 10 min). The cells were resuspended in 500 μL of prechilled PBS. Each cell suspension was diluted 150-fold in PBS to prepare samples appropriate for sorting. Three biological replicates were prepared for Sort-Seq experiments.

Sorting was performed on a FACSAria SORP (BD Biosciences). Gating based on FSC-Area and SSC-Area was carried out to exclude noncell particles. The population in this gated area is referred to as P1. The fluorescence background noise for the two relevant wavelengths was calibrated using the blank untransfected MG1655 strain. Note that the blank strain was completely negative for both sfGFP and mCherry expression. The resulting double-positive area in the region corresponding to the FITC-Area (sfGFP) and the PE-Texas Red-Area (mCherry) is referred to as Q2 after P1. The prepared library cells were analyzed by cytometry to determine the density distribution contour of the fluorescence in Q2. Subsequently, in the histogram of sfGFP/mCherry, 12 bins were set to evenly split the overall distribution of the population in Q2 (**Supplementary Figs. 13 and 19**, **Supplementary Tables 4 and 5**), referred to as P2 to P13 after Q2, to ensure that the number of cells in each bin was equal and improve the sorting efficiency. For calibration, ~2 × 10^6^ unsorted cells in gate Q2 were first collected for each sample. In the main sorting process, the three replicates were individually sorted into the 12 bins as described above. Each sample was successively sorted three times using four-way sorting. In each of these sorting runs, cells falling in nonadjacent bins were collected to eliminate the conflicting events between them. Thus, P2, P5, P8 and P11 were simultaneously sorted in one run, as were P3, P6, P9 and P12. During sorting, the cell flow rate was kept at ~8000 events/s, and ~5 × 10^5^ cells were collected in each bin.

The sorted cells were collected in 36 (3 samples × 12 bins) 5-mL polystyrene round-bottom centrifuge tubes (BD Falcon), each of which contained 500 μL of PBS. The entire contents of each tube were then each transferred to 100-mL flasks containing 20 mL of LB medium with kanamycin and cultured at 37°C for 7 h. These cells were then subjected to plasmid library extraction. Together with the cells from gate Q2 of each of the three samples mentioned above, we obtained 39 plasmid libraries in total.

The promoter and RBS regions of *sfgfp* in each library were amplified through PCR (KAPA HiFi PCR Kit; 95°C for 3 min, 25 cycles [98°C for 20 s, 63°C for 15 s, 72°C for 5 s], 72°C for 30 s), using 12 8-nt barcoded primers to identify different sorting bins (primers sorting_P2 to _P13, **Supplementary Table 2**). The barcodes were designed according to the following principles. (1) The Levenshtein distance between every two barcodes was ≥ 4; (2) the GC content was 20% to 80%; and (3) there were no more than four consecutive identical bases. In a 25-μL PCR, 5 ng of plasmid library was added as a template. PCR products from the 12 bins for each sample were mixed, thus obtaining three sorted PCR products (from sorted library cells in different sorting bins for three samples) and three unsorted PCR products (from unsorted library cells in the Q2 gate for three samples). The resulting PCR products were analyzed and purified by electrophoresis. The sequencing libraries were prepared as described above and were then delivered to Novogene for sequencing.

### Sort-Seq data processing

According to NGS data, the read count *R_i,k_* for *Variant_i_* in *bin_k_* can be observed. Additionally, by analyzing the cytometry data, we can obtain the ratio of cells sorted into *bin_k_* against all cells, which is denoted by *C_k_*. Hence, assuming an unbiased NGS quantification process, the proportion of *Variant_i_* in *bin_k_* is *Q_ik_* = *R_i,k_*/*R_k_*. Here, *R_k_* is the total read count for the NGS library derived from *bin_k_*. Therefore, the probability of sorting *Variant_i_* into *bin_k_*) should be 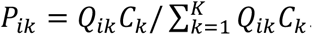.

For the library of *tnaC* variants and the malonyl-CoA biosensors, data processing was performed as described above. However, for the promoter library, a sorting error did exist (**Supplementary Figs. 14a, 14b, 14d and 14f**). We ascribed this error to random screening with an error rate, *ε*, and hence, we modified *P_i,k_* as shown in Eq. 20.

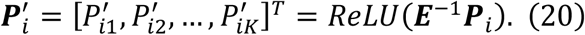

where

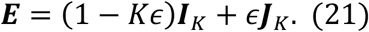

Here, ***I**_K_* is an identity matrix of size *K*, and ***J**_K_* is a matrix of ones of size *K*. Finally, the binned distribution was obtained by 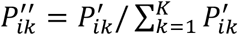. For both the promoter and combination libraries, ε was set to 0.05, which made the binned distribution more precise (**Supplementary Figs. 14c, 14e and 14g**). The strains with 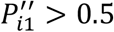 or 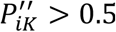 were ruled out, as they were not effectively sorted. Moreover, to ensure the quality of the results, we eliminated the data with low consistency among replicates. Specifically, if the CV of the calculated mean or SD among biological replicates was greater than 0.5, the related data were removed from the dataset.

### Evaluation of the expression patterns of essential and nonessential genes

The essential genes were identified based on a comprehensive pooled CRISPRi screening dataset^71^ (threshold: fitness ≤ −6), whereas other genes were regarded as nonessential. We calculated the transcriptional strength of each gene. Specifically, the genes belonging to the operons closest to the downstream side of a promoter were considered to be driven by this promoter, and the transcription strength of a gene was calculated as the summation of its promoter strengths.

### Identification of RBS strengths

To identify the translational strengths of the 13 RBSs, we defined each promoter strength as the expression strength in the promoter library, and then we divided each expression strength from the combination library by the corresponding promoter strength. The median of the calculated results for combinations with the same RBS is defined as the translational strength of that RBS. The resulting order of the RBS strengths was as follows: apFAB872(0.15) < apFAB914(0.23) < apFAB864(0.36) < apFAB865(0.67) < apFAB927(0.75) < apFAB827(0.95) < apFAB894(1.02) < apFAB909(1.04) < apFAB839(1.36) < apFAB833(1.38) < apFAB834(1.41) < apFAB820(1.57) < apFAB916(2.13) (**Supplementary Fig. 23**).

### Grouping promoters according to expression noise

Each noise-mean relationship was fitted by the empirical formula *CV*^2^ = *C*_1_ + *C*_2_ /*Mean* (**Fig. 5a and 5b**), and the residual of each expression pattern was calculated and sorted. In addition, to ensure the reliability of the analysis, we only considered the sequences with appropriate mean expression levels (> 0.15 for the promoter library; > 0.1 for the combination library). The 20 candidates from the promoter library and 25 from the combination library with the largest residues were reconstructed. For the promoter library, the 10% with the largest residuals was classified as the high-noise group, the 10% with the smallest residuals was classified as the low-noise group, and the remaining sequences were grouped as medium-noise.

### Identification of potential RpoD-binding sites

The DPinteract database contains computational predictions of possible RpoD-binding sites with 15-19 nucleotide spacing in the *E. coli* genome^45^. We searched these sequences in the promoter library and counted the number of potential RpoD-binding sites of each promoter. To avoid redundant results, we only accounted for independent hexamer pairs. Specifically, if the −35 or −10 region of two RpoD-binding sites overlapped, we only considered the RpoD-binding site with a higher z score; otherwise, both of them were retained (**Supplementary Fig. 31**).

## DATA AVAILABILITY

Raw NGS data of Sort-Seq have been deposited into the NCBI Short Read Archive with BioProject accession number PRJNA800535. The plasmid maps related to this work can be accessed via Github (https://github.com/fenghuibao/dSort-Seq). The dSort-Seq calculation tool can be accessed through our laboratory website (http://www.thu-big.net/dsort-seq/).

## ACKNOWLEDGMENTS

We would like to thank Drs. X. Zheng and Y. Zhou for kindly sharing their experimental skills and data. We thank Dr. Y. Wu for construction of the pMPTPV plasmid. We thank B. Yu for her help with the FACS experiments and F. Liu for her help with NGS library construction. We thank Prof. F. Zhang (Washington University in St. Louis) for critical discussions regarding this work. This work was supported by the National Key Research and Development Program of China (2019YFA0904800), the National Natural Science Foundation of China (U2032210), and the Foshan-Tsinghua Innovation Special Fund (THFS01).

## Author Contributions

H.F. conceived the general framework of this work, including the model construction and experimental design. H.F. and F.L. prepared the promoter library and performed the raw Sort-Seq experiment and the subsequent validation experiment. F.L. did the experiment for the combination library. H.F. analyzed all experimental results. H.F., F.L. and C.Z wrote the manuscript based on discussions and contributions of all authors. T.W. provided valuable opinions for the work and helped to revise the manuscript. C.Z., A.Z. and X.X. supervised the project.

## CONFLICT OF INTEREST

The authors declare that there is no conflict of interest regarding the publication of this article.

## Notes

### Competing Interest Statement

The authors have declared no competing interest.

