## supplementary_materials for "Deep-learning-assisted Sort-Seq enables high-throughput profiling of gene expression characteristics with high precision"

**Algorithm 1** Parameter learning model of dSort-Seq. The default values are set as $t = 8,\ m = 640,\ \alpha_1 = 0.99,$ $\alpha_2 = 0.01$.

**for** number of training iterations **do**

**for** $t$ steps **do**

- Sample minibatch of $m$ samples $\{x^{(1)}_{fake}, x^{(2)}_{fake}, ..., x^{(m)}_{fake}\}$ from $f(x|\boldsymbol{\pi}, \boldsymbol{\lambda}, \boldsymbol{\mu}, \boldsymbol{\sigma})$.
- Sample minibatch of $m$ samples $\{x^{(1)}_{true}, x^{(2)}_{true}, ..., x^{(m)}_{true}\}$ from real fluorescence intensity distribution.
- Update the discriminator by ascending its stochastic gradient:

$$\nabla_{\theta_d} \frac{1}{m} \sum_{j=1}^{m} [log(D(x^j_{true})) + log(1 - D(x^j_{fake}))].$$

**end for**

- Sample minibatch of $m$ samples $\{x^{(1)}_{fake}, x^{(2)}_{fake}, ..., x^{(m)}_{fake}\}$ from $f(x|\boldsymbol{\pi}, \boldsymbol{\lambda}, \boldsymbol{\mu}, \boldsymbol{\sigma})$.
- Update the generator by ascending its stochastic gradient:

$$\nabla_{\boldsymbol{\lambda},\boldsymbol{\mu},\boldsymbol{\sigma}} \Bigg[ \alpha_1 \frac{1}{n} \sum_{i=1}^{n} \sum_{k=1}^{K} \Bigg( P_{ik} \log \Bigg( \lambda_i \int_{b_{k-1}}^{b_k} N(x_i|\mu_{1i}, \sigma^2_{1i}) dx_i + (1 - \lambda_i) \int_{b_{k-1}}^{b_k} N(x_i|\mu_{2i}, \sigma^2_{2i}) dx_i \Bigg) \Bigg) + \alpha_2 \frac{1}{m} \sum_{j=1}^{m} (log(D(x^j_{fake}))) \Bigg]$$

**end for**

**Supplementary Figure 1** The parameter estimation algorithm of dSort-Seq

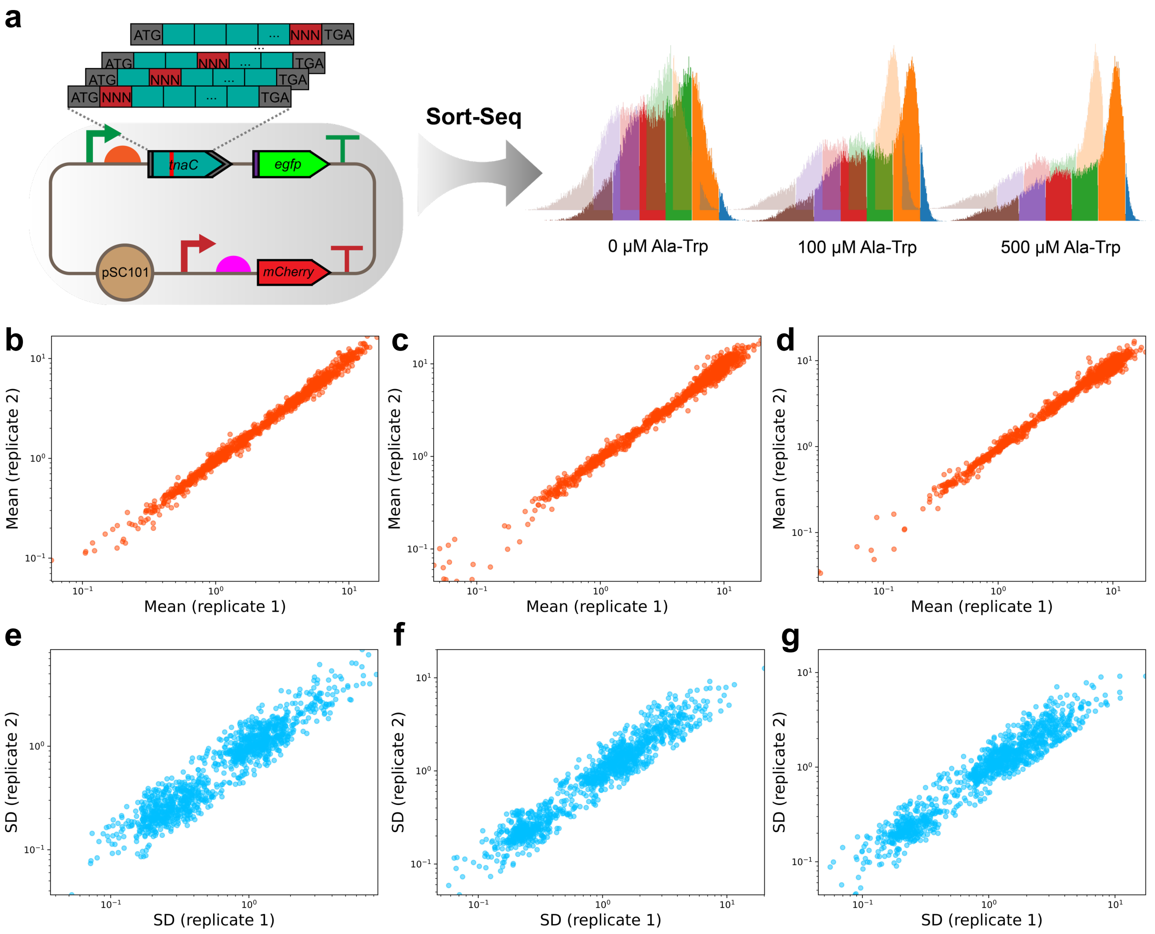


**Supplementary Figure 2** The dSort-Seq profiling of *tnaC*-mediated tryptophan-dependent gene expression. (**a**) Sort-Seq characterization of the *tnaC* variant library under 3 different ligand concentrations (0, 100 and 500 μM Ala-Trp). Cells were sorted into 6 bins according to their responses to ligand. Two biological replicates were examined for each Sort-Seq experiment. (**b-d**) The mean expression revealed strong correlations between replicates (**b,** 0 μM Ala-Trp, $n$ = 1,267; **c,** 100 μM Ala-Trp, $n$ = 1,244; **d,** 500 μM Ala-Trp, $n$ = 1,221). (**e-g**) The standard deviation (SD) of expression revealed high consistency between replicates (**e,** 0 μM Ala-Trp; **f,** 100 μM Ala-Trp; **g,** 500 μM Ala-Trp).

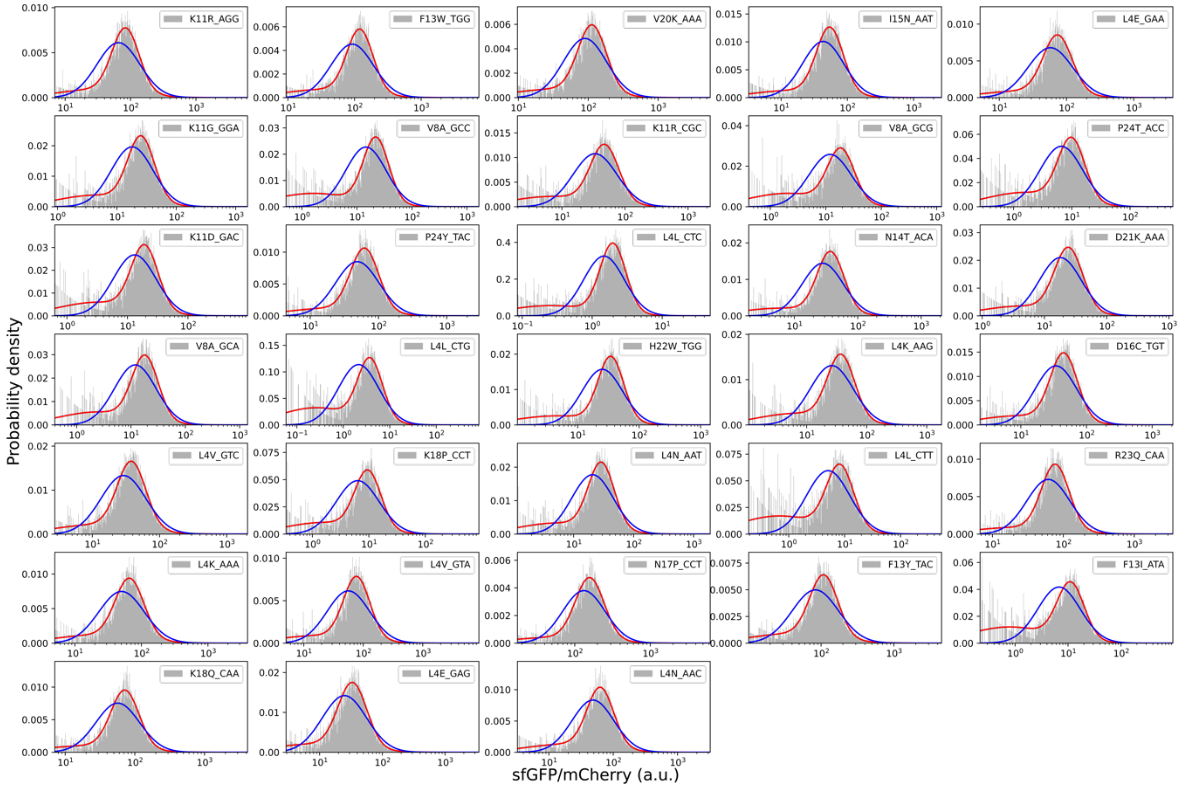


**Supplementary Figure 3** Individual cytometry assay of fluorescence intensity distributions of 33 *tnaC* variants under 0 μM Ala-Trp. Each distribution was fitted with a two-component LGMM (indicated by the red line) and a log-normal distribution (indicated by the blue line).

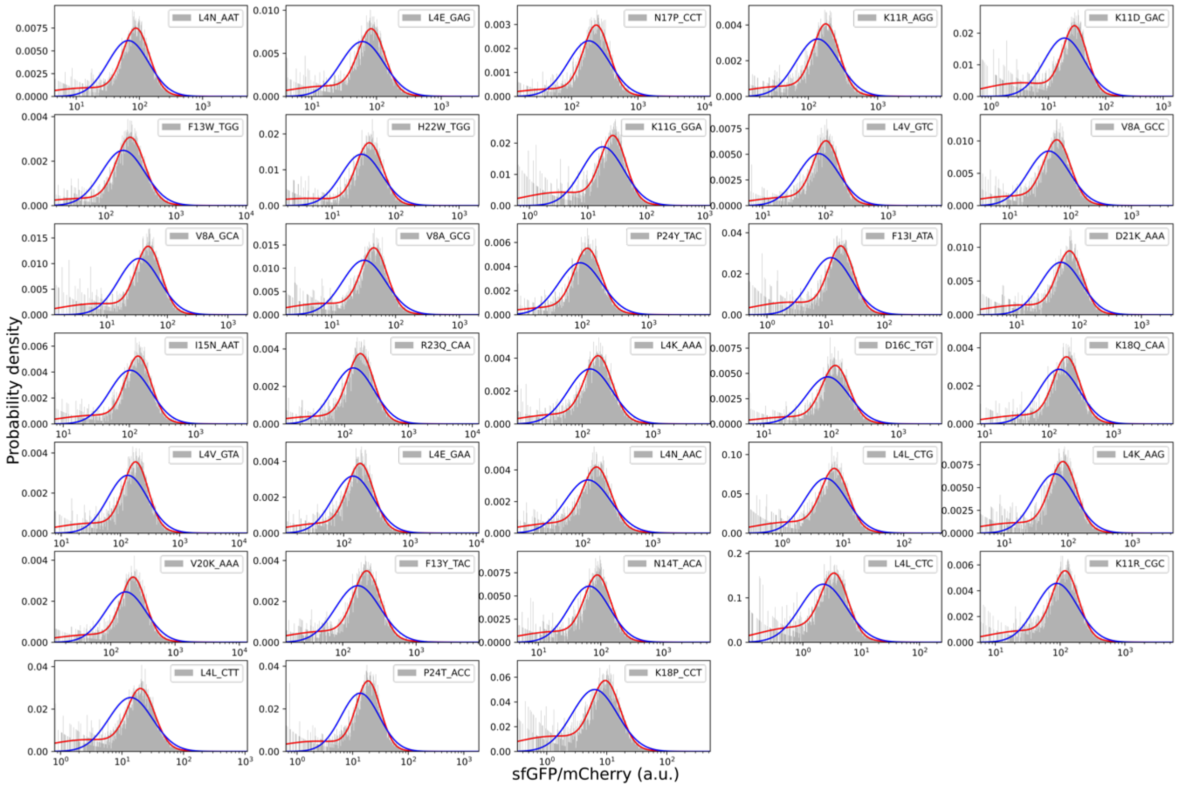


**Supplementary Figure 4** Individual cytometry assay of fluorescence intensity distributions of 33 *tnaC* variants under 100 μM Ala-Trp. Each distribution was fitted with a two-component LGMM (indicated by the red line) and a log-normal distribution (indicated by the blue line).

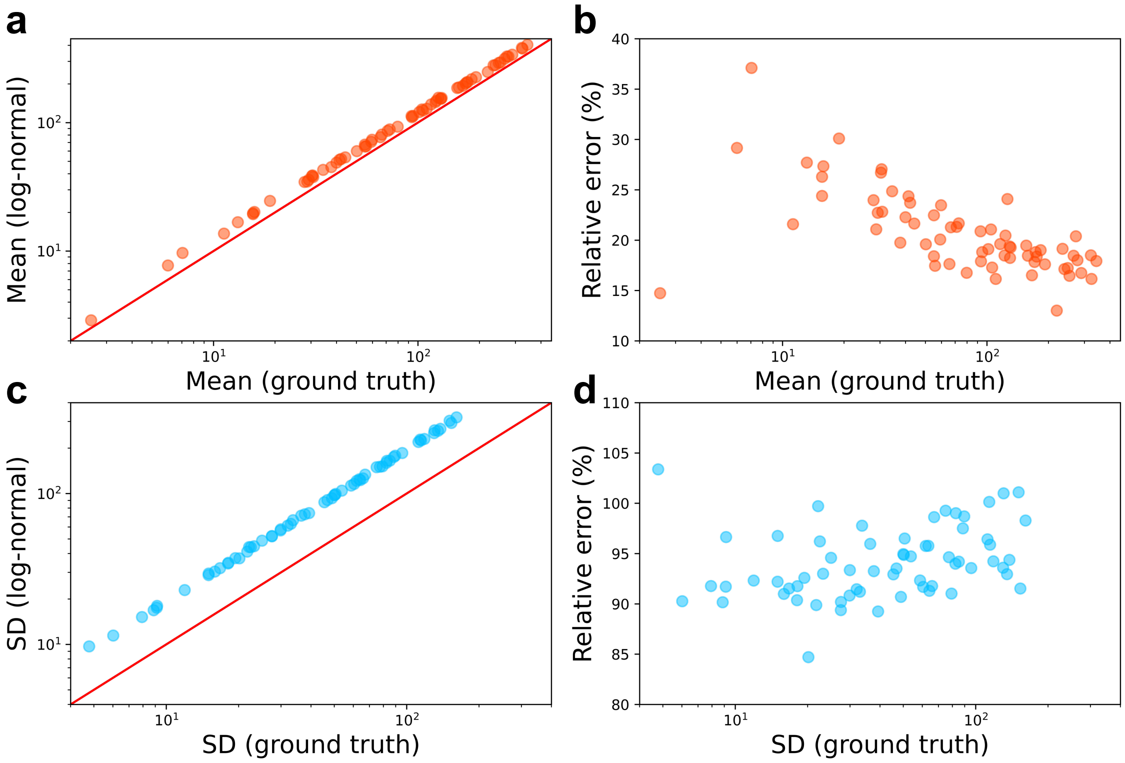


**Supplementary Figure 5** The response and SD of *tnaC* variants inferred from the log-normal-based method are subject to error. (**a**) Comparison of log-normal derived expression strength and ground truth (red line, calculated with LGMM). (**b**) The relationship between relative error and expression strength. The error is greater at lower expression levels. (**c**) Comparison of log-normal derived SD and ground truth (red line, calculated with LGMM). (**d**) The log-normal-based method shows high relative error in the calculation of SD.

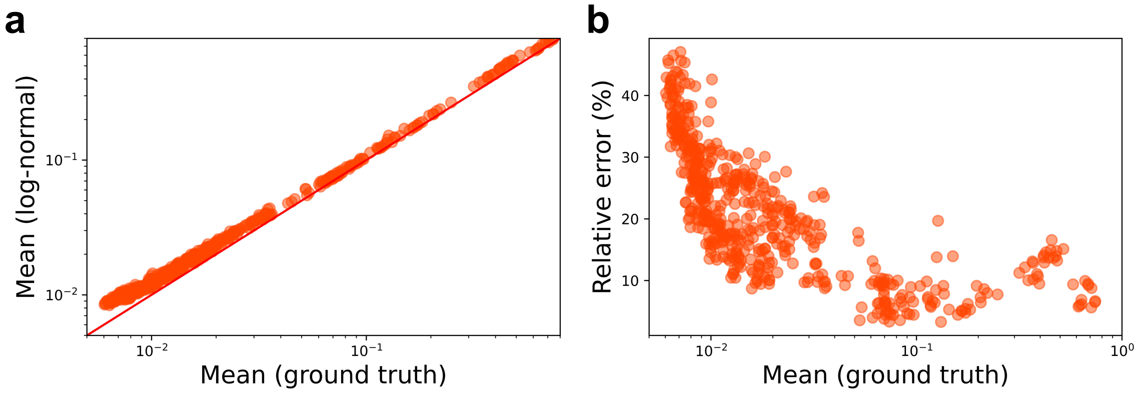


**Supplementary Figure 6** The responses of malonyl-CoA biosensors inferred from the log-normal distribution were subjected to error. (**a**) Comparison of log-normal derived mean expression levels and ground truth (red line, calculated with LGMM). Generally, log-normal would overestimate the expression strength. (**b**) The relationship between relative error and expression strength. The error is greater at lower expression levels.

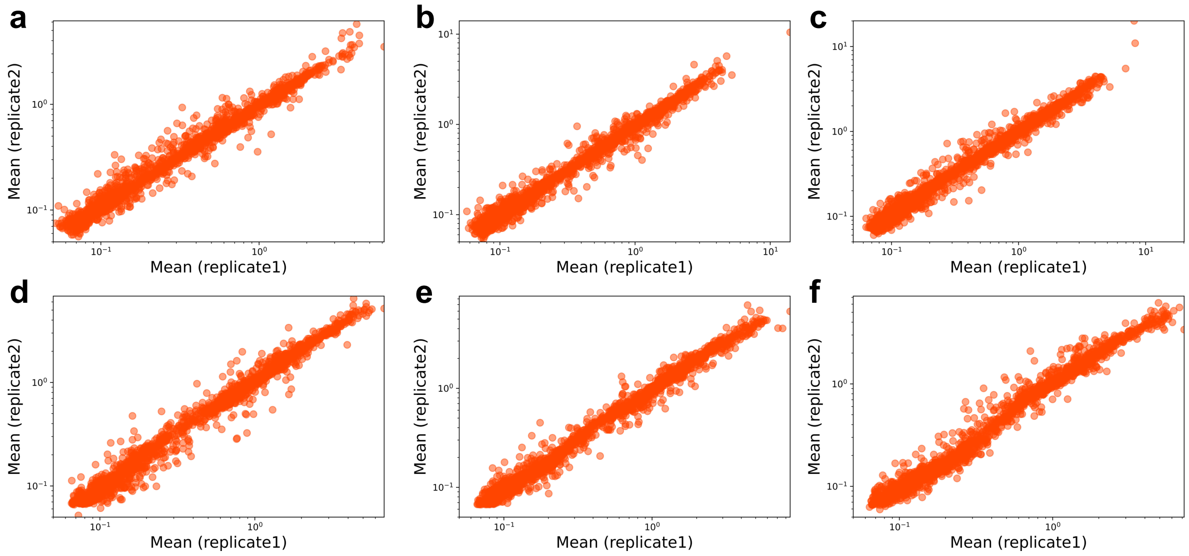


**Supplementary Figure 7** The expression strengths (mean values of YPet/mCherry) showed strong correlations between biological replicates at different concentrations (**a**, 0 mg/L cerulenin, *n* = 1,937, Pearson's *r* = 0.978; **b**, 1 mg/L cerulenin, *n* = 2,091, Pearson's *r* = 0.984; **c**, 2 mg/L cerulenin, *n* = 2,221, Pearson's *r* = 0.950; **d**, 3 mg/L cerulenin, *n* = 2,075, Pearson's *r* = 0.987; **e**, 5 mg/L cerulenin, *n* = 2,233, Pearson's *r* = 0.982; **f**, 8 mg/L cerulenin, *n* = 2,222, Pearson's *r* = 0.979).

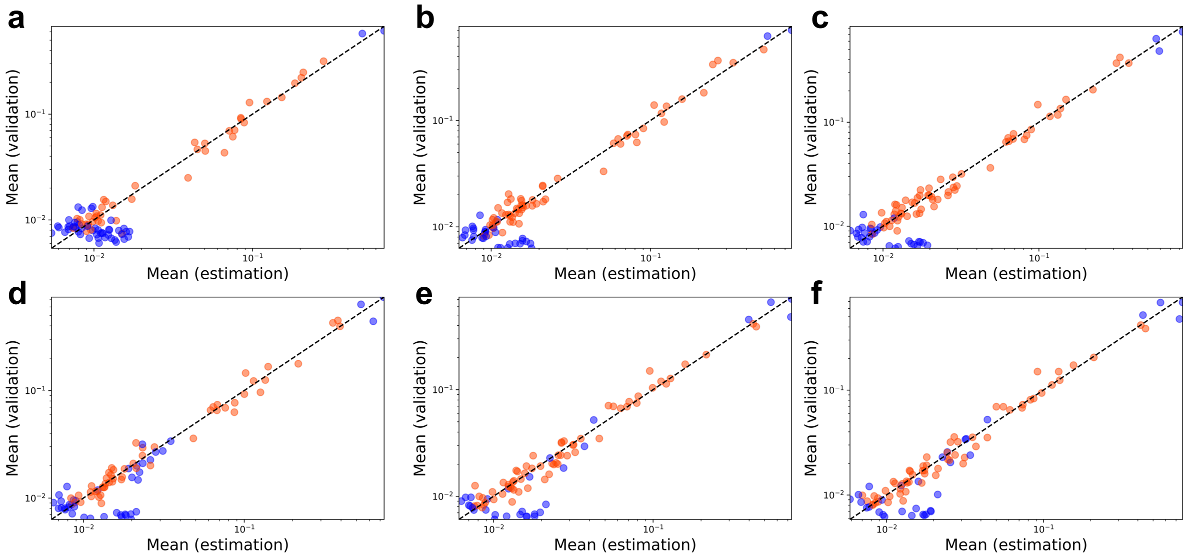


**Supplementary Figure 8** **(a-f)** Individually analyzed responses of reconstructed malonyl-CoA biosensors were highly correlated with those estimated from Sort-Seq (indicated by red dots) and the machine learning approach (indicated by blue dots) at different concentrations (**a**, 0 mg/L cerulenin, Pearson's $r$ = 0.990; **b**, 1 mg/L cerulenin, Pearson's $r$ = 0.987; **c**, 2 mg/L cerulenin, Pearson's $r$ = 0.987; **d**, 3 mg/L cerulenin, Pearson's $r$ = 0.979; **e**, 5 mg/L cerulenin, Pearson's $r$ = 0.975; **f**, 8 mg/L cerulenin, Pearson's $r$ = 0.970). Linear regression was applied to fit the data values within the same scale.

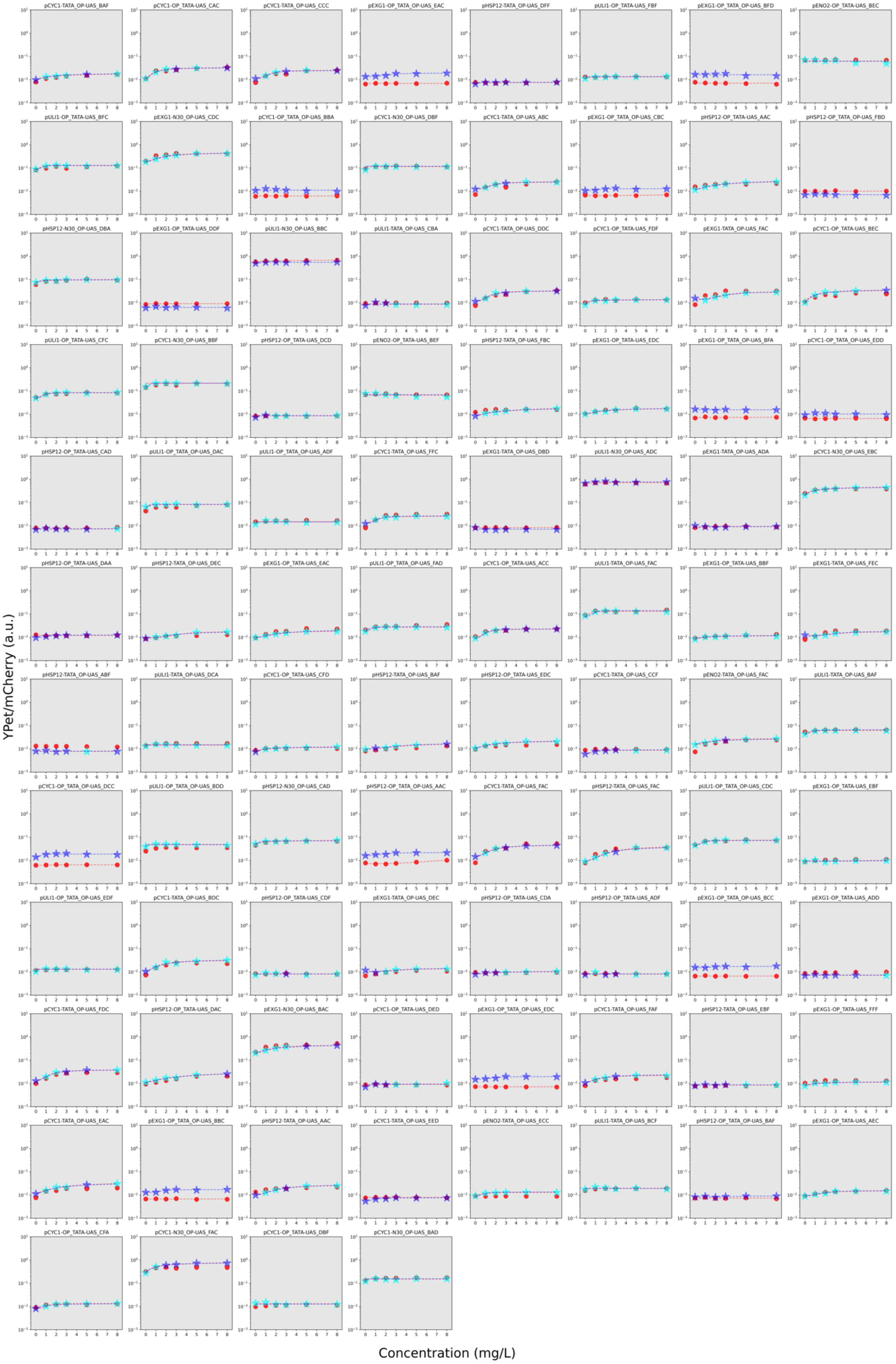

pCYC1-TATA_OP-UAS_BAF
pCYC1-OP_TATA-UAS_CAC
pCYC1-TATA_OP-UAS_CCC
pEXG1-OP_TATA_OP-UAS_EAC
pHSP12-TATA_OP-UAS_DFF
pULI1-OP_TATA-UAS_FBF
pEXG1-OP_TATA_OP-UAS_BFD
pENO2-OP_TATA-UAS_BEC
pULI1-OP_TATA-UAS_BFC
pEXG1-N30_OP-UAS_CDC
pCYC1-OP_TATA_OP-UAS_BBA
pCYC1-N30_OP-UAS_DBF
pCYC1-TATA_OP-UAS_ABC
pEXG1-OP_TATA_OP-UAS_CBC
pHSP12-OP_TATA-UAS_AAC
pHSP12-OP_TATA_OP-UAS_FBD
pHSP12-N30_OP-UAS_DBA
pEXG1-OP_TATA-UAS_DDF
pULI1-N30_OP-UAS_BBC
pULI1-OP_TATA-UAS_CBA
pCYC1-TATA_OP-UAS_DDC
pCYC1-OP_TATA-UAS_FDF
pEXG1-TATA_OP-UAS_FAC
pCYC1-OP_TATA-UAS_BEC
pULI1-OP_TATA-UAS_CFC
pCYC1-N30_OP-UAS_BBF
pHSP12-OP_TATA-UAS_DCD
pENO2-OP_TATA-UAS_BEF
pHSP12-TATA_OP-UAS_FBC
pEXG1-OP_TATA-UAS_EDC
pEXG1-OP_TATA_OP-UAS_BFA
pCYC1-OP_TATA_OP-UAS_EDD
pHSP12-OP_TATA-UAS_CAD
pULI1-OP_TATA_OP-UAS_DAC
pULI1-OP_TATA_OP-UAS_ADF
pCYC1-TATA_OP-UAS_FFC
pEXG1-TATA_OP-UAS_DBD
pULI1-N30_OP-UAS_ADC
pEXG1-TATA_OP-UAS_ADA
pCYC1-N30_OP-UAS_EBC
pHSP12-OP_TATA-UAS_DAA
pHSP12-TATA_OP-UAS_DEC
pEXG1-OP_TATA-UAS_EAC
pULI1-OP_TATA_OP-UAS_FAD
pCYC1-OP_TATA-UAS_ACC
pULI1-TATA_OP-UAS_FAC
pEXG1-OP_TATA-UAS_BBF
pEXG1-TATA_OP-UAS_FEC
pHSP12-TATA_OP-UAS_ABF
pULI1-TATA_OP-UAS_DCA
pCYC1-OP_TATA-UAS_CFD
pHSP12-TATA_OP-UAS_BAF
pHSP12-OP_TATA-UAS_EDC
pCYC1-OP_TATA-UAS_CCF
pENO2-TATA_OP-UAS_FAC
pULI1-TATA_OP-UAS_BAF
pCYC1-OP_TATA_OP-UAS_DCC
pULI1-OP_TATA_OP-UAS_BDD
pHSP12-N30_OP-UAS_CAD
pHSP12-OP_TATA_OP-UAS_AAC
pCYC1-TATA_OP-UAS_FAC
pHSP12-TATA_OP-UAS_FAC
pULI1-OP_TATA_OP-UAS_CDC
pEXG1-OP_TATA-UAS_EBF
pULI1-OP_TATA_OP-UAS_EDF
pCYC1-TATA_OP-UAS_BDC
pHSP12-OP_TATA-UAS_CDF
pEXG1-TATA_OP-UAS_DEC
pHSP12-OP_TATA-UAS_CDA
pHSP12-OP_TATA-UAS_ADF
pEXG1-OP_TATA_OP-UAS_BCC
pEXG1-OP_TATA-UAS_ADD
pCYC1-TATA_OP-UAS_FDC
pHSP12-OP_TATA-UAS_DAC
pEXG1-N30_OP-UAS_BAC
pCYC1-OP_TATA-UAS_DED
pEXG1-OP_TATA_OP-UAS_EDC
pCYC1-TATA_OP-UAS_FAF
pHSP12-TATA_OP-UAS_EBF
pEXG1-OP_TATA-UAS_FFF
pCYC1-TATA_OP-UAS_EAC
pEXG1-OP_TATA_OP-UAS_BBC
pHSP12-TATA_OP-UAS_AAC
pCYC1-TATA_OP-UAS_EED
pENO2-TATA_OP-UAS_ECC
pULI1-TATA_OP-UAS_BCF
pHSP12-OP_TATA_OP-UAS_BAF
pEXG1-OP_TATA-UAS_AEC
pCYC1-OP_TATA-UAS_CFA
pCYC1-N30_OP-UAS_FAC
pCYC1-OP_TATA-UAS_DBF
pCYC1-N30_OP-UAS_BAD
YPet/mCherry (a.u.)
Concentration (mg/L)

**Supplementary Figure 9** The dose–response relationships of 92 individually characterized malonyl-CoA biosensors are highly consistent with the dSort-Seq and machine learning results. All the data points represent the mean values of YPet/mCherry. Red dots represent individual characterization data, cyan stars represent data from dSort-Seq characterizations, and blue stars denote data from machine learning predictions. The dashed lines represent response curves fitted by the Hill equation (see **Methods**).

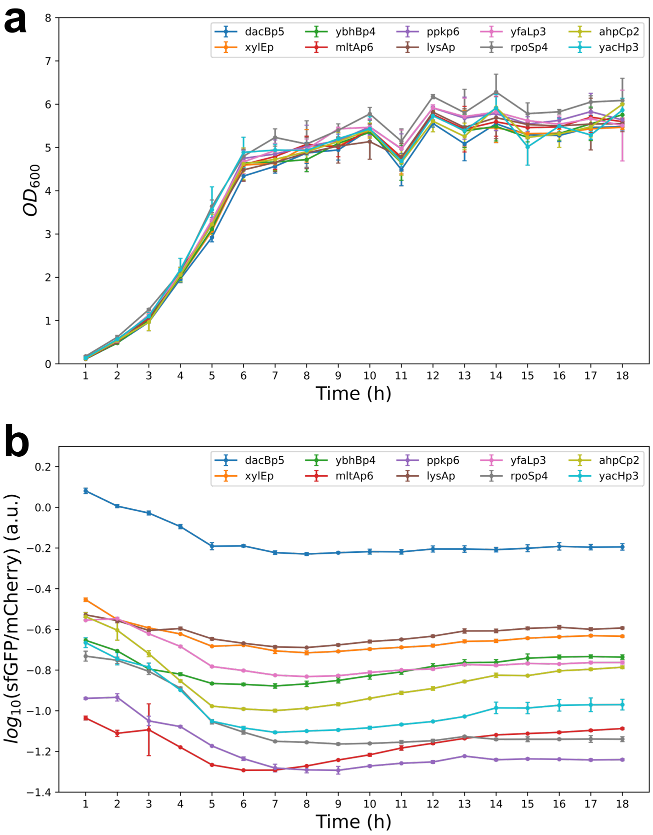


**Supplementary Figure 10** Feasibility verification of the pMPTPV_dual_fluorescence plasmid used for Sort-Seq. Ten strains carrying different promoters were constructed and characterized. (**a**) The growth curves of the 10 strains showed no apparent difference. (**b**) The median value of sfGFP/mCherry of the 10 strains remained stable after cultivation for 16 h.

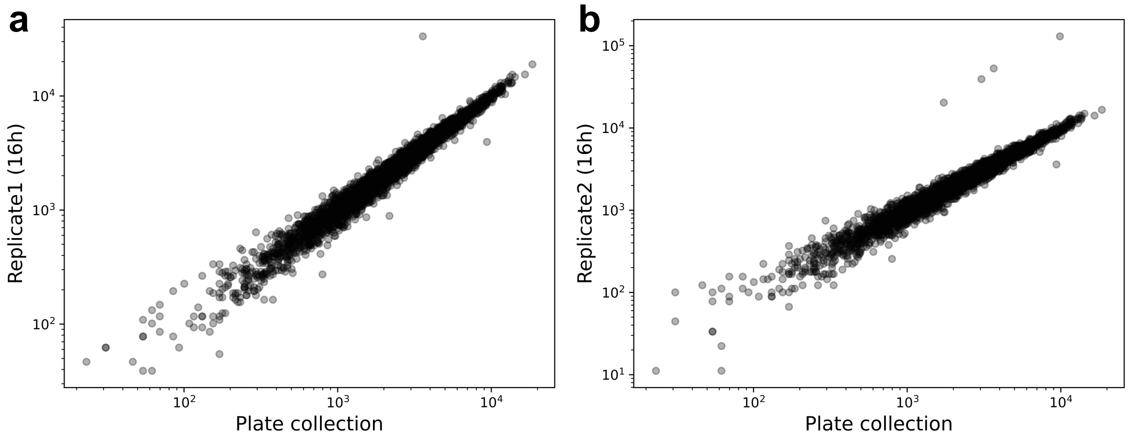


**Supplementary Figure 11** The promoter library showed negligible variation in growth (**a**, replicate 1, Pearson's $r$ = 0.985; **b**, replicate 2, Pearson's $r$ = 0.978). In **a** and **b**, the x-axis represents the NGS read count of the library collected from plates, and the y-axis represents the NGS read count after culturing in LB broth for 16 h.

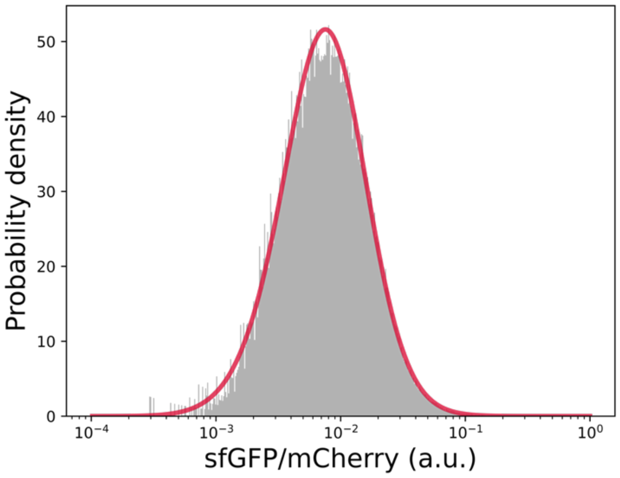


**Supplementary Figure 12** The autofluorescence intensity distribution of the library, which was measured by cytometry assay of the pMPTPV strain with only mCherry expression and no sfGFP expression. The mean (0.01611) and SD (0.00987) of autofluorescence were negligible relative to each candidate of the library.

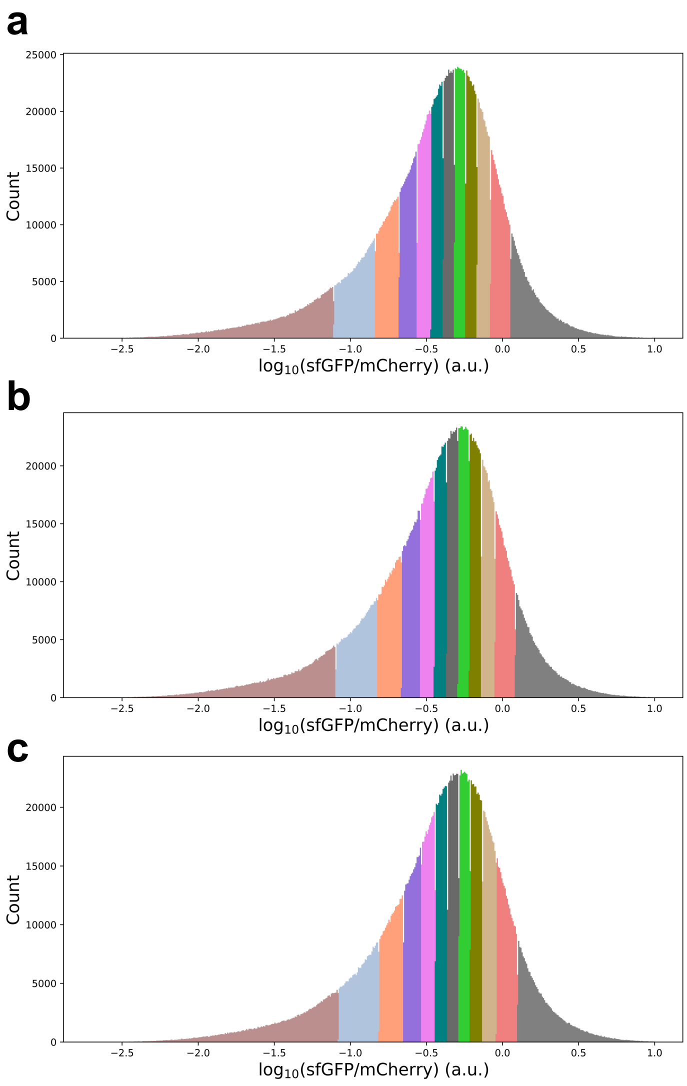


**Supplementary Figure 13** Overall fluorescence intensity distributions of the promoter library cells (**a, b** and **c** represent three biological replicates). For each sorting experiment, 12 bins were set to evenly split overall distribution, which displayed in different colors.

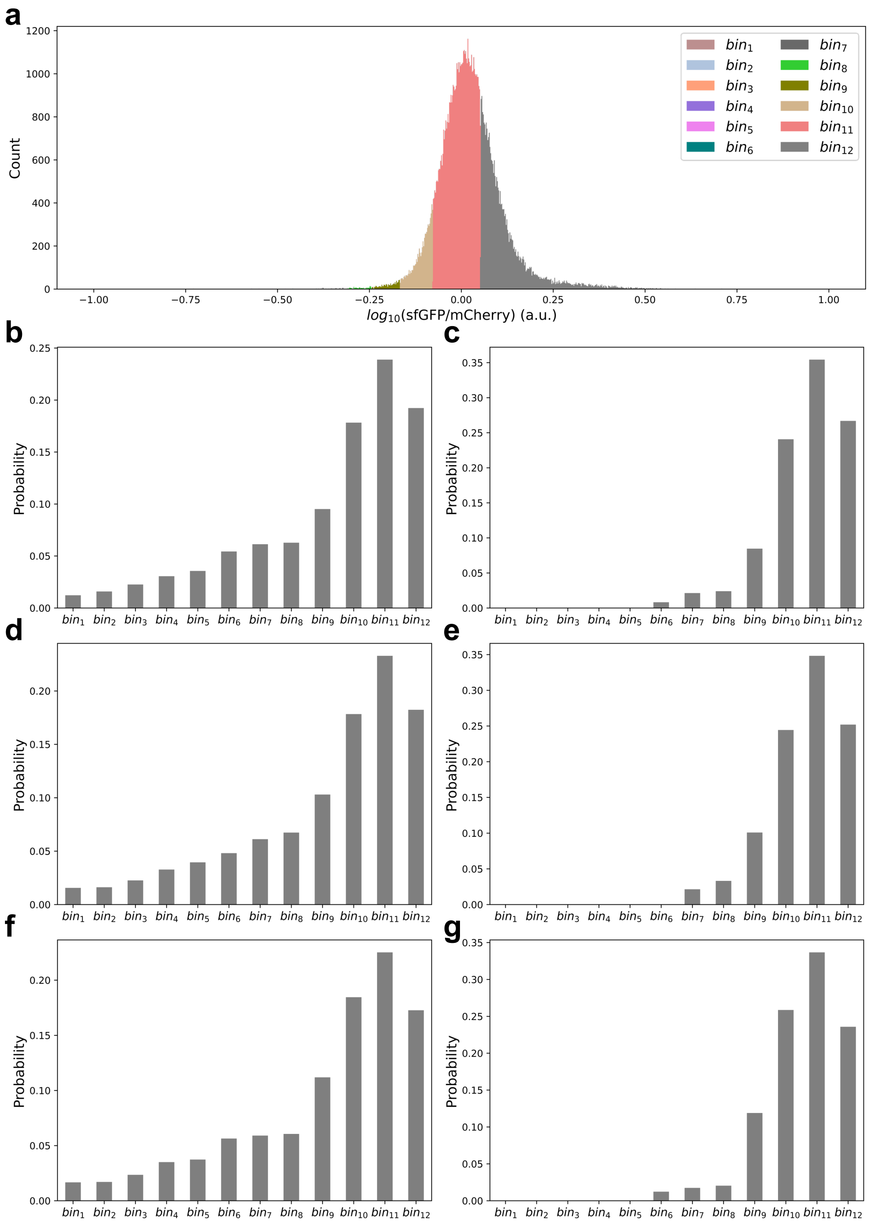


**Supplementary Figure 14** An example illustrating error elimination in Sort-Seq data processing. (**a**) Fluorescence intensity distribution of a variant in the promoter library. The sorting bins are displayed in different colors. (**b-g**), Binned distributions of the variant of three biological replicates before (**b**, **d**, and **f**) and after (**c**, **e**, and **g**) Sort-Seq data processing.

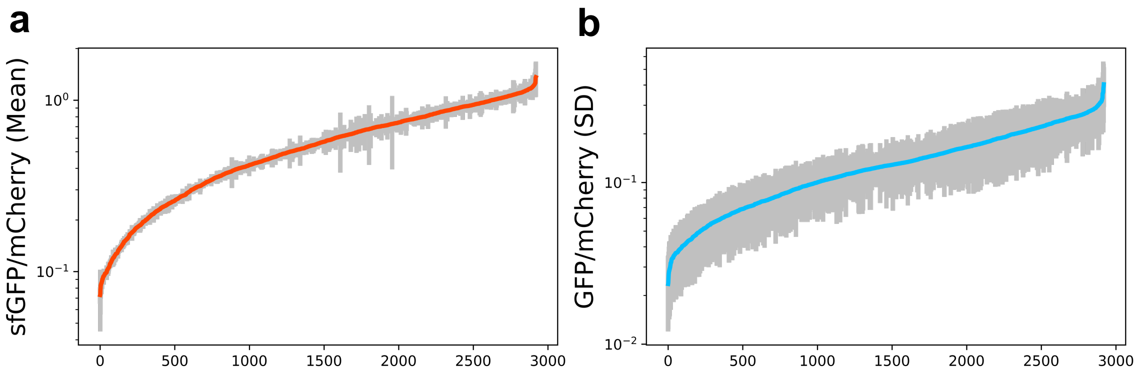


**Supplementary Figure 15** The expression strength (**a**) and the SD (**b**) of the promoter library calculated from dSort-Seq reveal strong correlations among three biological replicates ($n$ = 2,920). The data are presented as geometric mean values ± SDs.

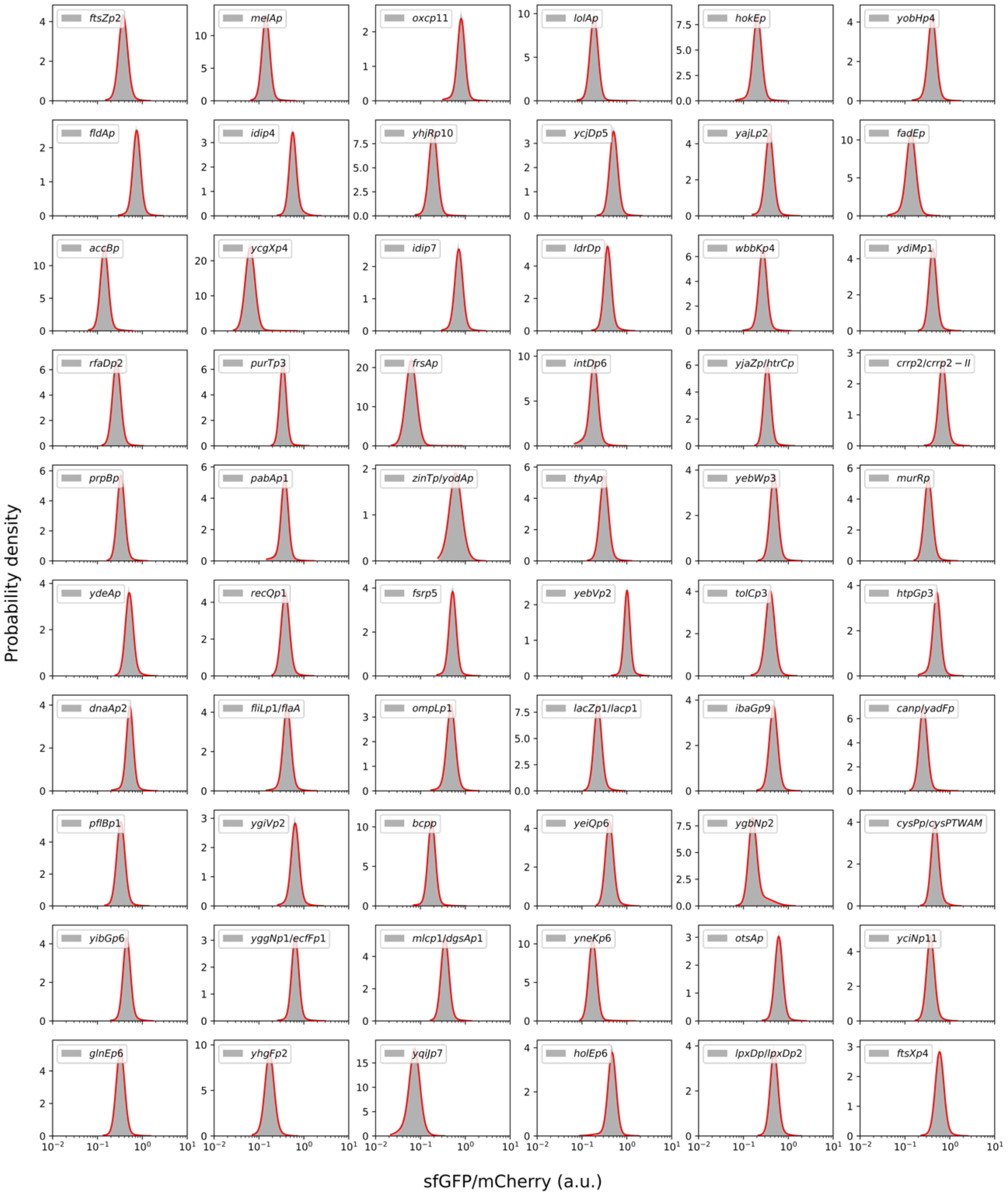


**Supplementary Figure 16** Individual cytometry assay of fluorescence intensity distributions of 60 randomly selected variants from the promoter library. Each distribution was fitted with a two-component LGMM.

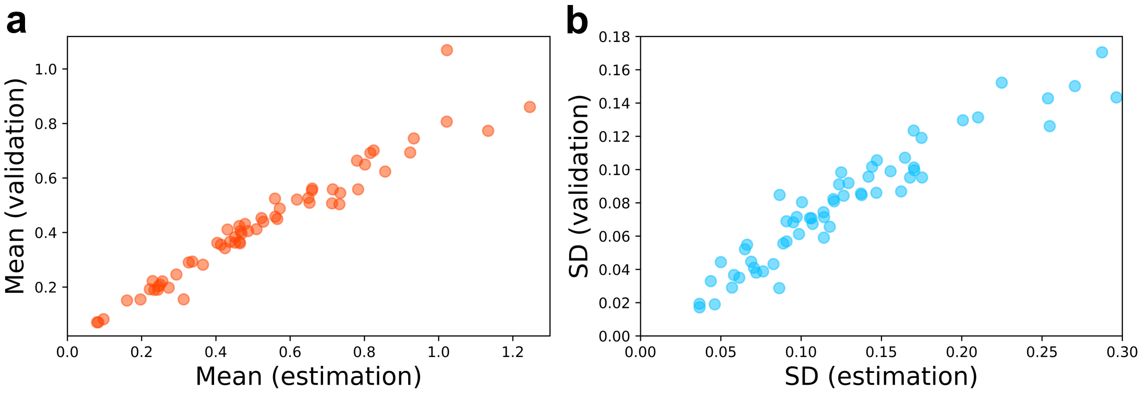


**Supplementary Figure 17** The expression properties estimated from dSort-Seq were validated with cytometry assays of individual colonies in terms of both the mean (**a,** $n = 60$, Pearson's $r = 0.981$) and SD (**b,** $n = 60$, Pearson's $r = 0.921$).

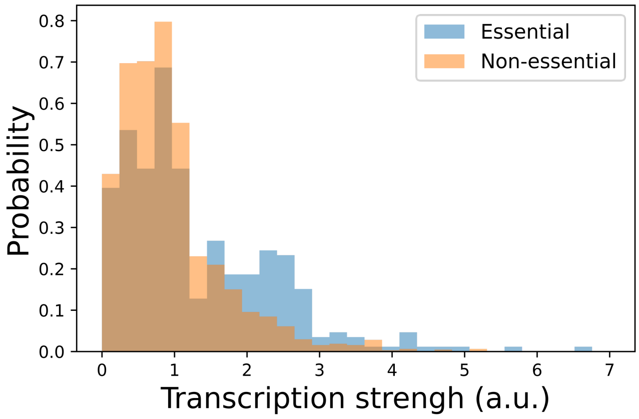


**Supplementary Figure 18** Comparison of transcriptional strengths between the essential and nonessential genes of *E. coli.*

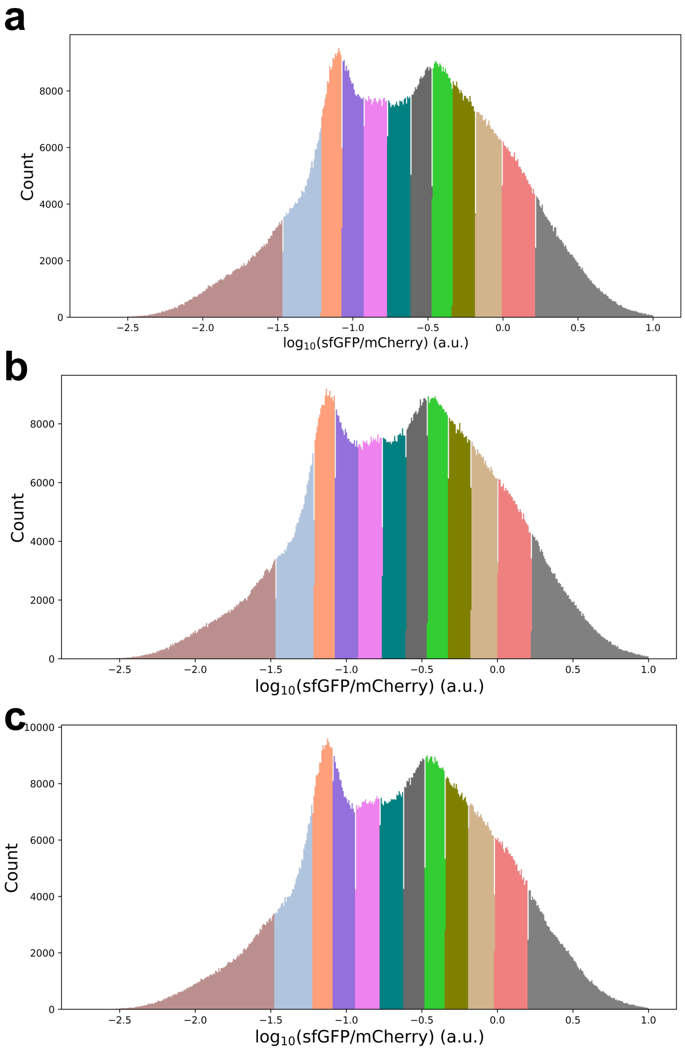


**Supplementary Figure 19** Overall fluorescence intensity distributions of the combined library cells (**a, b** and **c** represent three biological replicates). For each sorting experiment, 12 bins were set to evenly split overall distribution, which displayed in different colors.

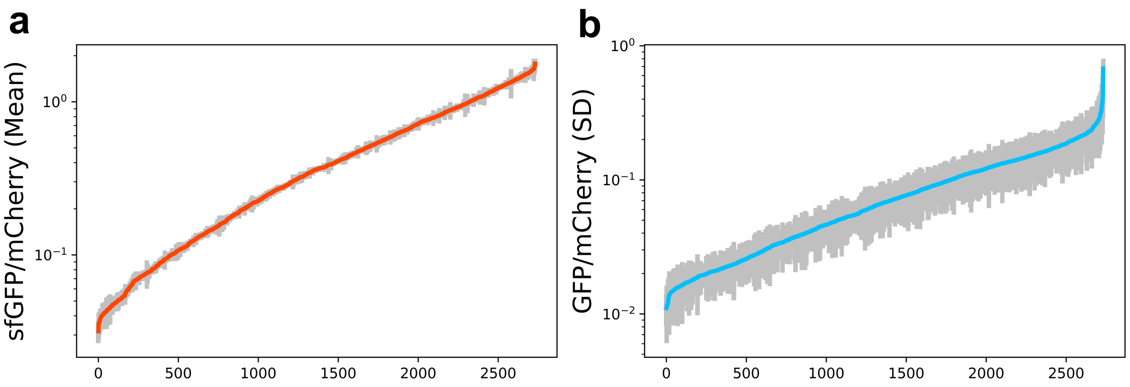


**Supplementary Figure 20** The expression strength (**a**) and the SD (**b**) of the combination library calculated from dSort-Seq reveal strong correlations among three biological replicates (*n* = 2,733). The data are presented as geometric mean values ± SDs.

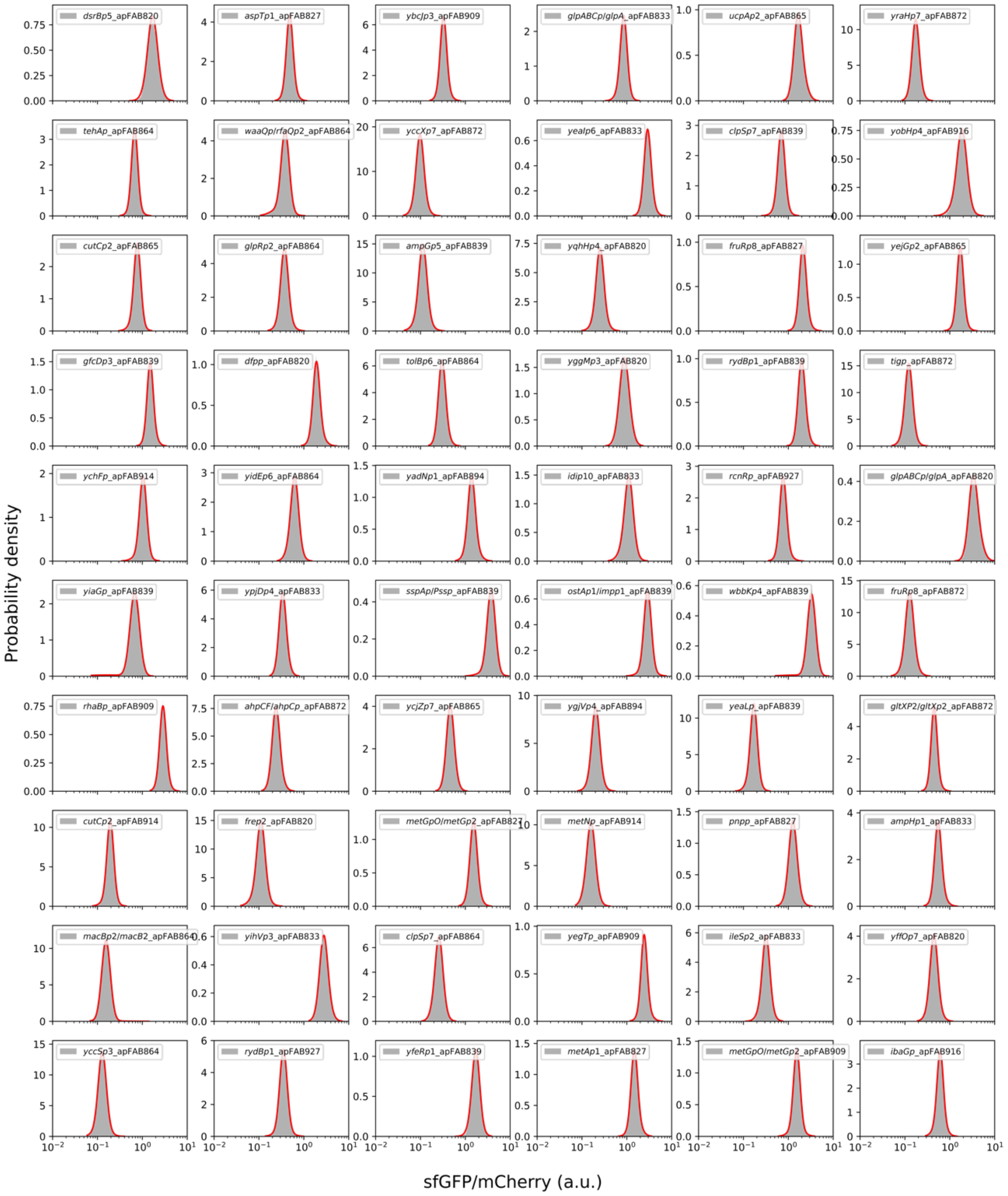


**Supplementary Figure 21** Individual cytometry assay of fluorescence intensity distributions of 60 randomly selected variants from the combination library. Each distribution was fitted with a two-component LGMM.

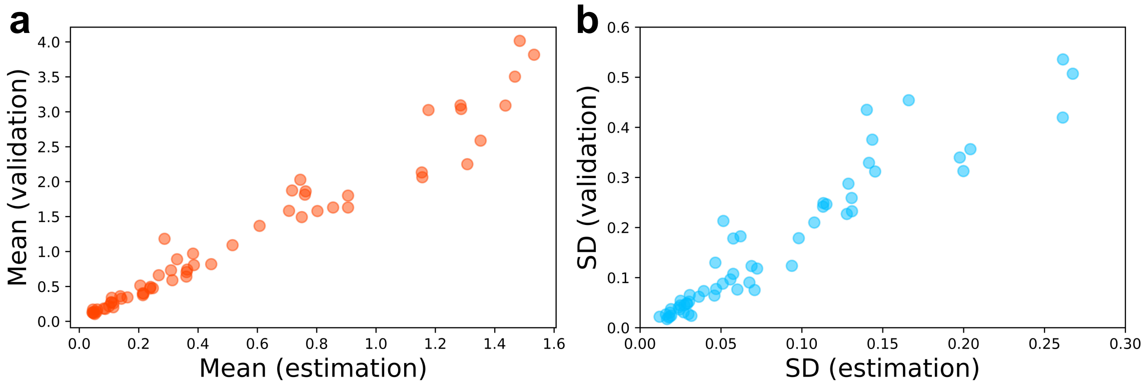


**Supplementary Figure 22** Individual validation for dSort-Seq profiling of the combination library. The expression properties estimated from dSort-Seq were validated with cytometry assays of individual colonies in terms of both the mean (**a,** $n$ = 60, Pearson's $r$ = 0.976) and SD (**b,** $n$ = 60, Pearson's $r$ = 0.937).

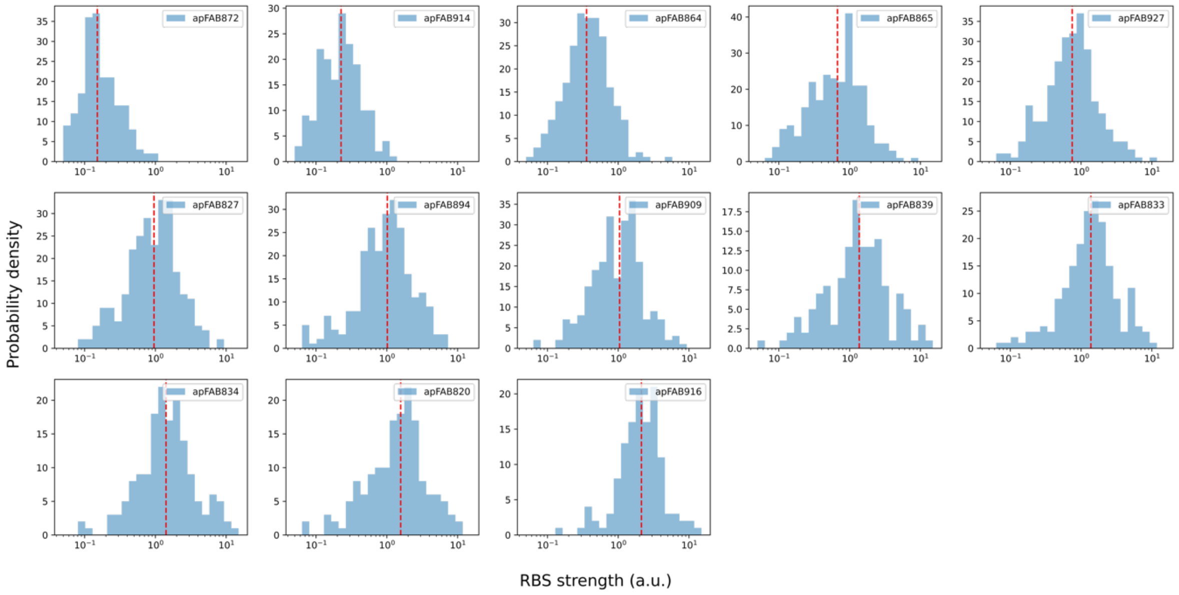


**Supplementary Figure 23** The distribution of RBS intensity, which was calculated by dividing each expression strength of the combination library by the corresponding promoter strength. The median of each distribution (indicated by the red dashed line) was defined as the RBS strength.

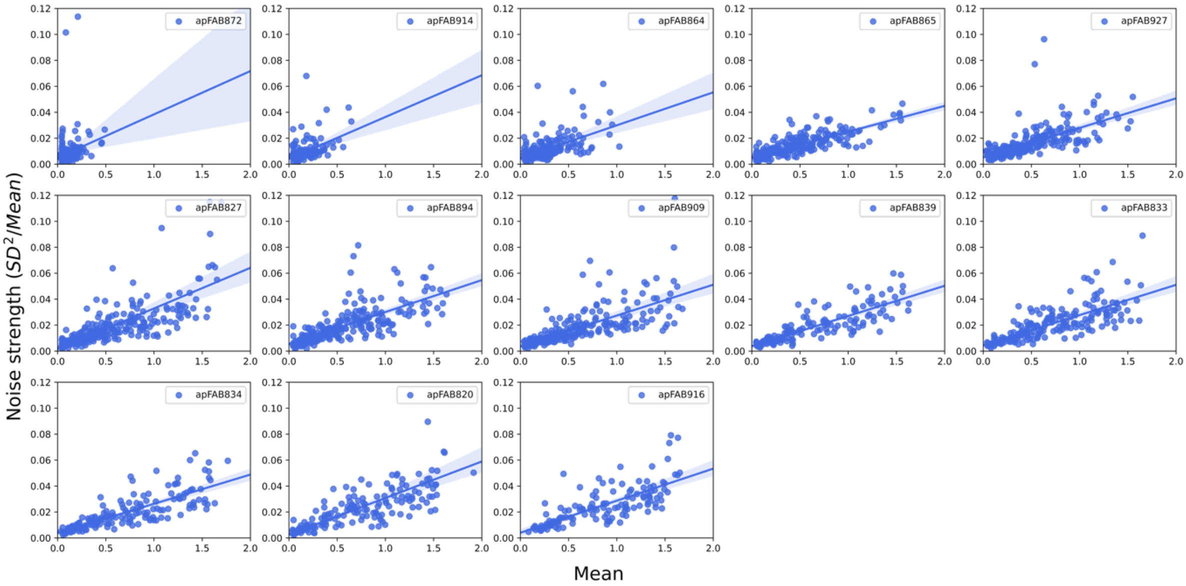


**Supplementary Figure 24** Noise strength as a function of the mean expression level when the transcription module varies. Each subplot corresponds to a group of strains with the same translation module. The lines are linear regressions (parameters are given in **Supplementary Table 6**), which are shaded to show the 95% confidence intervals.

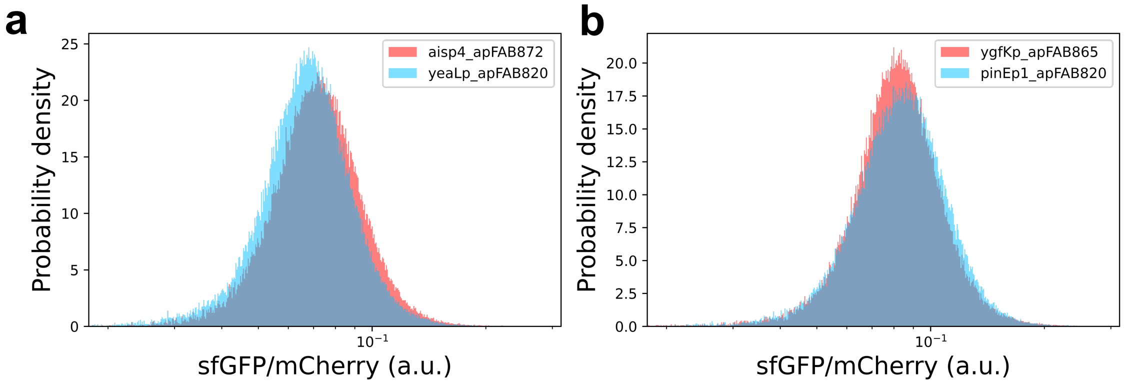


**Supplementary Figure 25** Fluorescence intensity distributions of strains with similar mean expression levels but different combinations of promoter and RBS (RBS strength: apFAB820 (1.57) > apFAB865 (0.67) > apFAB872 (0.15), see **Methods**). (**a**) aisp4_apFAB872: Mean = 0.07777, SD = 0.01469; yeaLp_apFAB820: Mean = 0.07318, SD = 0.01383. (**b**) ygfKp_apFAB865: Mean = 0.08869, SD = 0.01617; pinEp1_apFAB820: Mean = 0.09078, SD = 0.01737.

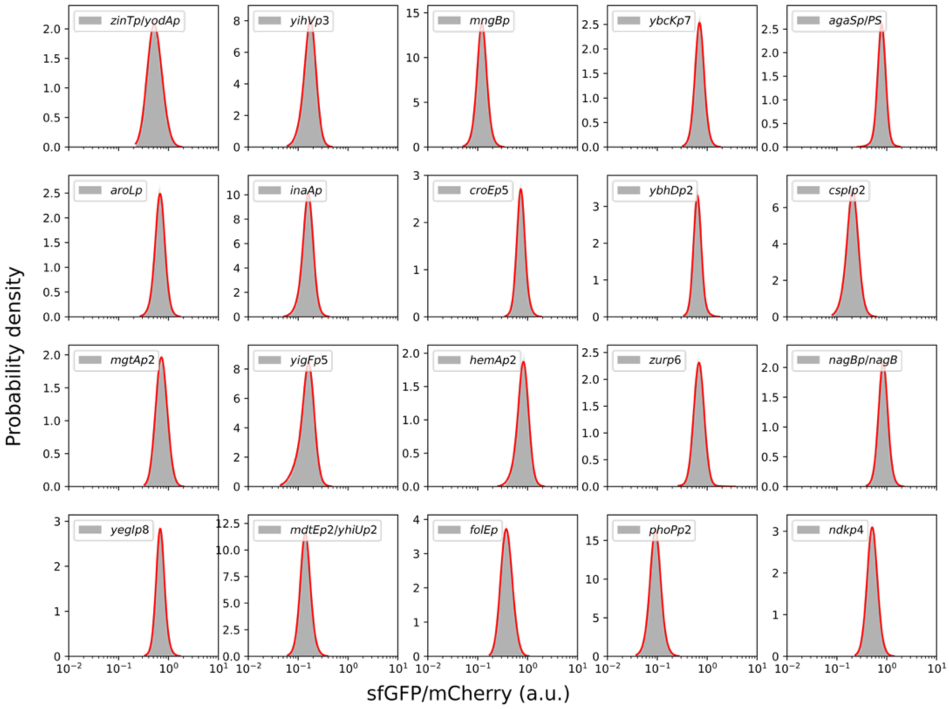


**Supplementary Figure 26** Individual cytometry assay of fluorescence intensity distributions of 20 promoters exhibiting high expression noise. Each distribution was fitted with a two-component LGMM.

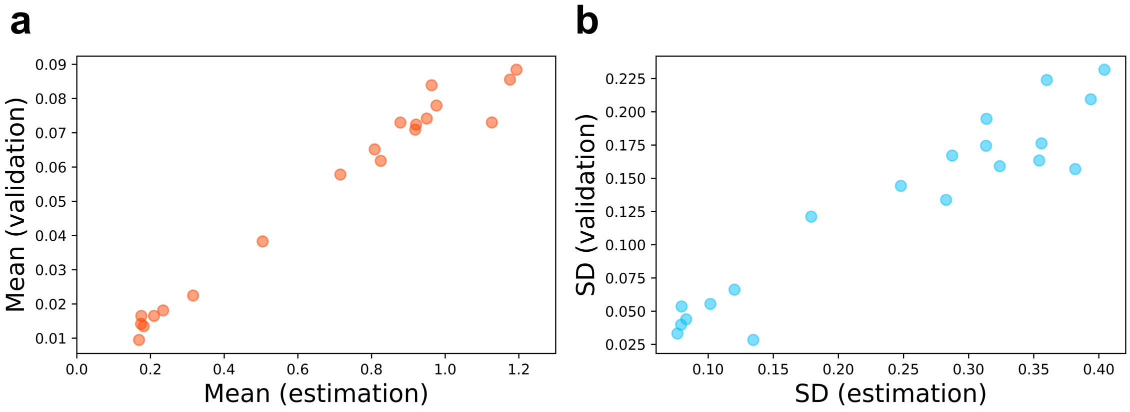


**Supplementary Figure 27** The expression properties of the promoters with high expression noise estimated from dSort-Seq were consistent with cytometry assays of individual fluorescence intensity distributions in terms of both mean (**a,** $n$ = 20, Pearson's $r$ = 0.966) and SD (**b,** $n$ = 20, Pearson's $r$ = 0.950).

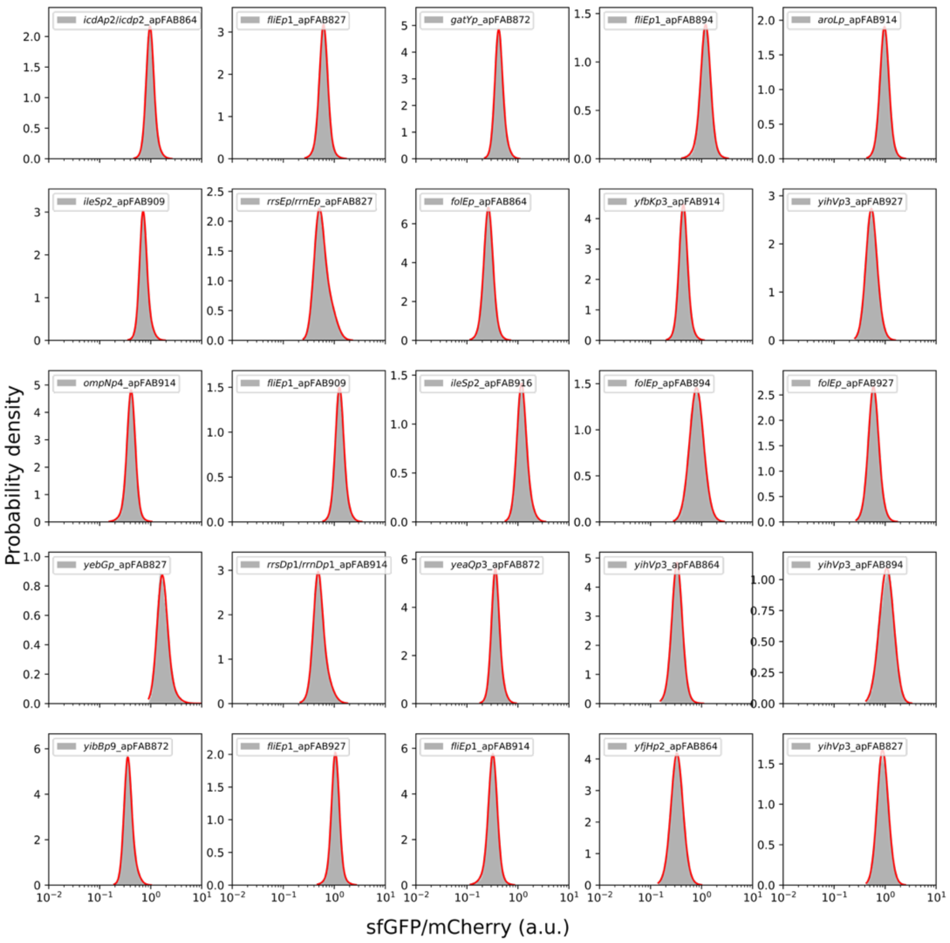


**Supplementary Figure 28** Individual cytometry assay of fluorescence intensity distributions of 25 combinations exhibiting high expression noise. Each distribution was fitted with a two-component LGMM.

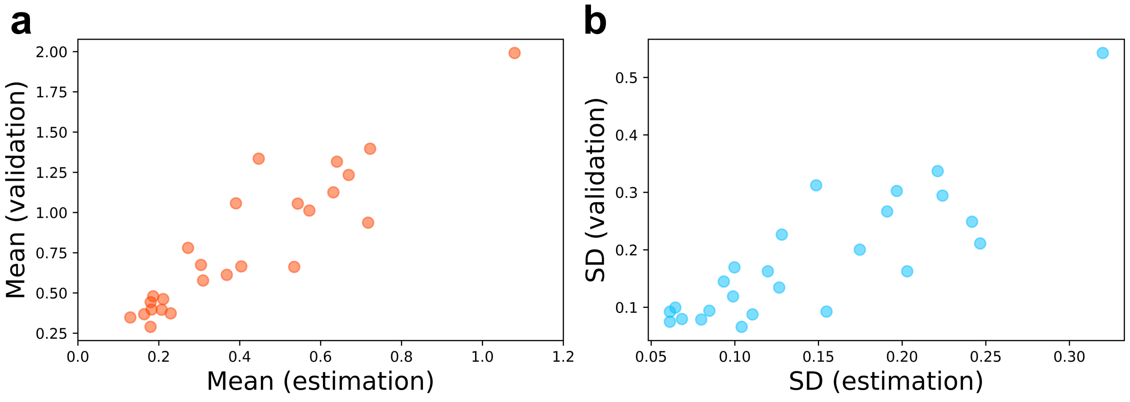


**Supplementary Figure 29** The expression properties of the combinations with high expression noise estimated from dSort-Seq were consistent with cytometry assays of individual fluorescence intensity distributions in terms of both mean (**a,** $n$ = 25, Pearson's $r$ = 0.915) and SD (**b,** $n$ = 25, Pearson's $r$ = 0.842).

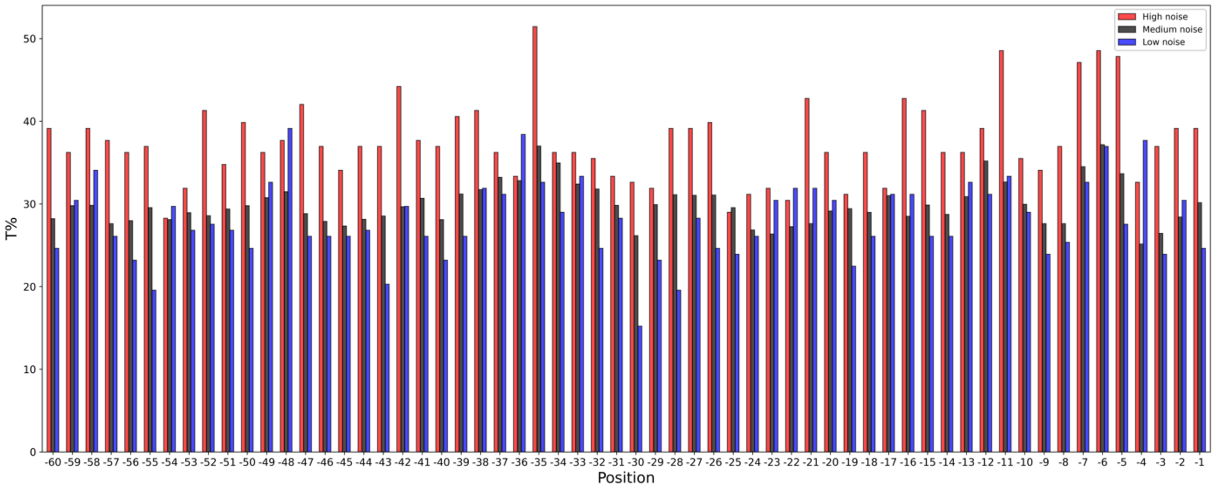


**Supplementary Figure 30** The thymidine proportion in the high-noise group was generally higher than that in other groups at all positions along the promoter sequence, indicating that it is not an increase in thymidine in a specific position that leads to high expression noise.

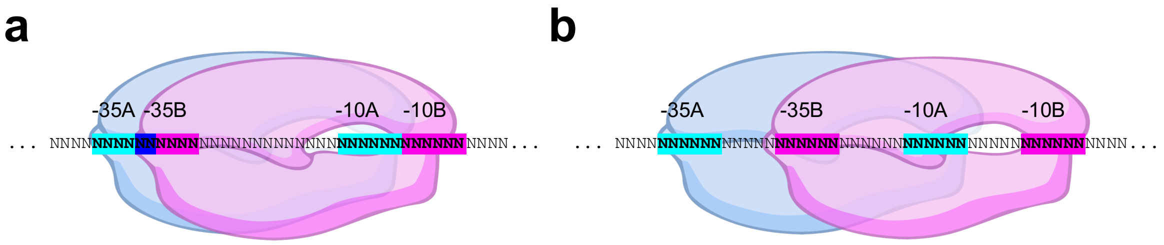


**Supplementary Figure 31** Different scenarios for searching the DPinteract database. (**a**) If the -35 or -10 regions of two RpoD-binding sites overlapped, only the promoter with a higher z score was accounted for. (**b**) Otherwise, both promoters were retained in subsequent analysis.

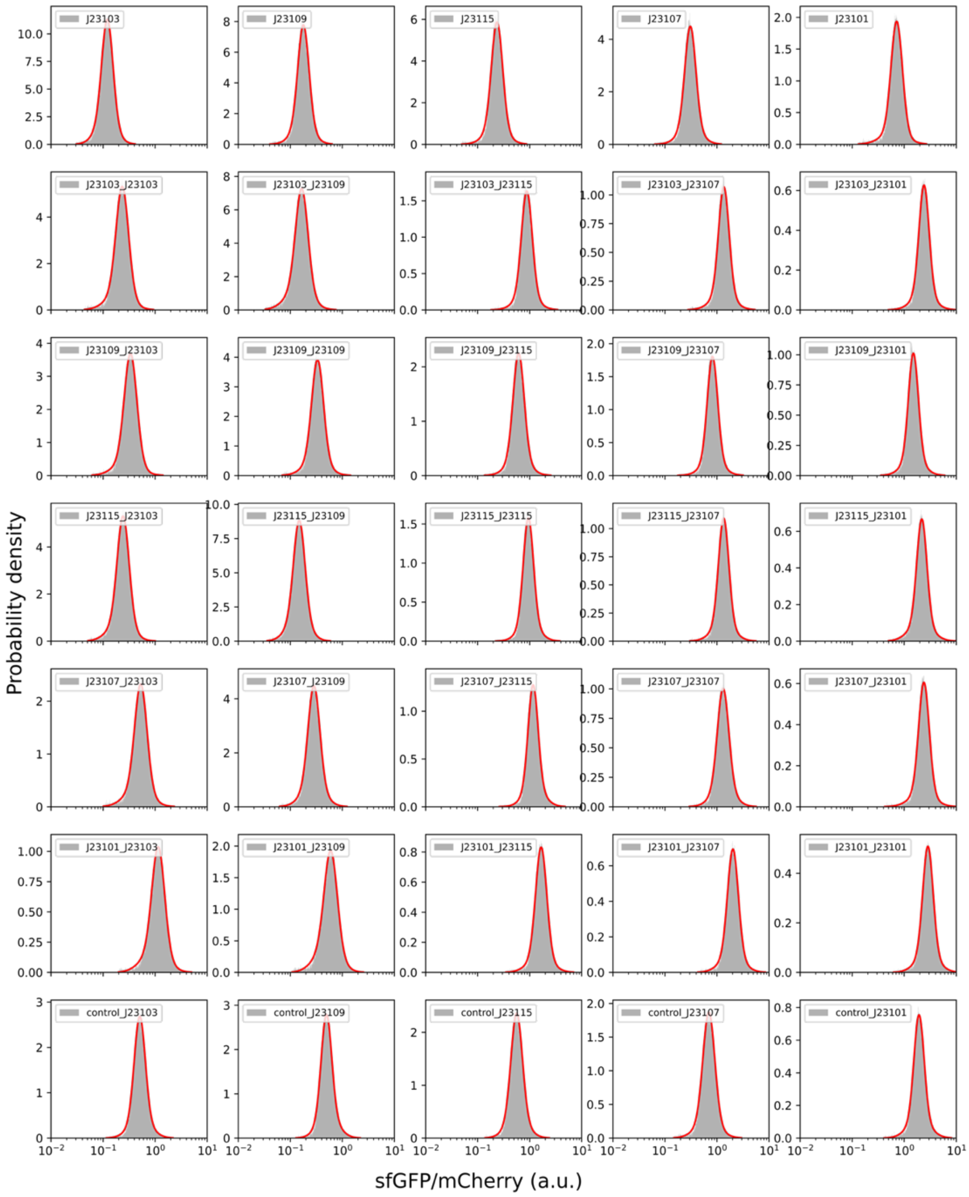


**Supplementary Figure 32** Individual cytometry assay of fluorescence intensity distributions of 5 constitutive Anderson promoters, 25 tandem promoters with overlapping RpoD-binding sites and 5 Anderson promoters with the same length as the tandem promoters. Each distribution was fitted with a two-component LGMM.

**Supplementary Table 1** Strains and plasmids used in this work

| Strains/plasmids | Characteristics | Sources |
|---|---|---|
| **Strains** | | |
| *E. coli* K12 MG1655 | Wild type | ATCC 700926 |
| BY4700 | *MATa ura3Δ0* | Gift from DaiLab |
| **Plasmids** | | |
| pSCSen_tnaC_change | Precursors for construction of *tnaC* variant reporter plasmids, pSC101, $Amp^R$ | [1] |
| pSCSen_tnaC_WT | Two-reporter plasmid with wild type *tnaC*, $Amp^R$ | This study |
| pSCSen_tnaC_L4R_CGG | Two-reporter plasmid with mutation L4R_CGG in *tnaC*, $Amp^R$ | This study |
| pSCSen_tnaC_L4Q_CAG | Two-reporter plasmid with mutation L4Q_CAG in *tnaC*, $Amp^R$ | This study |
| pSCSen_tnaC_D21T_ACC | Two-reporter plasmid with mutation D21T_ACC in *tnaC*, $Amp^R$ | This study |
| pSCSen_tnaC_D21S_TCC | Two-reporter plasmid with mutation D21S_TCC in *tnaC*, $Amp^R$ | This study |
| pCYC1-OP_TATA-UAS_FAC | Two-reporter plasmid with the combination of *pCYC1*, OP_TATA and $UAS_F$-$UAS_A$-$UAS_C$, $Amp^R$ | [2] |
| pCYC1-OP_TATA-UAS_DDC | Two-reporter plasmid with the combination of *pCYC1*, OP_TATA and $UAS_D$-$UAS_D$-$UAS_C$, $Amp^R$ | This study |
| pCYC1-OP_TATA-UAS_EBC | Two-reporter plasmid with the combination of *pCYC1*, OP_TATA and $UAS_E$-$UAS_B$-$UAS_C$, $Amp^R$ | This study |
| pHSP12-TATA_OP-UAS_FAC | Two-reporter plasmid with the combination of *pHSP12*, TATA_OP and $UAS_F$-$UAS_A$-$UAS_C$, $Amp^R$ | [2] |
| pHSP12-TATA_OP-UAS_BDC | Two-reporter plasmid with the combination of *pHSP12*, TATA_OP and $UAS_B$-$UAS_D$-$UAS_C$, $Amp^R$ | This study |
| pCYC1-OP_TATA-UAS_BEC | Two-reporter plasmid with the combination of *pCYC1*, OP_TATA and $UAS_B$-$UAS_E$-$UAS_C$, $Amp^R$ | This study |
| pEXG1-TATA_OP-UAS_FAC | Two-reporter plasmid with the combination of *pEXG1*, TATA_OP and $UAS_F$-$UAS_A$-$UAS_C$, $Amp^R$ | [2] |
| pEXG1-N30_OP-UAS_FDA | Two-reporter plasmid with the combination of *pEXG1*, N30_OP and $UAS_F$-$UAS_D$-$UAS_A$, $Amp^R$ | This study |
| pCYC1-TATA_OP-UAS_EAC | Two-reporter plasmid with the combination of *pCYC1*, TATA_OP and $UAS_E$-$UAS_A$-$UAS_C$, $Amp^R$ | This study |
| pCYC1-OP_TATA-UAS_FDC | Two-reporter plasmid with the combination of *pCYC1*, OP_TATA and $UAS_F$-$UAS_D$-$UAS_C$, $Amp^R$ | This study |

| | | |
|---|---|---|
| pCYC1-OP_TATA-UAS_BDC | Two-reporter plasmid with the combination of *pCYC1*, OP_TATA and $UAS_B$-$UAS_D$-$UAS_C$, $Amp^R$ | This study |
| POT1-pTEF2-mCherry-tADH1 | Positive control plasmid for mCherry, $Amp^R$ | [2] |
| POT1-pCYC1-YPet-tPGK1 | Positive control plasmid for YPet, $Amp^R$ | [2] |
| pACYCDuet-1 | p15A replication of origin, $Cm^R$ | Lab stock |
| pMPTPV | Derived from pACYCDuet-1 by replacing the chloramphenicol resistance gene with the kanamycin resistance gene, replacing the *lacI* expression cassette with *mcherry*, $Kan^R$ | Lab stock |
| pMPTPV_dual_fluorescence | Derived from pMPTPV, by inserting the *sfgfp* cassette into the opposite strand of *mcherry*, which is controlled by a variable region containing two *Bsm*BI restriction sites, $Kan^R$ | This study |
| pMPTPV_dual_dacBp5_BBa_J61106 | Two-reporter plasmid, in which the expression of *sfgfp* is under the control of promoter *dacBp5* and RBS BBa_J61106, $Kan^R$ | This study |
| pMPTPV_dual_ybhBp5_BBa_J61106 | Two-reporter plasmid, in which the expression of *sfgfp* is under the control of promoter *ybhBp5* and RBS BBa_J61106, $Kan^R$ | This study |
| pMPTPV_dual_ppkp6_BBa_J61106 | Two-reporter plasmid, in which the expression of *sfgfp* is under the control of promoter *ppkp6* and RBS BBa_J61106, $Kan^R$ | This study |
| pMPTPV_dual_yfaLp3_BBa_J61106 | Two-reporter plasmid, in which the expression of *sfgfp* is under the control of promoter *yfaLp3* and RBS BBa_J61106, $Kan^R$ | This study |
| pMPTPV_dual_ahpCp2_BBa_J61106 | Two-reporter plasmid, in which the expression of *sfgfp* is under the control of promoter *ahpCp2* and RBS BBa_J61106, $Kan^R$ | This study |
| pMPTPV_dual_xylEp_BBa_J61106 | Two-reporter plasmid, in which the expression of *sfgfp* is under the control of promoter *xylEp* and RBS BBa_J61106, $Kan^R$ | This study |
| pMPTPV_dual_mltAp6_BBa_J61106 | Two-reporter plasmid, in which the expression of *sfgfp* is under the control of promoter *mltAp6* and RBS BBa_J61106, $Kan^R$ | This study |
| pMPTPV_dual_lysAp_BBa_J61106 | Two-reporter plasmid, in which the expression of *sfgfp* is under the control of promoter *lysAp* and RBS BBa_J61106, $Kan^R$ | This study |
| pMPTPV_dual_rpoSp4_BBa_J61106 | Two-reporter plasmid, in which the expression of *sfgfp* is under the control of promoter *rpoSp4*and RBS BBa_J61106, $Kan^R$ | This study |
| pMPTPV_dual_yacHp3_BBa_J61106 | Two-reporter plasmid, in which the expression of *sfgfp* is under the control | This study |

| | | |
|---|---|---|
| | of promoter *yacHp3* and RBS BBa_J61106, Kan$^R$ | |
| pMPTPV_dual_aisp4_apFAB872 | Two-reporter plasmid, in which the expression of *sfgfp* is under the control of promoter *aisp4* and RBS apFAB872, Kan$^R$ | This study |
| pMPTPV_dual_yeaLp_apFAB820 | Two-reporter plasmid, in which the expression of *sfgfp* is under the control of promoter *yeaLp* and RBS apFAB820, Kan$^R$ | This study |
| pMPTPV_dual_ygfKp_apFAB865 | Two-reporter plasmid, in which the expression of *sfgfp* is under the control of promoter *ygfKp* and RBS apFAB865, Kan$^R$ | This study |
| pMPTPV_dual_pinEp1_apFAB820 | Two-reporter plasmid, in which the expression of *sfgfp* is under the control of promoter *pinEp1*and RBS apFAB820, Kan$^R$ | This study |
| pMPTPV_dual_yihVp3_BBa_J61106 | Two-reporter plasmid, in which the expression of *sfgfp* is under the con trol of promoter *yihVp3* and RBS B Ba_J61106, Kan$^R$ | This study |
| pMPTPV_dual_zinTp_BBa_J61106 | Two-reporter plasmid, in which the expression of *sfgfp* is under the control of promoter *zinTp* and RBS BBa_J61106, Kan$^R$ | This study |
| pMPTPV_dual_inaAp_BBa_J61106 | Two-reporter plasmid, in which the expression of *sfgfp* is under the control of promoter *inaAp* and RBS BBa_J61106, Kan$^R$ | This study |
| pMPTPV_dual_mngBp_BBa_J61106 | Two-reporter plasmid, in which the expression of *sfgfp* is under the control of promoter *mngBp* and RBS BBa_J61106, Kan$^R$ | This study |
| pMPTPV_dual_aroLp_BBa_J61106 | Two-reporter plasmid, in which the expression of *sfgfp* is under the control of promoter *aroLp* and RBS BBa_J61106, Kan$^R$ | This study |
| pMPTPV_dual_folEp_BBa_J61106 | Two-reporter plasmid, in which the expression of *sfgfp* is under the control of promoter *folEp* and RBS BBa_J61106, Kan$^R$ | This study |
| pMPTPV_dual_agaSp_BBa_J61106 | Two-reporter plasmid, in which the expression of *sfgfp* is under the control of promoter *agaSp* and RBS BBa_J61106, Kan$^R$ | This study |
| pMPTPV_dual_ybhDp2_BBa_J61106 | Two-reporter plasmid, in which the expression of *sfgfp* is under the control of promoter *ybhDp2* and RBS BBa_J61106, Kan$^R$ | This study |
| pMPTPV_dual_mgtAp2_BBa_J61106 | Two-reporter plasmid, in which the expression of *sfgfp* is under the control of promoter *mgtAp2* and RBS BBa_J61106, Kan$^R$ | This study |
| pMPTPV_dual_cspIp2_BBa_J61106 | Two-reporter plasmid, in which the expression of *sfgfp* is under the control of promoter *cspIp2* and RBS BBa_J61106, Kan$^R$ | This study |

| | | |
|---|---|---|
| pMPTPV_dual_hemAp2_BBa_J61106 | Two-reporter plasmid, in which the expression of *sfgfp* is under the control of promoter *hemAp2* and RBS BBa_J61106, $Kan^R$ | This study |
| pMPTPV_dual_yigFp5_BBa_J61106 | Two-reporter plasmid, in which the expression of *sfgfp* is under the control of promoter *yigFp5* and RBS BBa_J61106, $Kan^R$ | This study |
| pMPTPV_dual_mdtEp2_BBa_J61106 | Two-reporter plasmid, in which the expression of *sfgfp* is under the control of promoter *mdtEp2* and RBS BBa_J61106, $Kan^R$ | This study |
| pMPTPV_dual_nagBp_BBa_J61106 | Two-reporter plasmid, in which the expression of *sfgfp* is under the control of promoter *nagBp* and RBS BBa_J61106, $Kan^R$ | This study |
| pMPTPV_dual_zurp6_BBa_J61106 | Two-reporter plasmid, in which the expression of *sfgfp* is under the control of promoter zur*p6* and RBS BBa_J61106, $Kan^R$ | This study |
| pMPTPV_dual_croEp5_BBa_J61106 | Two-reporter plasmid, in which the expression of *sfgfp* is under the control of promoter *croEp5* and RBS BBa_J61106, $Kan^R$ | This study |
| pMPTPV_dual_yegIp8_BBa_J61106 | Two-reporter plasmid, in which the expression of *sfgfp* is under the control of promoter *yegIp8* and RBS BBa_J61106, $Kan^R$ | This study |
| pMPTPV_dual_ybcKp7_BBa_J61106 | Two-reporter plasmid, in which the expression of *sfgfp* is under the control of promoter *ybcKp7* and RBS BBa_J61106, $Kan^R$ | This study |
| pMPTPV_dual_phoPp2_BBa_J61106 | Two-reporter plasmid, in which the expression of *sfgfp* is under the control of promoter *phoPp2* and RBS BBa_J61106, $Kan^R$ | This study |
| pMPTPV_dual_ndkp4_BBa_J61106 | Two-reporter plasmid, in which the expression of *sfgfp* is under the control of promoter *ndkp4* and RBS BBa_J61106, $Kan^R$ | This study |
| pMPTPV_dual_icdAp2_apFAB864 | Two-reporter plasmid, in which the expression of *sfgfp* is under the control of promoter *icdAp2* and RBS apFAB864, $Kan^R$ | This study |
| pMPTPV_dual_fliEp1_apFAB894 | Two-reporter plasmid, in which the expression of *sfgfp* is under the control of promoter *fliEp1* and RBS apFAB894, $Kan^R$ | This study |
| pMPTPV_dual_gatYp_apFAB872 | Two-reporter plasmid, in which the expression of *sfgfp* is under the control of promoter *gatYp* and RBS apFAB872, $Kan^R$ | This study |
| pMPTPV_dual_yibBp9_apFAB872 | Two-reporter plasmid, in which the expression of *sfgfp* is under the control of promoter *yibBp9* and RBS apFAB872, $Kan^R$ | This study |
| pMPTPV_dual_fliEp1_apFAB827 | Two-reporter plasmid, in which the expression of *sfgfp* is under the control of promoter *fliEp1* and RBS apFAB827, $Kan^R$ | This study |

| | | |
|---|---|---|
| pMPTPV_dual_yeaQp3_apFAB872 | Two-reporter plasmid, in which the expression of *sfgfp* is under the control of promoter *yeaQp3* and RBS apFAB872, $Kan^R$ | This study |
| pMPTPV_dual_ileSp2_apFAB909 | Two-reporter plasmid, in which the expression of *sfgfp* is under the control of promoter *ileSp2* and RBS apFAB909, $Kan^R$ | This study |
| pMPTPV_dual_rrsEp_apFAB827 | Two-reporter plasmid, in which the expression of *sfgfp* is under the control of promoter *rrsEp* and RBS apFAB827, $Kan^R$ | This study |
| pMPTPV_dual_yihVp3_apFAB927 | Two-reporter plasmid, in which the expression of *sfgfp* is under the control of promoter *yihVp3* and RBS apFAB927, $Kan^R$ | This study |
| pMPTPV_dual_ompNp4_apFAB914 | Two-reporter plasmid, in which the expression of *sfgfp* is under the control of promoter *ompNp4* and RBS apFAB914, $Kan^R$ | This study |
| pMPTPV_dual_ileSp2_apFAB916 | Two-reporter plasmid, in which the expression of *sfgfp* is under the control of promoter *ileSp2* and RBS apFAB916, $Kan^R$ | This study |
| pMPTPV_dual_folEp_apFAB864 | Two-reporter plasmid, in which the expression of *sfgfp* is under the control of promoter *folEp* and RBS apFAB864, $Kan^R$ | This study |
| pMPTPV_dual_fliEp1_apFAB909 | Two-reporter plasmid, in which the expression of *sfgfp* is under the control of promoter *fliEp1* and RBS apFAB909, $Kan^R$ | This study |
| pMPTPV_dual_folEp_apFAB894 | Two-reporter plasmid, in which the expression of *sfgfp* is under the control of promoter *folEp* and RBS apFAB894, $Kan^R$ | This study |
| pMPTPV_dual_yebGp_apFAB827 | Two-reporter plasmid, in which the expression of *sfgfp* is under the control of promoter *yebGp* and RBS apFAB827, $Kan^R$ | This study |
| pMPTPV_dual_folEp_apFAB927 | Two-reporter plasmid, in which the expression of *sfgfp* is under the control of promoter *folEp* and RBS apFAB927, $Kan^R$ | This study |
| pMPTPV_dual_yihVp3_apFAB864 | Two-reporter plasmid, in which the expression of *sfgfp* is under the control of promoter *yihVp3* and RBS apFAB864, $Kan^R$ | This study |
| pMPTPV_dual_rrsDp1_apFAB914 | Two-reporter plasmid, in which the expression of *sfgfp* is under the control of promoter *rrsDp1* and RBS apFAB914, $Kan^R$ | This study |
| pMPTPV_dual_fliEp1_apFAB927 | Two-reporter plasmid, in which the expression of *sfgfp* is under the control of promoter *fliEp1* and RBS apFAB927, $Kan^R$ | This study |
| pMPTPV_dual_aroLp_apFAB914 | Two-reporter plasmid, in which the expression of *sfgfp* is under the control of promoter *aroLp* and RBS apFAB914, $Kan^R$ | This study |

| | | |
|---|---|---|
| pMPTPV_dual_yihVp3_apFAB894 | Two-reporter plasmid, in which the expression of *sfgfp* is under the control of promoter *yihVp3* and RBS apFAB894, $Kan^R$ | This study |
| pMPTPV_dual_fliEp1_apFAB914 | Two-reporter plasmid, in which the expression of *sfgfp* is under the control of promoter *fliEp1* and RBS apFAB914, $Kan^R$ | This study |
| pMPTPV_dual_yfbKp3_apFAB914 | Two-reporter plasmid, in which the expression of *sfgfp* is under the control of promoter *yfbKp3* and RBS apFAB914, $Kan^R$ | This study |
| pMPTPV_dual_yfjHp2_apFAB864 | Two-reporter plasmid, in which the expression of *sfgfp* is under the control of promoter *yfjHp2* and RBS apFAB864, $Kan^R$ | This study |
| pMPTPV_dual_yihVp3_apFAB827 | Two-reporter plasmid, in which the expression of *sfgfp* is under the control of promoter *yihVp3* and RBS apFAB827, $Kan^R$ | This study |
| pMPTPV_dual_J23103_BBa_J61106 | Two-reporter plasmid, in which the expression of *sfgfp* is under the control of promoter J23103 and RBS BBa_J61106, $Kan^R$ | This study |
| pMPTPV_dual_J23109_BBa_J61106 | Two-reporter plasmid, in which the expression of *sfgfp* is under the control of promoter J23109 and RBS BBa_J61106, $Kan^R$ | This study |
| pMPTPV_dual_J23115_BBa_J61106 | Two-reporter plasmid, in which the expression of *sfgfp* is under the control of promoter J23115 and RBS BBa_J61106, $Kan^R$ | This study |
| pMPTPV_dual_J23107_BBa_J61106 | Two-reporter plasmid, in which the expression of *sfgfp* is under the control of promoter J23107 and RBS BBa_J61106, $Kan^R$ | This study |
| pMPTPV_dual_J23101_BBa_J61106 | Two-reporter plasmid, in which the expression of *sfgfp* is under the control of promoter J23101 and RBS BBa_J61106, $Kan^R$ | This study |
| pMPTPV_dual_J23103_J23103_BBa_J61106 | Two-reporter plasmid, in which the expression of *sfgfp* is under the control of promoter J23103_J23103 and RBS BBa_J61106, $Kan^R$ | This study |
| pMPTPV_dual_J23103_J23109_BBa_J61106 | Two-reporter plasmid, in which the expression of *sfgfp* is under the control of promoter J23103_J23109 and RBS BBa_J61106, $Kan^R$ | This study |
| pMPTPV_dual_J23103_J23115_BBa_J61106 | Two-reporter plasmid, in which the expression of *sfgfp* is under the control of promoter J23103_J23115 and RBS BBa_J61106, $Kan^R$ | This study |
| pMPTPV_dual_J23103_J23107_BBa_J61106 | Two-reporter plasmid, in which the expression of *sfgfp* is under the control of promoter J23103_J23107 and RBS BBa_J61106, $Kan^R$ | This study |
| pMPTPV_dual_J23103_J23101_BBa_J61106 | Two-reporter plasmid, in which the expression of *sfgfp* is under the control of promoter J23103_J23101 and RBS BBa_J61106, $Kan^R$ | This study |

| | | |
|---|---|---|
| pMPTPV_dual_J23109_J23103_BBa_J61106 | Two-reporter plasmid, in which the expression of *sfgfp* is under the control of promoter J23109_J23103 and RBS BBa_J61106, $Kan^R$ | This study |
| pMPTPV_dual_J23109_J23109_BBa_J61106 | Two-reporter plasmid, in which the expression of *sfgfp* is under the control of promoter J23109_J23109 and RBS BBa_J61106, $Kan^R$ | This study |
| pMPTPV_dual_J23109_J23115_BBa_J61106 | Two-reporter plasmid, in which the expression of *sfgfp* is under the control of promoter J23109_J23115 and RBS BBa_J61106, $Kan^R$ | This study |
| pMPTPV_dual_J23109_J23107_BBa_J61106 | Two-reporter plasmid, in which the expression of *sfgfp* is under the control of promoter J23109_J23107 and RBS BBa_J61106, $Kan^R$ | This study |
| pMPTPV_dual_J23109_J23101_BBa_J61106 | Two-reporter plasmid, in which the expression of *sfgfp* is under the control of promoter J23109_J23101 and RBS BBa_J61106, $Kan^R$ | This study |
| pMPTPV_dual_J23115_J23103_BBa_J61106 | Two-reporter plasmid, in which the expression of *sfgfp* is under the control of promoter J23115_J23103 and RBS BBa_J61106, $Kan^R$ | This study |
| pMPTPV_dual_J23115_J23109_BBa_J61106 | Two-reporter plasmid, in which the expression of *sfgfp* is under the control of promoter J23115_J23109 and RBS BBa_J61106, $Kan^R$ | This study |
| pMPTPV_dual_J23115_J23115_BBa_J61106 | Two-reporter plasmid, in which the expression of *sfgfp* is under the control of promoter J23115_J23115 and RBS BBa_J61106, $Kan^R$ | This study |
| pMPTPV_dual_J23115_J23107_BBa_J61106 | Two-reporter plasmid, in which the expression of *sfgfp* is under the control of promoter J23115_J23107 and RBS BBa_J61106, $Kan^R$ | This study |
| pMPTPV_dual_J23115_J23101_BBa_J61106 | Two-reporter plasmid, in which the expression of *sfgfp* is under the control of promoter J23115_J23101 and RBS BBa_J61106, $Kan^R$ | This study |
| pMPTPV_dual_J23107_J23103_BBa_J61106 | Two-reporter plasmid, in which the expression of *sfgfp* is under the control of promoter J23107_J23103 and RBS BBa_J61106, $Kan^R$ | This study |
| pMPTPV_dual_J23107_J23109_BBa_J61106 | Two-reporter plasmid, in which the expression of *sfgfp* is under the control of promoter J23107_J23109 and RBS BBa_J61106, $Kan^R$ | This study |
| pMPTPV_dual_J23107_J23115_BBa_J61106 | Two-reporter plasmid, in which the expression of *sfgfp* is under the control of promoter J23107_J23115 and RBS BBa_J61106, $Kan^R$ | This study |
| pMPTPV_dual_J23107_J23107_BBa_J61106 | Two-reporter plasmid, in which the expression of *sfgfp* is under the control of promoter J23107_J23107 and RBS BBa_J61106, $Kan^R$ | This study |
| pMPTPV_dual_J23107_J23101_BBa_J61106 | Two-reporter plasmid, in which the expression of *sfgfp* is under the control of promoter J23107_J23101 and RBS BBa_J61106, $Kan^R$ | This study |

| | | |
|---|---|---|
| pMPTPV_dual_J23101_J23103_BBa_J61106 | Two-reporter plasmid, in which the expression of *sfgfp* is under the control of promoter J23101_J23103 and RBS BBa_J61106, $Kan^R$ | This study |
| pMPTPV_dual_J23101_J23109_BBa_J61106 | Two-reporter plasmid, in which the expression of *sfgfp* is under the control of promoter J23101_J23109 and RBS BBa_J61106, $Kan^R$ | This study |
| pMPTPV_dual_J23101_J23115_BBa_J61106 | Two-reporter plasmid, in which the expression of *sfgfp* is under the control of promoter J23101_J23115 and RBS BBa_J61106, $Kan^R$ | This study |
| pMPTPV_dual_J23101_J23107_BBa_J61106 | Two-reporter plasmid, in which the expression of *sfgfp* is under the control of promoter J23101_J23107 and RBS BBa_J61106, $Kan^R$ | This study |
| pMPTPV_dual_J23101_J23101_BBa_J61106 | Two-reporter plasmid, in which the expression of *sfgfp* is under the control of promoter J23101_J23101 and RBS BBa_J61106, $Kan^R$ | This study |
| pMPTPV_dual_control_J23103_BBa_J61106 | Two-reporter plasmid, in which the expression of *sfgfp* is under the control of promoter control_J23103 and RBS BBa_J61106, $Kan^R$ | This study |
| pMPTPV_dual_control_J23109_BBa_J61106 | Two-reporter plasmid, in which the expression of *sfgfp* is under the control of promoter control_J23109 and RBS BBa_J61106, $Kan^R$ | This study |
| pMPTPV_dual_control_J23115_BBa_J61106 | Two-reporter plasmid, in which the expression of *sfgfp* is under the control of promoter control_J23115 and RBS BBa_J61106, $Kan^R$ | This study |
| pMPTPV_dual_control_J23107_BBa_J61106 | Two-reporter plasmid, in which the expression of *sfgfp* is under the control of promoter control_J23107 and RBS BBa_J61106, $Kan^R$ | This study |
| pMPTPV_dual_control_J23101_BBa_J61106 | Two-reporter plasmid, in which the expression of *sfgfp* is under the control of promoter control_J23101 and RBS BBa_J61106, $Kan^R$ | This study |

**Supplementary Table 2** Primers and other oligonucleotides used in this work

| Primers | Sequences | Usages |
|---|---|---|
| F-tnaC_WT | TTATATGAATATCTTACATATATGTGTGACCTCAAAATGGTTCAATATTGACAACAAAATTGTCGATCACCGCCCTTGA | Oligonucleotides used in Golden Gate Assembly to construct pSCSen_tnaC_WT |
| R-tnaC_WT | AATCTCAAGGGCGGTGATCGACAATTTTGTTGTCAATATTGAACCATTTTGAGGTCACACATATATGTAAGATATTCAT | |
| F-tnaC_L4R_CGG | TTATATGAATATCCGGCATATATGTGTGACCTCAAAATGGTTCAATATTGACAACAAAATTGTCGATCACCGCCCTTGA | Oligonucleotides used in Golden Gate Assembly to construct pSCSen_tnaC_ L4R_CGG |
| R-tnaC_L4R_CGG | AATCTCAAGGGCGGTGATCGACAATTTTGTTGTCAATATTGAACCATTTTGAGGTCACACATATATGCCGGATATTCAT | |
| F-tnaC_L4Q_CAG | TTATATGAATATCCAGCATATATGTGTGACCTCAAAATGGTTCAATATTGACAACAAAATTGTCGATCACCGCCCTTGA | Oligonucleotides used in Golden Gate Assembly to construct pSCSen_tnaC_ L4Q_CAG |
| R--tnaC_L4Q_CAG | AATCTCAAGGGCGGTGATCGACAATTTTGTTGTCAATATTGAACCATTTTGAGGTCACACATATATGCTGGATATTCAT | |
| F-tnaC_D21T_ACC | TTATATGAATATCTTACATATATGTGTGACCTCAAAATGGTTCAATATTGACAACAAAATTGTCACCCACCGCCCTTGA | Oligonucleotides used in Golden Gate Assembly to construct pSCSen_tnaC_ D21T_ACC |
| R-tnaC_D21T_ACC | AATCTCAAGGGCGGTGGGTGACAATTTTGTTGTCAATATTGAACCATTTTGAGGTCACACATATATGTAAGATATTCAT | |
| F-tnaC_D21S_TCC | TTATATGAATATCTTACATATATGTGTGACCTCAAAATGGTTCAATATTGACAACAAAATTGTCTCCCACCGCCCTTGA | Oligonucleotides used in Golden Gate Assembly to construct pSCSen_tnaC_ D21S_TCC |
| R- tnaC_D21S_TCC | AATCTCAAGGGCGGTGGGAGACAATTTTGTTGTCAATATTGAACCATTTTGAGGTCACACATATATGTAAGATATTCAT | |
| oligo_1 | AATCAAGCTTATAAAAGAATTATATACTACTATTAGTACCTAGTCTTAATTAGCACTGTTGGGCGTGAGTGGAGGCGCCGG | Oligonucleotides used in Golden Gate Assembly to construct pCYC1-OP_TATA-UAS_DDC, pCYC1-OP_TATA-UAS_EBC, pHSP12-TATA_OP-UAS_BDC, pCYC1-TATA_OP-UAS_EAC, pCYC1-OP_TATA-UAS_BDC, pCYC1-OP_TATA-UAS_FDC, pEXG1-N30_OP-UAS_FDA and pCYC1-OP_TATA-UAS_BEC, for details, see **Supplementary Table 3** |
| oligo_2 | GTACTAATAGTAGTATATAATTAGCTTGATTAGAATATTACAAGTTAATTAATC | |
| oligo_3 | CGCCCAACAGTGCTCTTTTATAAGCTTGATTAGAATATTACAAGTTAATTAACA | |
| oligo_4 | GAGGCCGCTTTCTCAAAGCATAGGCGCGCCACTGAAATTTGGGGGCGGTGTAGCATGTGATTAATTAACTTGTAATATTCT | |
| oligo_5 | GAGGCCGCTTTCTCAAAGGCCAGGCGCGCCGCTCAACGGCACTGAAATTTTAGCATGTGATTAATTAACTTGTAATATTCT | |

| | | |
|---|---|---|
| oligo_6 | GAGGCCGCTTTCTCAAAGGCCAGGCGC<br>GCCGCTCAACGGCACAGAGGGGCTAGC<br>ATGTGATTAATTAACTTGTAATATTCT | Oligonucleotides used in Golden Gate Assembly to construct pCYC1-OP_TATA-UAS_DDC, pCYC1-OP_TATA-UAS_EBC, pHSP12-TATA_OP-UAS_BDC, pCYC1-TATA_OP-UAS_EAC, pCYC1-OP_TATA-UAS_BDC, pCYC1-OP_TATA-UAS_FDC, pEXG1-N30_OP-UAS_FDA and pCYC1-OP_TATA-UAS_BEC for details, see **Supplementary Table 3** |
| oligo_7 | AGTAGTATATAATTCTTTTATAAGCTTG<br>ATTAGAATATTACAAGTTAATTAATC | |
| oligo_8 | ACATGCTAAAATTTCAGTGCCGTTGAG<br>CGGCGCGCCTGGCCTTTGAGAAAGCGG | |
| oligo_9 | GAGGTTAGTTTCTCAAAGCATAGGCGC<br>GCCGCTCAACGGCACAGAGGGGCTAGC<br>ATGTGATTAATTAACTTGTAATATTCT | |
| oligo_10 | ACATGCTAGCCGTTGAGCAAATTTCAG<br>TGGCGCGCCTGGCCTTTGAGAAAGCGG | |
| oligo_11 | ACATGCTACACCGCCCCCAAATTTCAG<br>TGGCGCGCCTATGCTTTGAGAAAGCGG | |
| oligo_12 | GAGGGATCTTTCTCAAAGTGGTGGCGC<br>GCCCCTCCTTGAAACAGAGGGGCGGGG<br>GCGGTGTTAATTAACTTGTAATATTCT | |
| oligo_13 | AATCAAGCTAATTATATACTACTATTA<br>GTACCTAGTCTTAATTTATAAAAGAGC<br>ACTGTTGGGCGTGAGTGGAGGCGCCGG | |
| oligo_14 | GAGGCCGCTTTCTCAAAGGCCAGGCGC<br>GCCACTGAAATTTGCTCAACGGCTAGC<br>ATGTGATTAATTAACTTGTAATATTCT | |
| oligo_15 | TTTTAATTAAGACTAGGTACTAATAGT<br>AGTATATAATTCCGGCGCCTCCACTCA | |
| oligo_16 | AATCAAGCTTATAAAAGAGCACTGTTG<br>GGCGTGAGTGGAGGCGCCGGAATTATA<br>TACTACTATTAGTACCTAGTCTTAATT | |
| oligo_17 | ACATGCTAGCCCCTCTGTGCCGTTGAG<br>CGGCGCGCCTGGCCTTTGAGAAAGCGG | |
| oligo_18 | GAGGCCGCTTTCTCAAAGGCCAGGCGC<br>GCCCCTCCTTGAAACAGAGGGGCTAGC<br>ATGTGATTAATTAACTTGTAATATTCT | |
| oligo_19 | ACATGCTAGCCCCTCTGTGCCCCTCTGT<br>GGCGCGCCTGGCCTTTGAGAAAGCGG | |
| oligo_20 | TTTTCCGGCGCCTCCACTCACGCCCAAC<br>AGTGCTCTTTTATAAATTAAGACTAG | |
| oligo_21 | CCGCCCCCGCCCCTCTGTTTCAAGGAG<br>GGGCGCGCCACCACTTTGAGAAAGATC | |

| | | |
|---|---|---|
| oligo_22 | TTTTCCGGCGCCTCCACTCACGCCCAACAGTGCTAATTAAGACTAGGTACTAAT | Oligonucleotides used in Golden Gate Assembly to construct pCYC1-OP_TATA-UAS_DDC, pCYC1-OP_TATA-UAS_EBC, pHSP12-TATA_OP-UAS_BDC, pCYC1-TATA_OP-UAS_EAC, pCYC1-OP_TATA-UAS_BDC, pCYC1-OP_TATA-UAS_FDC, pEXG1-N30_OP-UAS_FDA and pCYC1-OP_TATA-UAS_BEC, for details, see **Supplementary Table 3** |
| oligo_23 | ACATGCTAGCCCCTCTGTTTCAAGGAGGGCGCGCCTGGCCTTTGAGAAAGCGG | |
| oligo_24 | ACATGCTAGCCCCTCTGTGCCGTTGAGCGGCGCGCCTATGCTTTGAGAAACTAA | |
| oligo_25 | GAGGCCGCTTTCTCAAAGGCCAGGCGCGCCACAGAGGGGCACAGAGGGGCTAGCATGTGATTAATTAACTTGTAATATTCT | |
| F-Malonyl-CoA-Sen | cgGGTCTCACCTCAGGCggaatat | Primers used to amplify the backbones of pCYC1-OP_TATA-UAS_FAC, pHSP12-OP_TATA-UAS_FAC, pEXG1-OP_TATA-UAS_FAC. These backbones were used in Golden Gate Assembly to construct other Malonyl-CoA biosensors |
| R-Malonyl-CoA-Sen | tcGGTCTCGAAAAAAGCATCGAAAAAATCTA | |
| F-dacBp5_ BBa_J61106 | GGATcgggaccagaagcaaaaaaataccgacccgggtacaagtcccaggtcagctacaattcacTCTAGAGAAAGATAGGAGACACTAGT | Oligonucleotides used in Golden Gate Assembly to construct pMPTPV_dual_dacBp5_BBa_J61106 |
| R-dacBp5_ BBa_J61106 | GCATACTAGTGTCTCCTATCTTTCTCTAGAgtgaattgtagctgacctgggacttgtacccgggtcggtatttttttgcttctggtcccg | |
| F-xylEp_ BBa_J61106 | GGATgtaaacgcattgtaaaaaatgataattgccttaactgcctgacaattccaacatcaatgcTCTAGAGAAAGATAGGAGACACTAGT | Oligonucleotides used in Golden Gate Assembly to construct pMPTPV_dual_xylEp_BBa_J61106 |
| R-xylEp_ BBa_J61106 | GCATACTAGTGTCTCCTATCTTTCTCTAGAgcattgatgttggaattgtcaggcagttaaggcaattatcattttttacaatgcgtttac | |
| F-ybhBp4_ BBa_J61106 | GGATatgaaacatttttttgccaataacgagaagtccgcgtgagggtttctggctacactttctTCTAGAGAAAGATAGGAGACACTAGT | Oligonucleotides used in Golden Gate Assembly to construct pMPTPV_dual_ybhBp4_BBa_J61106 |
| R-ybhBp4_ BBa_J61106 | GCATACTAGTGTCTCCTATCTTTCTCTAGAagaaagtgtagccagaaaccctcacgcggacttctcgttattggcaaaaaaatgtttcat | |
| F-mltAp6_ BBa_J61106 | GGATcccgatcacggatttggcatgattttgaacaaaaacaaatcactaaagcctatttttttgtTCTAGAGAAAGATAGGAGACACTAGT | Oligonucleotides used in Golden Gate Assembly to construct pMPTPV_dual_mltAp6_BBa_J61106 |
| R-mltAp6_ BBa_J61106 | GCATACTAGTGTCTCCTATCTTTCTCTAGAacaaaaaaataggctttagtgatttgtttttgttcaaaatcatgccaaatccgtgatcggg | |
| F-ppkp6_ BBa_J61106 | GGATtcgcaagctccagcagtttttttcccccttttctggcatagttggacatctgccaatattTCTAGAGAAAGATAGGAGACACTAGT | Oligonucleotides used in Golden Gate Assembly to construct pMPTPV_dual_ppkp6_BBa_J61106 |
| R-ppkp6_ BBa_J61106 | GCATACTAGTGTCTCCTATCTTTCTCTAGAaatattggcagatgtccaactatgccagaaaagggggaaaaaaactgctggagcttgcga | |

| | | |
|---|---|---|
| F-lysAp_ BBa_J61106 | GGATctcgcaatccggtaatccatatcatttttgcatagactcgacataaatcgatattttttaTCTAGAGAAAGATAGGAGACACTAGT | Oligonucleotides used in Golden Gate Assembly to construct pMPTPV_dual_lysAp_BBa_J61106 |
| R-lysAp_ BBa_J61106 | GCATACTAGTGTCTCCTATCTTTCTCTAGAtaaaaaatatcgatttatgtcgagtctatgcaaaaatgatatggattaccggattgcgag | |
| F-yfaLp3_ BBa_J61106 | GGATaggttattaacttgaattatgacgcactgatattattcatcaaataataacaaaatagccTCTAGAGAAAGATAGGAGACACTAGT | Oligonucleotides used in Golden Gate Assembly to construct pMPTPV_dual_yfaLp3_BBa_J61106 |
| R-yfaLp3_ BBa_J61106 | GCATACTAGTGTCTCCTATCTTTCTCTAGAggctattttgttattatttgatgaataatatcagtgcgtcataattcaagttaataacct | |
| F-rpoSp4_ BBa_J61106 | GGATagtgcctacgcccataacgacacaatgctggtccgggaacaacaagaagttaaggcggggTCTAGAGAAAGATAGGAGACACTAGT | Oligonucleotides used in Golden Gate Assembly to construct pMPTPV_dual_rpoSp4_BBa_J61106 |
| R-rpoSp4_ BBa_J61106 | GCATACTAGTGTCTCCTATCTTTCTCTAGAccccgccttaacttcttgttgttcccggaccagcattgtgtcgttatgggcgtaggcact | |
| F-ahpCp2_ BBa_J61106 | GGATtaggtaagagcttagatcaggtgattgccctttgtttatgagggtgttgtaatccatgtcTCTAGAGAAAGATAGGAGACACTAGT | Oligonucleotides used in Golden Gate Assembly to construct pMPTPV_dual_ahpCp2_BBa_J61106 |
| R-ahpCp2_ BBa_J61106 | GCATACTAGTGTCTCCTATCTTTCTCTAGAgacatggattacaacaccctcataaacaaagggcaatcacctgatctaagctcttaccta | |
| F-yacHp3_ BBa_J61106 | GGATaatcccaagattagtcaaaatttaaactaccgcctctttatactcggattcacagcacctTCTAGAGAAAGATAGGAGACACTAGT | Oligonucleotides used in Golden Gate Assembly to construct pMPTPV_dual_yacHp3_BBa_J61106 |
| R-yacHp3_ BBa_J61106 | GCATACTAGTGTCTCCTATCTTTCTCTAGAaggtgctgtgaatccgagtataaagaggcggtagtttaaatttgactaatcttgggatt | |
| F-aisp4_apFAB872 | GGATtaagaaactaatattagacgtaaatattgaaatttttatattttttcttatttaggctttATCTTAATCTAGCCCGGGATAATTT | Oligonucleotides used in Golden Gate Assembly to construct pMPTPV_dual_aisp4_apFAB872 |
| R-aisp4_apFAB872 | GCATAAATTATCCCGGGCTAGATTAAGATaaagcctaaataagaaaaaatataaaaatttcaatatttacgtctaatattagtttctta | |
| F-yeaLp_apFAB820 | GGATtttatgcaaaaaacgtaaagattttctacttccttcctgcagcaagcgtaaagtaagcagATCTTAATCTAGCTGCGGAGGGTTT | Oligonucleotides used in Golden Gate Assembly to construct pMPTPV_dual_yeaLp_apFAB820 |
| R-yeaLp_apFAB820 | GCATAAACCCTCCGCAGCTAGATTAAGATctgcttactttacgcttgctgcaggaaggaagtagaaaatcttacgttttttgcataaa | |
| F-ygfKp_apFAB865 | GGATcctcacatttttttatatttcccgccaaacctggcaagagtggtgcgattgttgctctatATCTTAATCTAGCGCGGGAGTATTT | Oligonucleotides used in Golden Gate Assembly to construct pMPTPV_dual_ygfKp_apFAB865 |
| R-ygfKp_apFAB865 | GCATAAATACTCCCGCGCTAGATTAAGATatagagcaacaatcgcaccactcttgccaggtttggcgggaaatataaaaaaatgtgagg | |

| | | |
|---|---|---|
| F-pinEp1_apFAB820 | GGATcaatttatttgtcggataagcctttgccagttccgggtattcttcagcagaaaaagcggcATCTTAATCTAGCTGCGGAGGGTTT | Oligonucleotides used in Golden Gate Assembly to construct pMPTPV_dual_pinEp1_apFAB820 |
| R-pinEp1_apFAB820 | GCATAAACCCTCCGCAGCTAGATTAAGATgccgctttttctgctgaagaatacccggaactggcaaaggcttatccgacaaataaattg | |
| Lib_F | TGCGACTCCTGCATTAGGAAATG | Forward primer used to clone the promoter and RBS region |
| Lib_R | CAGTGAACAGCTCTTCGCCTTT | Reverse primer used to clone the promoter and RBS region |
| sorting_P2 | CCGAACCATGCGACTCCTGCATTAGGAAATG | |
| sorting_P3 | TTCTCGGGTGCGACTCCTGCATTAGGAAATG | |
| sorting_P4 | GCCTACTCTGCGACTCCTGCATTAGGAAATG | |
| sorting_P5 | TAGAGAGATGCGACTCCTGCATTAGGAAATG | |
| sorting_P6 | AAACAACCTGCGACTCCTGCATTAGGAAATG | |
| sorting_P7 | CTTACAGTTGCGACTCCTGCATTAGGAAATG | Forward primers used to clone the promoter and RBS region, each with an 8-nt barcode to identify the corresponding sorting bin |
| sorting_P8 | GAAGGCATTGCGACTCCTGCATTAGGAAATG | |
| sorting_P9 | CTTGGGACTGCGACTCCTGCATTAGGAAATG | |
| sorting_P10 | ACTATGCGTGCGACTCCTGCATTAGGAAATG | |
| sorting_P11 | TCTCCAACTGCGACTCCTGCATTAGGAAATG | |
| sorting_P12 | CCGTGGTTTGCGACTCCTGCATTAGGAAATG | |
| sorting_P13 | GTAACCCGTGCGACTCCTGCATTAGGAAATG | |

| | | |
|---|---|---|
| F-yihVp3_BBa_J61106 | GGATttttgtcatttatgacatgctttgcttgtctgtttttgatcgtatttgtaatttatcgtcTCTAGAGAAAGATAGGAGACACTAGT | Oligonucleotides used in Golden Gate Assembly to construct pMPTPV_dual_yihVp3_BBa_J61106 |
| R-yihVp3_BBa_J61106 | GCATACTAGTGTCTCCTATCTTTCTCTAGAgacgataaattacaaatacgatcaaaaacagacaagcaaagcatgtcataaatgacaaaa | |
| F-zinTp_BBa_J61106 | GGATtgctctcgtttcctaagagttgttgcattttgctatatgttacaatataacattacacatTCTAGAGAAAGATAGGAGACACTAGT | Oligonucleotides used in Golden Gate Assembly to construct pMPTPV_dual_zinTp_BBa_J61106 |
| R-zinTp_BBa_J61106 | GCATACTAGTGTCTCCTATCTTTCTCTAGAatgtgtaatgttatattgtaacatatagcaaaatgcaacaactcttaggaaacgagagca | |
| F-inaAp_BBa_J61106 | GGATttcattaatacgacacgtttcattaagattttcctcaggtaaccacctatagtcattgtcTCTAGAGAAAGATAGGAGACACTAGT | Oligonucleotides used in Golden Gate Assembly to construct pMPTPV_dual_inaAp_BBa_J61106 |
| R-inaAp_BBa_J61106 | GCATACTAGTGTCTCCTATCTTTCTCTAGAgacaatgactataggtggttacctgaggaaaatcttaatgaaacgtgtcgtattaatgaa | |
| F-mngBp_BBa_J61106 | GGATcggcctttcgacacccggtgccggcattttctcgctctttttacttcatgataatggcgcTCTAGAGAAAGATAGGAGACACTAGT | Oligonucleotides used in Golden Gate Assembly to construct pMPTPV_dual_mngBp_BBa_J61106 |
| R-mngBp_BBa_J61106 | GCATACTAGTGTCTCCTATCTTTCTCTAGAgcgccattatcatgaagtaaaaagagcgagaaaatgccggcaccgggtgtcgaaaggccg | |
| F-aroLp_BBa_J61106 | GGATtaaatgtaatttattatttacacttcattcttgaatatttattggtatagtaaggggtgtTCTAGAGAAAGATAGGAGACACTAGT | Oligonucleotides used in Golden Gate Assembly to construct pMPTPV_dual_aroLp_BBa_J61106 |
| R-aroLp_BBa_J61106 | GCATACTAGTGTCTCCTATCTTTCTCTAGAacaccccttactataccaataaatattcaagaatgaagtgtaaataataaattacattta | |
| F-folEp_BBa_J61106 | GGATcgtggctccttgttgtgttgtttgcaatttcttcatcataccgtttattcaatgtgctgtTCTAGAGAAAGATAGGAGACACTAGT | Oligonucleotides used in Golden Gate Assembly to construct pMPTPV_dual_folEp_BBa_J61106 |
| R-folEp_BBa_J61106 | GCATACTAGTGTCTCCTATCTTTCTCTAGAacagcacattgaataaacggtatgatgaagaaattgcaaacaacacaacaaggagccacg | |
| F-agaSp_BBa_J61106 | GGATtcgttttatttctttttctccattgaactttcagtttcttttctatagattttaatcaaTCTAGAGAAAGATAGGAGACACTAGT | Oligonucleotides used in Golden Gate Assembly to construct pMPTPV_dual_agaSp_BBa_J61106 |
| R-agaSp_BBa_J61106 | GCATACTAGTGTCTCCTATCTTTCTCTAGAttgattaaaatctatagaaaagaaactgaaagttcaatggagaaaaaagaaataaaacga | |
| F-ybhDp2_BBa_J61106 | GGATatttttttcataataatgccctcaatttaaaaacataacttattgcgatatggttacattTCTAGAGAAAGATAGGAGACACTAGT | Oligonucleotides used in Golden Gate Assembly to construct pMPTPV_dual_ybhDp2_BBa_J61106 |
| R-ybhDp2_BBa_J61106 | GCATACTAGTGTCTCCTATCTTTCTCTAGAaatgtaaccatatcgcaataagttatgtttttaaattgagggcattattatgaaaaaaat | |

| | | |
|---|---|---|
| F-mgtAp2_BBa_J61106 | GGATatctttgatggtcagccgattttgcatcctgttgtcctgtaacgtgttgtttaattatttTCTAGAGAAAGATAGGAGACACTAGT | Oligonucleotides used in Golden Gate Assembly to construct pMPTPV_dual_mgtAp2_BBa_J61106 |
| R-mgtAp2_BBa_J61106 | GCATACTAGTGTCTCCTATCTTTCTCTAGAaaataattaaacaacacgttacaggacaacaggatgcaaaatcggctgaccatcaaagat | |
| F-cspIp2_BBa_J61106 | GGATattttattatttgtacctcttgagatttccttgttggtttttctctctgatatttttttTCTAGAGAAAGATAGGAGACACTAGT | Oligonucleotides used in Golden Gate Assembly to construct pMPTPV_dual_cspIp2_BBa_J61106 |
| R-cspIp2_BBa_J61106 | GCATACTAGTGTCTCCTATCTTTCTCTAGAaaaaaaaatatcagagagaaaaaccaacaaggaaatctcaagaggtacaaataataaaat | |
| F-hemAp2_BBa_J61106 | GGATacgttgcagttataacccttaatgctagcgttaccgtccgctatcgtctatgttcaagttTCTAGAGAAAGATAGGAGACACTAGT | Oligonucleotides used in Golden Gate Assembly to construct pMPTPV_dual_hemAp2_BBa_J61106 |
| R-hemAp2_BBa_J61106 | GCATACTAGTGTCTCCTATCTTTCTCTAGAaacttgaacatagacgatagcggacggtaacgctagcattaagggttataactgcaacgt | |
| F-yigFp5_BBa_J61106 | GGATtcaggcaatggttcgatgatttttttgatttgttatttttccactttcagtatgctaatgTCTAGAGAAAGATAGGAGACACTAGT | Oligonucleotides used in Golden Gate Assembly to construct pMPTPV_dual_yigFp5_BBa_J61106 |
| R-yigFp5_BBa_J61106 | GCATACTAGTGTCTCCTATCTTTCTCTAGAcattagcatactgaaagtggaaaaataacaaatcaaaaaaatcatcgaaccattgcctga | |
| F-mdtEp2_BBa_J61106 | GGATtttgtcgttgacctcaccatgtcgatcactgtgcctgtatcccaccttactggctgacaaTCTAGAGAAAGATAGGAGACACTAGT | Oligonucleotides used in Golden Gate Assembly to construct pMPTPV_dual_mdtEp2_BBa_J61106 |
| R-mdtEp2_BBa_J61106 | GCATACTAGTGTCTCCTATCTTTCTCTAGAttgtcagccagtaaggtgggatacaggcacagtgatcgacatggtgaggtcaacgacaaa | |
| F-nagBp_BBa_J61106 | GGATatatcgttattatcactcccttttactggctaaaccagaaaacttattttatcattcaaaTCTAGAGAAAGATAGGAGACACTAGT | Oligonucleotides used in Golden Gate Assembly to construct pMPTPV_dual_nagBp_BBa_J61106 |
| R-nagBp_BBa_J61106 | GCATACTAGTGTCTCCTATCTTTCTCTAGAtttgaatgataaaataagttttctggtttagccagtaaaagggagtgataataacgatat | |
| F-zurp6_BBa_J61106 | GGATaatagcgagttaacttgaagcaatttatttaaaaatgaattattaattattatcgtgttgTCTAGAGAAAGATAGGAGACACTAGT | Oligonucleotides used in Golden Gate Assembly to construct pMPTPV_dual_zurp6_BBa_J61106 |
| R-zurp6_BBa_J61106 | GCATACTAGTGTCTCCTATCTTTCTCTAGAcaacacgataataattaataattcatttttaaataaattgcttcaagttaactcgctatt | |
| F-croEp5_BBa_J61106 | GGATacccttgtaccgttaaggtacaagtatcttgaaggttcatttcaatcatgtaatatgtacTCTAGAGAAAGATAGGAGACACTAGT | Oligonucleotides used in Golden Gate Assembly to construct pMPTPV_dual_croEp5_BBa_J61106 |
| R-croEp5_BBa_J61106 | GCATACTAGTGTCTCCTATCTTTCTCTAGAgtacatattacatgattgaaatgaaccttcaagatacttgtaccttaacggtacaagggt | |

| | | |
|---|---|---|
| F-yegIp8_BBa_J61106 | GGATccccatatagaaataattttaatggttagttgttattattcattgaaatataatataactTCTAGAGAAAGATAGGAGACACTAGT | Oligonucleotides used in Golden Gate Assembly to construct pMPTPV_dual_yegIp8_BBa_J61106 |
| R-yegIp8_BBa_J61106 | GCATACTAGTGTCTCCTATCTTTCTCTAGAagttatattatatttcaatgaataataacaactaaccattaaaattatttctatatgggg | |
| F-ybcKp7_BBa_J61106 | GGATttatatgagcctcgttttatgctttttgttaatgtctttattttttatgtattcttttgtTCTAGAGAAAGATAGGAGACACTAGT | Oligonucleotides used in Golden Gate Assembly to construct pMPTPV_dual_ybcKp7_BBa_J61106 |
| R-ybcKp7_BBa_J61106 | GCATACTAGTGTCTCCTATCTTTCTCTAGAacaaaagaatacataaaaaataaagacattaacaaaaagcataaaacgaggctcatataa | |
| F-phoPp2_BBa_J61106 | GGATgattatccgctttttattttttcactttacctcccctccccgctggtttatttaatgtttTCTAGAGAAAGATAGGAGACACTAGT | Oligonucleotides used in Golden Gate Assembly to construct pMPTPV_dual_phoPp2_BBa_J61106 |
| R-phoPp2_BBa_J61106 | GCATACTAGTGTCTCCTATCTTTCTCTAGAaaacattaaataaaccagcggggaggggaggtaaagtgaaaaaataaaaagcggataatc | |
| F-ndkp4_BBa_J61106 | GGATtgcaatagtctgttttgtccctgattgttgctaaaactcattttattttaaaaaaatgttTCTAGAGAAAGATAGGAGACACTAGT | Oligonucleotides used in Golden Gate Assembly to construct pMPTPV_dual_ndkp4_BBa_J61106 |
| R-ndkp4_BBa_J61106 | GCATACTAGTGTCTCCTATCTTTCTCTAGAaacatttttttaaaataaaatgagttttagcaacaatcagggacaaaacagactattgca | |
| F-icdAp2_apFAB864 | GGATtctaaaagaagtttttttgcatggtattttcagagattatgaattgccgcattatagcctaATCTTAATCTAGCCCGGGAGCATTT | Oligonucleotides used in Golden Gate Assembly to construct pMPTPV_dual_icdAp2_apFAB864 |
| R-icdAp2_apFAB864 | GCATAAATGCTCCCGGGCTAGATTAAGATtaggctataatgcggcaattcataatctctgaaaataccatgcaaaaaaacttcttttaga | |
| F-fliEp1_apFAB894 | GGATcgtttttttgttgctatttagcgcctttgtcttattgacttactggtaggctttgctaccATCTTAATCTAGCGCTGGAGGCTTT | Oligonucleotides used in Golden Gate Assembly to construct pMPTPV_dual_fliEp1_apFAB894 |
| R-fliEp1_apFAB894 | GCATAAAGCCTCCAGCGCTAGATTAAGATggtagcaaagcctaccagtaagtcaataagacaaaggcgctaaatagcaacaaaaaaacg | |
| F-gatYp_apFAB872 | GGATtattgtcgtttttgtgatcgttatctcgatatttaaaaacaaataatttcattatattttATCTTAATCTAGCCCGGGATAATTT | Oligonucleotides used in Golden Gate Assembly to construct pMPTPV_dual_gatYp_apFAB872 |
| R-gatYp_apFAB872 | GCATAAATTATCCCGGGCTAGATTAAGATaaaatataatgaaattatttgtttttaaatatcgagataacgatcacaaaaacgacaata | |
| F-yibBp9_apFAB872 | GGATtccgatgtatgataaataacggaagcatttgaatggataatatacagtatacattctattATCTTAATCTAGCCCGGGATAATTT | Oligonucleotides used in Golden Gate Assembly to construct pMPTPV_dual_yibBp9_apFAB872 |
| R-yibBp9_apFAB872 | GCATAAATTATCCCGGGCTAGATTAAGATaatagaatgtatactgtatattatccattcaaatgcttccgttatttatcatacatcgga | |

| | | |
|---|---|---|
| F-fliEp1_apFAB827 | GGATcgtttttttgttgctatttagcgcctttgtcttattgacttactggtaggctttgctaccATCTTAATCTAGCAGGGGAGTCTTT | Oligonucleotides used in Golden Gate Assembly to construct pMPTPV_dual_fliEp1_apFAB827 |
| R-fliEp1_apFAB827 | GCATAAAGACTCCCCTGCTAGATTAAGATggtagcaaagcctaccagtaagtcaataagacaaaggcgctaaatagcaacaaaaaaacg | |
| F-yeaQp3_apFAB872 | GGATgatgtttgtggttggtaattatcactgatcgttgatggcgcaatttctgcgccatcgcaaATCTTAATCTAGCCCGGGATAATTT | Oligonucleotides used in Golden Gate Assembly to construct pMPTPV_dual_yeaQp3_apFAB872 |
| R-yeaQp3_apFAB872 | GCATAAATTATCCCGGGCTAGATTAAGATttgcgatggcgcagaaattgcgccatcaacgatcagtgataattaccaaccacaaacatc | |
| F-ileSp2_apFAB909 | GGATgccatctccgggcgtgtagtccacggtgatgaattagggcgcactataggtttcccgacgATCTTAATCTAGCGAAGGATAGTTT | Oligonucleotides used in Golden Gate Assembly to construct pMPTPV_dual_ileSp2_apFAB909 |
| R-ileSp2_apFAB909 | GCATAAACTATCCTTCGCTAGATTAAGATcgtcgggaaacctatagtgcgccctaattcatcaccgtggactacacgcccggagatggc | |
| F-rrsEp_apFAB827 | GGATagtcatttttctgcaatttttctattgcggcctgcggagaactccctataatgcgcctccATCTTAATCTAGCAGGGGAGTCTTT | Oligonucleotides used in Golden Gate Assembly to construct pMPTPV_dual_rrsEp_apFAB827 |
| R-rrsEp_apFAB827 | GCATAAAGACTCCCCTGCTAGATTAAGATggaggcgcattatagggagttctccgcaggccgcaatagaaaaattgcagaaaaatgact | |
| F-yihVp3_apFAB927 | GGATttttgtcatttatgacatgctttgcttgtctgtttttgatcgtatttgtaatttatcgtcATCTTAATCTAGCTGGGGACTGTTT | Oligonucleotides used in Golden Gate Assembly to construct pMPTPV_dual_yihVp3_apFAB927 |
| R-yihVp3_apFAB927 | GCATAAACAGTCCCCAGCTAGATTAAGATgacgataaattacaaatacgatcaaaaacagacaagcaaagcatgtcataaatgacaaaa | |
| F-ompNp4_apFAB914 | GGATcttattatcatttcagaggaattatttgattaaggttttacttaaggcgtaacaaatgatATCTTAATCTAGCGGCGGATCTTTT | Oligonucleotides used in Golden Gate Assembly to construct pMPTPV_dual_ompNp4_apFAB914 |
| R-ompNp4_apFAB914 | GCATAAAAGATCCGCCGCTAGATTAAGATatcatttgttacgccttaagtaaaaccttaatcaaataattcctctgaaatgataataag | |
| F-ileSp2_apFAB916 | GGATgccatctccgggcgtgtagtccacggtgatgaattagggcgcactataggtttcccgacgATCTTAATCTAGCATTGGAGGGTTT | Oligonucleotides used in Golden Gate Assembly to construct pMPTPV_dual_ileSp2_apFAB916 |
| R-ileSp2_apFAB916 | GCATAAACCCTCCAATGCTAGATTAAGATcgtcgggaaacctatagtgcgccctaattcatcaccgtggactacacgcccggagatggc | |
| F-folEp_apFAB864 | GGATcgtggctccttgttgtgttgtttgcaatttcttcatcataccgtttattcaatgtgctgtATCTTAATCTAGCCCGGGAGCATTT | Oligonucleotides used in Golden Gate Assembly to construct pMPTPV_dual_folEp_apFAB864 |
| R-folEp_apFAB864 | GCATAAATGCTCCCGGGCTAGATTAAGATacagcacattgaataaacggtatgatgaagaaattgcaaacaacacaacaaggagccacg | |

| | | |
|---|---|---|
| F-fliEp1_apFAB909 | GGATcgttttttttgttgctatttagcgcctttgtcttattgacttactggtaggctttgctaccATCTTAATCTAGCGAAGGATAGTTT | Oligonucleotides used in Golden Gate Assembly to construct pMPTPV_dual_fliEp1_apFAB909 |
| R-fliEp1_apFAB909 | GCATAAACTATCCTTCGCTAGATTAAGATggtagcaaagcctaccagtaagtcaataagacaaaggcgctaaatagcaacaaaaaaaacg | |
| F-folEp_apFAB894 | GGATcgtggctccttgttgtgttgtttgcaatttcttcatcataccgtttattcaatgtgctgtATCTTAATCTAGCGCTGGAGGCTTT | Oligonucleotides used in Golden Gate Assembly to construct pMPTPV_dual_folEp_apFAB894 |
| R-folEp_apFAB894 | GCATAAAGCCTCCAGCGCTAGATTAAGATacagcacattgaataaacggtatgatgaagaaattgcaaacaacacaacaaggagccacg | |
| F-yebGp_apFAB827 | GGATgtctttattgctgatgttgatttctcaaccgaaaagaaatatactgtataaaatcacagtATCTTAATCTAGCAGGGGAGTCTTT | Oligonucleotides used in Golden Gate Assembly to construct pMPTPV_dual_yebGp_apFAB827 |
| R-yebGp_apFAB827 | GCATAAAGACTCCCCTGCTAGATTAAGATactgtgattttatacagtatatttcttttcggttgagaaatcaacatcagcaataaagac | |
| F-folEp_apFAB927 | GGATcgtggctccttgttgtgttgtttgcaatttcttcatcataccgtttattcaatgtgctgtATCTTAATCTAGCTGGGGACTGTTT | Oligonucleotides used in Golden Gate Assembly to construct pMPTPV_dual_folEp_apFAB927 |
| R-folEp_apFAB927 | GCATAAACAGTCCCCAGCTAGATTAAGATacagcacattgaataaacggtatgatgaagaaattgcaaacaacacaacaaggagccacg | |
| F-yihVp3_apFAB864 | GGATttttgtcatttatgacatgctttgcttgtctgtttttgatcgtatttgtaatttatcgtcATCTTAATCTAGCCCGGGAGCATTT | Oligonucleotides used in Golden Gate Assembly to construct pMPTPV_dual_yihVp3_apFAB864 |
| R-yihVp3_apFAB864 | GCATAAATGCTCCCGGGCTAGATTAAGATgacgataaattacaaatacgatcaaaaacagacaagcaaagcatgtcataaatgacaaaa | |
| F-rrsDp1_apFAB914 | GGATcagaaaaaaagatcaaaaaaatacttgtgcaaaaaattgggatccctataatgcgcctccATCTTAATCTAGCGGCGGATCTTTT | Oligonucleotides used in Golden Gate Assembly to construct pMPTPV_dual_rrsDp1_apFAB914 |
| R-rrsDp1_apFAB914 | GCATAAAAGATCCGCCGCTAGATTAAGATggaggcgcattatagggatcccaatttttgcacaagtatttttttgatctttttttctg | |
| F-fliEp1_apFAB927 | GGATcgttttttttgttgctatttagcgcctttgtcttattgacttactggtaggctttgctaccATCTTAATCTAGCTGGGGACTGTTT | Oligonucleotides used in Golden Gate Assembly to construct pMPTPV_dual_fliEp1_apFAB927 |
| R-fliEp1_apFAB927 | GCATAAACAGTCCCCAGCTAGATTAAGATggtagcaaagcctaccagtaagtcaataagacaaaggcgctaaatagcaacaaaaaaaacg | |
| F-aroLp_apFAB914 | GGATtaaatgtaatttattatttacacttcattcttgaatatttattggtatagtaaggggtgtATCTTAATCTAGCGGCGGATCTTTT | Oligonucleotides used in Golden Gate Assembly to construct pMPTPV_dual_aroLp_apFAB914 |
| R-aroLp_apFAB914 | GCATAAAAGATCCGCCGCTAGATTAAGATacaccccttactataccaataaatattcaagaatgaagtgtaaataataaattacattta | |

| | | |
|---|---|---|
| F-yihVp3_apFAB894 | GGATttttgtcatttatgacatgctttgcttgtctgtttttgatcgtatttgtaatttatcgtcATCTTAATCTAGCGCTGGAGGCTTT | Oligonucleotides used in Golden Gate Assembly to construct pMPTPV_dual_yihVp3_apFAB894 |
| R-yihVp3_apFAB894 | GCATAAAGCCTCCAGCGCTAGATTAAGATgacgataaattacaaatacgatcaaaaacagacaagcaaagcatgtcataaatgacaaaa | |
| F-fliEp1_apFAB914 | GGATcgtttttttgttgctatttagcgcctttgtcttattgacttactggtaggctttgctaccATCTTAATCTAGCGGCGGATCTTTT | Oligonucleotides used in Golden Gate Assembly to construct pMPTPV_dual_fliEp1_apFAB914 |
| R-fliEp1_apFAB914 | GCATAAAAGATCCGCCGCTAGATTAAGATggtagcaaagcctaccagtaagtcaataagacaaaggcgctaaatagcaacaaaaaaacg | |
| F-yfbKp3_apFAB914 | GGATgggttgcgttgttgaatcattatttgctatgggtttcgatattgtattttattaagattaATCTTAATCTAGCGGCGGATCTTTT | Oligonucleotides used in Golden Gate Assembly to construct pMPTPV_dual_yfbKp3_apFAB914 |
| R-yfbKp3_apFAB914 | GCATAAAAGATCCGCCGCTAGATTAAGATtaatcttaataaaatacaatatcgaaacccatagcaaataatgattcaacaacgcaaccc | |
| F-yfjHp2_apFAB864 | GGATtagatggtggcccgtgccatacctgtttttggataaccgctggtaaccgtagtattctaATCTTAATCTAGCCCGGGAGCATTT | Oligonucleotides used in Golden Gate Assembly to construct pMPTPV_dual_yfjHp2_apFAB864 |
| R-yfjHp2_apFAB864 | GCATAAATGCTCCCGGGCTAGATTAAGATtagaatactacggttaccagcggttatccaaaaaacaggtatggcacgggccaccatcta | |
| F-yihVp3_apFAB827 | GGATttttgtcatttatgacatgctttgcttgtctgtttttgatcgtatttgtaatttatcgtcATCTTAATCTAGCAGGGGAGTCTTT | Oligonucleotides used in Golden Gate Assembly to construct pMPTPV_dual_yihVp3_apFAB827 |
| R-yihVp3_apFAB827 | GCATAAAGACTCCCCTGCTAGATTAAGATgacgataaattacaaatacgatcaaaaacagacaagcaaagcatgtcataaatgacaaaa | |
| F-J23103 | GGATctgatagctagctcagtcctagggattatgctagcTCTAGAGAAAGATAGGAGACACTAGT | Oligonucleotides used in Golden Gate Assembly to construct pMPTPV_dual_J23103_BBa_J61106 |
| R-J23103 | GCATACTAGTGTCTCCTATCTTTCTCTAGAgctagcataatccctaggactgagctagctatcag | |
| F-J23109 | GGATtttacagctagctcagtcctagggactgtgctagcTCTAGAGAAAGATAGGAGACACTAGT | Oligonucleotides used in Golden Gate Assembly to construct pMPTPV_dual_J23109_BBa_J61106 |
| R-J23109 | GCATACTAGTGTCTCCTATCTTTCTCTAGAgctagcacagtccctaggactgagctagctgtaaa | |
| F-J23115 | GGATtttatagctagctcagcccttggtacaatgctagcTCTAGAGAAAGATAGGAGACACTAGT | Oligonucleotides used in Golden Gate Assembly to construct pMPTPV_dual_J23115_BBa_J61106 |
| R-J23115 | GCATACTAGTGTCTCCTATCTTTCTCTAGAgctagcattgtaccaagggctgagctagctataaa | |

| | | |
|---|---|---|
| F-J23107 | GGATtttacggctagctcagccctaggtattatgctagcTCTAGAGAAAGATAGGAGACACTAGT | Oligonucleotides used in Golden Gate Assembly to construct pMPTPV_dual_J23107_BBa_J61106 |
| R-J23107 | GCATACTAGTGTCTCCTATCTTTCTCTAGAgctagcataatacctagggctgagctagccgtaaa | |
| F-J23101 | GGATtttacagctagctcagtcctaggtattatgctagcTCTAGAGAAAGATAGGAGACACTAGT | Oligonucleotides used in Golden Gate Assembly to construct pMPTPV_dual_J23101_BBa_J61106 |
| R-J23101 | GCATACTAGTGTCTCCTATCTTTCTCTAGAgctagcataatacctaggactgagctagctgtaaa | |
| F-J23103_J23103 | GGATctgatagctagctgatagctagcgattatctagggattatgctagcTCTAGAGAAAGATAGGAGACACTAGT | Oligonucleotides used in Golden Gate Assembly to construct pMPTPV_dual_J23103_J23103_BBa_J61106 |
| R-J23103_J23103 | GCATACTAGTGTCTCCTATCTTTCTCTAGAgctagcataatccctagataatcgctagctatcagctagctatcag | |
| F-J23103_J23109 | GGATctgatagctagtttacagctagcgattatctagggactgtgctagcTCTAGAGAAAGATAGGAGACACTAGT | Oligonucleotides used in Golden Gate Assembly to construct pMPTPV_dual_J23103_J23109_BBa_J61106 |
| R-J23103_J23109 | GCATACTAGTGTCTCCTATCTTTCTCTAGAgctagcacagtccctagataatcgctagctgtaaactagctatcag | |
| F-J23103_J23115 | GGATctgatagctagtttatagctagcgattatcttggtacaatgctagcTCTAGAGAAAGATAGGAGACACTAGT | Oligonucleotides used in Golden Gate Assembly to construct pMPTPV_dual_J23103_J23115_BBa_J61106 |
| R-J23103_J23115 | GCATACTAGTGTCTCCTATCTTTCTCTAGAgctagcattgtaccaagataatcgctagctataaactagctatcag | |
| F-J23103_J23107 | GGATctgatagctagtttacggctagcgattatctaggtattatgctagcTCTAGAGAAAGATAGGAGACACTAGT | Oligonucleotides used in Golden Gate Assembly to construct pMPTPV_dual_J23103_J23107_BBa_J61106 |
| R-J23103_J23107 | GCATACTAGTGTCTCCTATCTTTCTCTAGAgctagcataatacctagataatcgctagccgtaaactagctatcag | |
| F-J23103_J23101 | GGATctgatagctagtttacagctagcgattatctaggtattatgctagcTCTAGAGAAAGATAGGAGACACTAGT | Oligonucleotides used in Golden Gate Assembly to construct pMPTPV_dual_J23103_J23101_BBa_J61106 |
| R-J23103_J23101 | GCATACTAGTGTCTCCTATCTTTCTCTAGAgctagcataatacctagataatcgctagctgtaaactagctatcag | |
| F-J23109_J23103 | GGATtttacagctagctgatagctagcgactgtctagggattatgctagcTCTAGAGAAAGATAGGAGACACTAGT | Oligonucleotides used in Golden Gate Assembly to construct pMPTPV_dual_J23109_J23103_BBa_J61106 |
| R-J23109_J23103 | GCATACTAGTGTCTCCTATCTTTCTCTAGAgctagcataatccctagacagtcgctagctatcagctagctgtaaa | |

| | | |
|---|---|---|
| F-J23109_J23109 | GGATtttacagctagtttacagctagcgactgtctagggactgtgctagcTCTAGAGAAAGATAGGAGACACTAGT | Oligonucleotides used in Golden Gate Assembly to construct pMPTPV_dual_J23109_J23109_BBa_J61106 |
| R-J23109_J23109 | GCATACTAGTGTCTCCTATCTTTCTCTAGAgctagcacagtccctagacagtcgctagctgtaaactagctgtaaa | |
| F-J23109_J23115 | GGATtttacagctagtttatagctagcgactgtcttggtacaatgctagcTCTAGAGAAAGATAGGAGACACTAGT | Oligonucleotides used in Golden Gate Assembly to construct pMPTPV_dual_J23109_J23115_BBa_J61106 |
| R-J23109_J23115 | GCATACTAGTGTCTCCTATCTTTCTCTAGAgctagcattgtaccaagacagtcgctagctataaactagctgtaaa | |
| F-J23109_J23107 | GGATtttacagctagtttacggctagcgactgtctaggtattatgctagcTCTAGAGAAAGATAGGAGACACTAGT | Oligonucleotides used in Golden Gate Assembly to construct pMPTPV_dual_J23109_J23107_BBa_J61106 |
| R-J23109_J23107 | GCATACTAGTGTCTCCTATCTTTCTCTAGAgctagcataatacctagacagtcgctagccgtaaactagctgtaaa | |
| F-J23109_J23101 | GGATtttacagctagtttacagctagcgactgtctaggtattatgctagcTCTAGAGAAAGATAGGAGACACTAGT | Oligonucleotides used in Golden Gate Assembly to construct pMPTPV_dual_J23109_J23101_BBa_J61106 |
| R-J23109_J23101 | GCATACTAGTGTCTCCTATCTTTCTCTAGAgctagcataatacctagacagtcgctagctgtaaactagctgtaaa | |
| F-J23115_J23103 | GGATtttatagctagctgatagctagctacaatctagggattatgctagcTCTAGAGAAAGATAGGAGACACTAGT | Oligonucleotides used in Golden Gate Assembly to construct pMPTPV_dual_J23115_J23103_BBa_J61106 |
| R-J23115_J23103 | GCATACTAGTGTCTCCTATCTTTCTCTAGAgctagcataatccctagattgtagctagctatcagctagctataaa | |
| F-J23115_J23109 | GGATtttatagctagtttacagctagctacaatctagggactgtgctagcTCTAGAGAAAGATAGGAGACACTAGT | Oligonucleotides used in Golden Gate Assembly to construct pMPTPV_dual_J23115_J23109_BBa_J61106 |
| R-J23115_J23109 | GCATACTAGTGTCTCCTATCTTTCTCTAGAgctagcacagtccctagattgtagctagctgtaaactagctataaa | |
| F-J23115_J23115 | GGATtttatagctagtttatagctagctacaatcttggtacaatgctagcTCTAGAGAAAGATAGGAGACACTAGT | Oligonucleotides used in Golden Gate Assembly to construct pMPTPV_dual_J23115_J23115_BBa_J61106 |
| R-J23115_J23115 | GCATACTAGTGTCTCCTATCTTTCTCTAGAgctagcattgtaccaagattgtagctagctataaactagctataaa | |
| F-J23115_J23107 | GGATtttatagctagtttacggctagctacaatctaggtattatgctagcTCTAGAGAAAGATAGGAGACACTAGT | Oligonucleotides used in Golden Gate Assembly to construct pMPTPV_dual_J23115_J23107_BBa_J61106 |
| R-J23115_J23107 | GCATACTAGTGTCTCCTATCTTTCTCTAGAgctagcataatacctagattgtagctagccgtaaactagctataaa | |

| | | |
|---|---|---|
| F-J23115_J23101 | GGATtttatagctagtttacagctagctacaatctaggtattatgctagcTCTAGAGAAAGATAGGAGACACTAGT | Oligonucleotides used in Golden Gate Assembly to construct pMPTPV_dual_J23115_J23101_BBa_J61106 |
| R-J23115_J23101 | GCATACTAGTGTCTCCTATCTTTCTCTAGAgctagcataatacctagattgtagctagctgtaaactagctataaa | |
| F-J23107_J23103 | GGATtttacggctagctgatagctagctattatctagggattatgctagcTCTAGAGAAAGATAGGAGACACTAGT | Oligonucleotides used in Golden Gate Assembly to construct pMPTPV_dual_J23107_J23103_BBa_J61106 |
| R-J23107_J23103 | GCATACTAGTGTCTCCTATCTTTCTCTAGAgctagcataatccctagataatagctagctatcagctagccgtaaa | |
| F-J23107_J23109 | GGATtttacggctagtttacagctagctattatctagggactgtgctagcTCTAGAGAAAGATAGGAGACACTAGT | Oligonucleotides used in Golden Gate Assembly to construct pMPTPV_dual_J23107_J23109_BBa_J61106 |
| R-J23107_J23109 | GCATACTAGTGTCTCCTATCTTTCTCTAGAgctagcacagtccctagataatagctagctgtaaactagccgtaaa | |
| F-J23107_J23115 | GGATtttacggctagtttatagctagctattatcttggtacaatgctagcTCTAGAGAAAGATAGGAGACACTAGT | Oligonucleotides used in Golden Gate Assembly to construct pMPTPV_dual_J23107_J23115_BBa_J61106 |
| R-J23107_J23115 | GCATACTAGTGTCTCCTATCTTTCTCTAGAgctagcattgtaccaagataatagctagctataaactagccgtaaa | |
| F-J23107_J23107 | GGATtttacggctagtttacggctagctattatctaggtattatgctagcTCTAGAGAAAGATAGGAGACACTAGT | Oligonucleotides used in Golden Gate Assembly to construct pMPTPV_dual_J23107_J23107_BBa_J61106 |
| R-J23107_J23107 | GCATACTAGTGTCTCCTATCTTTCTCTAGAgctagcataatacctagataatagctagccgtaaactagccgtaaa | |
| F-J23107_J23101 | GGATtttacggctagtttacagctagctattatctaggtattatgctagcTCTAGAGAAAGATAGGAGACACTAGT | Oligonucleotides used in Golden Gate Assembly to construct pMPTPV_dual_J23107_J23101_BBa_J61106 |
| R-J23107_J23101 | GCATACTAGTGTCTCCTATCTTTCTCTAGAgctagcataatacctagataatagctagctgtaaactagccgtaaa | |
| F-J23101_J23103 | GGATtttacagctagctgatagctagctattatctagggattatgctagcTCTAGAGAAAGATAGGAGACACTAGT | Oligonucleotides used in Golden Gate Assembly to construct pMPTPV_dual_J23101_J23103_BBa_J61106 |
| R-J23101_J23103 | GCATACTAGTGTCTCCTATCTTTCTCTAGAgctagcataatccctagataatagctagctatcagctagctgtaaa | |
| F-J23101_J23109 | GGATtttacagctagtttacagctagctattatctagggactgtgctagcTCTAGAGAAAGATAGGAGACACTAGT | Oligonucleotides used in Golden Gate Assembly to construct pMPTPV_dual_J23101_J23109_BBa_J61106 |
| R-J23101_J23109 | GCATACTAGTGTCTCCTATCTTTCTCTAGAgctagcacagtccctagataatagctagctgtaaactagctgtaaa | |

| | | |
|---|---|---|
| F-J23101_J23115 | GGATtttacagctagtttatagctagctattatcttggtacaatgctagcTCTAGAGAAAGATAGGAGACACTAGT | Oligonucleotides used in Golden Gate Assembly to construct pMPTPV_dual_J23101_J23115_BBa_J61106 |
| R-J23101_J23115 | GCATACTAGTGTCTCCTATCTTTCTCTAGAgctagcattgtaccaagataatagctagctataaactagctgtaaa | |
| F-J23101_J23107 | GGATtttacagctagtttacggctagctattatctaggtattatgctagcTCTAGAGAAAGATAGGAGACACTAGT | Oligonucleotides used in Golden Gate Assembly to construct pMPTPV_dual_J23101_J23107_BBa_J61106 |
| R-J23101_J23107 | GCATACTAGTGTCTCCTATCTTTCTCTAGAgctagcataatacctagataatagctagccgtaaactagctgtaaa | |
| F-J23101_J23101 | GGATtttacagctagtttacagctagctattatctaggtattatgctagcTCTAGAGAAAGATAGGAGACACTAGT | Oligonucleotides used in Golden Gate Assembly to construct pMPTPV_dual_J23101_J23101_BBa_J61106 |
| R-J23101_J23101 | GCATACTAGTGTCTCCTATCTTTCTCTAGAgctagcataatacctagataatagctagctgtaaactagctgtaaa | |
| F-control_J23103 | GGATgcaggcgctagctgatagctagctcagtcctagggattatgctagcTCTAGAGAAAGATAGGAGACACTAGT | Oligonucleotides used in Golden Gate Assembly to construct pMPTPV_dual_control_J23103_BBa_J61106 |
| R-control_J23103 | GCATACTAGTGTCTCCTATCTTTCTCTAGAgctagcataatccctaggactgagctagctatcagctagcgcctgc | |
| F-control_J23109 | GGATgcaggcgctagtttacagctagctcagtcctagggactgtgctagcTCTAGAGAAAGATAGGAGACACTAGT | Oligonucleotides used in Golden Gate Assembly to construct pMPTPV_dual_control_J23109_BBa_J61106 |
| R-control_J23109 | GCATACTAGTGTCTCCTATCTTTCTCTAGAgctagcacagtccctaggactgagctagctgtaaactagcgcctgc | |
| F-control_J23115 | GGATgcaggcgctagtttatagctagctcagcccttggtacaatgctagcTCTAGAGAAAGATAGGAGACACTAGT | Oligonucleotides used in Golden Gate Assembly to construct pMPTPV_dual_control_J23115_BBa_J61106 |
| R-control_J23115 | GCATACTAGTGTCTCCTATCTTTCTCTAGAgctagcattgtaccaagggctgagctagctataaactagcgcctgc | |
| F-control_J23107 | GGATgcaggcgctagtttacggctagctcagccctaggtattatgctagcTCTAGAGAAAGATAGGAGACACTAGT | Oligonucleotides used in Golden Gate Assembly to construct pMPTPV_dual_control_J23107_BBa_J61106 |
| R-control_J23107 | GCATACTAGTGTCTCCTATCTTTCTCTAGAgctagcataatacctagggctgagctagccgtaaactagcgcctgc | |
| F-control_J23101 | GGATgcaggcgctagtttacagctagctcagtcctaggtattatgctagcTCTAGAGAAAGATAGGAGACACTAGT | Oligonucleotides used in Golden Gate Assembly to construct pMPTPV_dual_control_J23101_BBa_J61106 |
| R-control_J23101 | GCATACTAGTGTCTCCTATCTTTCTCTAGAgctagcataatacctaggactgagctagctgtaaactagcgcctgc | |

1

**Supplementary Table 3** Annealing formulation for constructing malonyl-CoA biosensors

| Plasmids | Oligo 1 | Oligo 2 | Oligo 3 | Oligo 4 | Oligo 5 |
|---|---|---|---|---|---|
| pCYC1-OP_TATA-UAS_DDC | oligo_25 | oligo_13 | oligo_20 | oligo_2 | oligo_19 |
| pCYC1-OP_TATA-UAS_EBC | oligo_14 | oligo_13 | oligo_20 | oligo_2 | oligo_10 |
| pHSP12-TATA_OP-UAS_BDC | oligo_9 | oligo_1 | oligo_22 | oligo_7 | oligo_24 |
| pCYC1-TATA_OP-UAS_EAC | oligo_4 | oligo_1 | oligo_22 | oligo_7 | oligo_11 |
| pCYC1-OP_TATA-UAS_BDC | oligo_6 | oligo_13 | oligo_20 | oligo_2 | oligo_17 |
| pCYC1-OP_TATA-UAS_FDC | oligo_18 | oligo_13 | oligo_20 | oligo_2 | oligo_23 |
| pEXG1-N30_OP-UAS_FDA | oligo_12 | oligo_16 | oligo_15 | oligo_3 | oligo_21 |
| pCYC1-OP_TATA-UAS_BEC | oligo_5 | oligo_13 | oligo_20 | oligo_2 | oligo_8 |

**Supplementary Table 4** Bin boundaries for all experiments

| | TnaC library | Malonyl-CoA library | Promoter library (rep1) | Promoter library (rep2) | Promoter library (rep3) | Combi-nation library (rep1) | Combi-nation library (rep2) | Combi-nation library (rep3) |
|---|---|---|---|---|---|---|---|---|
| **criteria** | $\log_{10}\frac{eGFP}{mCherry^{0.88098}}$* | $\log_{10}\frac{YPeT}{mCherry}$ | | $\log_{10}\frac{sfGFP}{mCherry}$ | | | $\log_{10}\frac{sfGFP}{mCherry}$ | |
| $\boldsymbol{b_0}$ | $-\infty$ | $-\infty$ | $-\infty$ | $-\infty$ | $-\infty$ | $-\infty$ | $-\infty$ | $-\infty$ |
| $\boldsymbol{b_1}$ | -0.48532 | -1.21148 | -1.14861 | -1.13581 | -1.14666 | -1.46654 | -1.46634 | -1.47661 |
| $\boldsymbol{b_2}$ | -0.09034 | -0.89106 | -0.86981 | -0.85744 | -0.87144 | -1.20980 | -1.21212 | -1.22398 |
| $\boldsymbol{b_3}$ | 0.29797 | -0.57793 | -0.70194 | -0.69102 | -0.70726 | -1.07125 | -1.07399 | -1.09029 |
| $\boldsymbol{b_4}$ | 0.69046 | -0.26665 | -0.57487 | -0.56612 | -0.58463 | -0.92695 | -0.92050 | -0.93965 |
| $\boldsymbol{b_5}$ | 1.09311 | 0.04661 | -0.47450 | -0.46784 | -0.48712 | -0.77107 | -0.76021 | -0.77859 |
| $\boldsymbol{b_6}$ | $+\infty$ | 0.35926 | -0.39001 | -0.38446 | -0.40488 | -0.61380 | -0.60255 | -0.62044 |
| $\boldsymbol{b_7}$ | - | 0.67608 | -0.31311 | -0.30829 | -0.32938 | -0.47166 | -0.46175 | -0.47971 |
| $\boldsymbol{b_8}$ | - | $+\infty$ | -0.23930 | -0.23449 | -0.25595 | -0.33505 | -0.32531 | -0.34466 |
| $\boldsymbol{b_9}$ | - | - | -0.16321 | -0.15781 | -0.17959 | -0.18268 | -0.17315 | -0.19295 |
| $\boldsymbol{b_{10}}$ | - | - | -0.07552 | -0.06968 | -0.09112 | -0.00661 | 0.00293 | -0.01845 |
| $\boldsymbol{b_{11}}$ | - | - | 0.05066 | 0.05769 | 0.03624 | 0.21727 | 0.22719 | 0.20235 |
| $\boldsymbol{b_{12}}$ | - | - | $+\infty$ | $+\infty$ | $+\infty$ | $+\infty$ | $+\infty$ | $+\infty$ |

* 0.88098 is the slope of the boundary lines on the log-log plot.

**Supplementary Table 5** The ratio of cells for each bin in one sorting experiment

| | **Promoter library (rep1)** | **Promoter library (rep2)** | **Promoter library (rep3)** | **Combination library (rep1)** | **Combination library (rep2)** | **Combination library (rep3)** |
|---|---|---|---|---|---|---|
| $bin_1$ | 0.08333 | 0.08333 | 0.08333 | 0.08333 | 0.08333 | 0.08333 |
| $bin_2$ | 0.08333 | 0.08333 | 0.08333 | 0.08333 | 0.08333 | 0.08333 |
| $bin_3$ | 0.08333 | 0.08333 | 0.08333 | 0.08333 | 0.08333 | 0.08333 |
| $bin_4$ | 0.08333 | 0.08333 | 0.08333 | 0.08333 | 0.08333 | 0.08333 |
| $bin_5$ | 0.08333 | 0.08333 | 0.08333 | 0.08333 | 0.08333 | 0.08333 |
| $bin_6$ | 0.08333 | 0.08333 | 0.08333 | 0.08333 | 0.08333 | 0.08333 |
| $bin_7$ | 0.08333 | 0.08333 | 0.08333 | 0.08333 | 0.08333 | 0.08333 |
| $bin_8$ | 0.08333 | 0.08333 | 0.08333 | 0.08333 | 0.08333 | 0.08333 |
| $bin_9$ | 0.08333 | 0.08333 | 0.08333 | 0.08333 | 0.08333 | 0.08333 |
| $bin_{10}$ | 0.08333 | 0.08333 | 0.08333 | 0.08333 | 0.08333 | 0.08333 |
| $bin_{11}$ | 0.08333 | 0.08333 | 0.08333 | 0.08333 | 0.08333 | 0.08333 |
| $bin_{12}$ | 0.08333 | 0.08333 | 0.08333 | 0.08333 | 0.08333 | 0.08333 |

**Supplementary Table 6** Slopes and intercepts of the linear regressions shown in **Supplementary Figure 28**

| RBS | Slope | Intercept |
|---|---|---|
| apFAB872 | 0.0334±0.0102 | 0.0050±0.0013 |
| apFAB914 | 0.0322±0.0043 | 0.0042±0.0008 |
| apFAB864 | 0.0254±0.0023 | 0.0045±0.0007 |
| apFAB865 | 0.0199±0.0009 | 0.0051±0.0005 |
| apFAB927 | 0.0228±0.0015 | 0.0052±0.0008 |
| apFAB827 | 0.0313±0.0018 | 0.0015±0.0013 |
| apFAB894 | 0.0246±0.0014 | 0.0055±0.0010 |
| apFAB909 | 0.0238±0.0015 | 0.0036±0.0010 |
| apFAB839 | 0.0233±0.0013 | 0.0037±0.0010 |
| apFAB833 | 0.0233±0.0015 | 0.0044±0.0012 |
| apFAB834 | 0.0225±0.0013 | 0.0038±0.0011 |
| apFAB820 | 0.0281±0.0020 | 0.0026±0.0018 |
| apFAB916 | 0.0247±0.0020 | 0.0040±0.0019 |

**Captions of Supplementary Data**

**Supplementary Data 1** The expression characteristics of each *tanC* variant at different ligand concentrations (calculated with dSort-Seq).

**Supplementary Data 2** The responses of each malonyl-CoA biosensor at different ligand concentrations (calculated with dSort-Seq).

**Supplementary Data 3** The expression characteristics of each *E. coli* endogenous promoter (calculated with dSort-Seq).

**Supplementary Data 4** The expression characteristics of each combination in the combination library (calculated with dSort-Seq).

**Supplementary Data 5** The regulatory information of endogenous promoters in *E. coli* K12 MG1655 (obtained from EcoCyC).

**Statistical information and software used in this work**

Plots were generated in Python 3.8 using the matplotlib (3.4.3) plotting libraries. All statistical analyses, data fitting, interpolation calculations and machine learning were performed using the SciPy (1.9.1), NumPy (1.23.4), scikit-learn (1.1.2), TensorFlow (2.10.0) and TensorFlow Probability (0.18.0) Python packages.
